# Nrf1 is an indispensable redox-determining factor for mitochondrial homeostasis by integrating multi-hierarchical regulatory networks

**DOI:** 10.1101/2022.05.04.490622

**Authors:** Shaofan Hu, Jing Feng, Meng Wang, Reziyamu Wufuer, Keli Liu, Zhengwen Zhang, Yiguo Zhang

## Abstract

To defend a vast variety of challenges in the oxygenated environments, all life forms have been evolutionally established a set of antioxidant, detoxification and cytoprotective systems during natural selection and adaptive survival, in order to maintain cell redox homeostasis and organ integrity in the healthy development and growth. Such antioxidant defense systems are predominantly regulated by two key transcription factors Nrf1 and Nrf2, but the underlying mechanism(s) for their coordinated redox control remains elusive. Here, we found that loss of full-length Nrf1 led to a dramatic increase in reactive oxygen species (ROS) and oxidative damages in *Nrf1α^-/-^* cells, and this increase was not eliminated by drastic elevation of Nrf2, even though the antioxidant systems were also substantially enhanced by hyperactive Nrf2. Further studies revealed that the increased ROS production in *Nrf1α^-/-^* resulted from a striking impairment in the mitochondrial oxidative respiratory chain and its gene expression regulated by nuclear respiratory factors, called αPal^NRF1^ and GABP^NRF2^. In addition to antioxidant capacity of cells, glycolysis was greatly augmented by aberrantly-elevated Nrf2, so to partially relieve the cellular energy demands, but aggravate its mitochondrial stress. The generation of ROS was also differentially regulated by Nrf1 and Nrf2 through miR-195 and/or mIR-497-mediated UCP2 pathway. Consequently, the epithelial-mesenchymal transformation (EMT) of *Nrf1α^-/-^* cells was activated by putative ROS-stimulated signaling *via* MAPK, HIF1α, NF-kB, PI3K and AKT, all players involved in cancer development and progression. Taken together, it is inferable that Nrf1 acts as a potent integrator of redox regulation by multi-hierarchical networks.

## INTRODUCTION

Under the oxygenated environments, almost all cellular life forms have been successively generating, transforming and eliminating those reactive oxygen species (ROS, also including reactive nitrogen and sulphur species) in a variety of cell processes (e.g., metabolism, proliferation, differentiation), immune regulation and vascular remodeling (1). The ROS primarily encompass superoxide anion (O_2_·^-^), hydrogen peroxide (H_2_O_2_) and hydroxyl radical (·OH^-^), with a certain +

er chemical reactivity towards lipids, proteins, and nucleic acids (RNA and DNA), so that they are endowed with a cytotoxic capability to beget oxidative damage, and even lead to cell death if the excessive ROS are yielded. Of note, approximately 90% of ROS in eukaryotic cells are generated from the mitochondria, which are hence accepted as a major source of the intracellularly-producing ROS. Clearly, approximately 2% of oxygens are consumed by mitochondria to produce ROS, and this number could be incremented when their electron transport chains (ETCs) are damaged and/or uncoupled with the oxidative phosphorylation (OXPHOS, for ATP production)(2). During the normal aerobic respiration, the ETC complexes can relay electrons from nicotinamide adenine dinucleotide (NADH) and dihydroflavin adenine dinucleotide (FADH2), respectively, to Complex IV, whereupon oxygen accepts those transferred electrons to become water. In this process, some electrons can enable, however, to escape from the ETC reactions to convert O_2_ into O_2_·^-^ Of and other ROS in order (3). Besides, NADPH oxidases, cytochrome p450 isoenzymes and the other oxidases existing particularly in the peroxisome and endoplasmic reticulum (ER) are considered as the secondary sources of ROS production(4).

In order to defend the potential challenges from oxidative stress, all life forms have also evolutionally established a set of antioxidant, detoxification and cytoprotective systems to maintain cellular redox homeostasis and organ integrity during their adaptive survival. Amongst them, the most direct antioxidant enzymes include superoxide dismutases [SODs, which convert O_2_·^-^ to less reactive H_2_O_2_], catalase [CAT, which continually coverts H_2_O_2_ to H_2_O and O_2_], glutathione peroxidases(GPXs) and peroxiredoxins (PRDXs); both can reduce H_2_O_2_ to H_2_O, by using the reducing power derived from glutathione or thioredoxins (TXNs)(5). Besides, the other important defensive mechanisms and mediators of redox signaling are represented by the thioredoxins and glutathione/glutaredoxin systems. Moreover, there exist many indirectly-acting antioxidant defense systems. For instance, heme oxygenase isoenzymes (i.e., HO-1 and HO-2) eliminate free heme and prevent it from yielding strong free radicals (e.g., ·OH^-^) during oxidative stress (6). Consequently, such a homeodynamic balance between oxidative stress and antioxidant defense system has been established and also perpetuated at a steady state of redox, just at which all living organisms are allowed for healthy survival under such a robust redox homeostasis.

Collectively, the levels of ROS (albeit with higher cytotoxicity) under the normal conditions are finely tuned and also restricted to considerably lower extents, such that they can also serve as a potent eustress that triggers certain hormestic effects on the physiology of living organisms insomuch as to preserve the healthy life processes (7,8). Thereby, a redox threshold is set at a certain steady-state of physiology. Once ROS are produced to exceed the preformed redox threshold, they exert as oxidative distress to stimulate a vast variety of pathophysiological responses and even pathological effects to different extents (9). The proper ROS levels are required to induce many biological activities, including development, growth, cell division, proliferation, differentiation, survival and apoptosis, as well as the immune responses (1,4). Rather, excessive ROS have manifested with some pathogenic relevance to many of human diseases, such as neurodegenerative, cardiovascular, inflammatory and autoimmune diseases, diabetes, aging and cancer (10). Of note, ROS have an capability to promote tumourgenesis by inducing the genomic instability and its DNA mutation of key genes or by activating those critical molecules involved in some signaling pathways leading to carcinogenesis (11). This is based on the facts that ROS can promote cancer development and progression in many aspects, including cellular proliferation induced by activation of the MEK-ERK1/2 signaling pathway (12), evasion from cell apoptosis or anoikis after activation of either NF-κB or PI3K-AKT signaling pathways (13,14), migration and invasion triggered by EMT (15), along with tumor angiogenesis stimulated by releasing vascular endothelial growth factor (VEGF) and angiopoietin (16). These, together with many other signaling pathways towards cognate gene regulatory networks mediated by Nrf1 (nuclear factor erythroid 2-related factor 1, also called Nfe2l1^Nrf1^), Nrf2 (nuclear factor erythroid 2-related factor 2, also called Nfe2l2^Nrf2^), AP-1, c-myc and p53, in addition to PTEN, PKC, Ras, Raf, JNK and/or p38 kinase(17), are stimulated or over-stimulated, which depends on their responses to varying extents of oxidative stress caused by ROS (and other reactive oxidants) (9,18).

In mammalians, Nrf1 and Nrf2 are two principal regulators of the intracellular redox homeostasis by coordinately governing transcriptional expression of distinct subsets of cognate genes through their functional heterodimers with a partner of small Maf (sMaf) or other bZIP proteins (e.g., AP-1 and ATF4), that are recruited for directly binding to those consensus sites, called antioxidant and/or electrophile response elements (AREs/EpREs), within their target promoter regions (19,20). Clearly, both Nrf1 and Nrf2 are differentially activated in distinctive tempo-spatial responses to different extents of ROS-led oxidative stress (Figure 1A). Under normal conditions, Nrf1 is anchored to the ER and most degraded *via* the ubiquitin-mediated proteasome pathway. When Nrf1 is required for biological responses to stimulation, this CNC-bZIP protein is allowed for dynamic dislocation from the ER lumen across membranes into extra-ER compartments, in which it is subjected to its selective proteolytic cleavage by DDI1/2 or other cytosolic proteases so as to become a mature transcription factor, before entering the nucleus and transactivating target genes(21,22). By contrast, most of Nrf2 is inactive under normal conditions, because it is segregated in the cytoplasm by physic interaction with Keap1, a redox-sensitive adaptor for Cullin3, so as to target this CNC-bZIP protein to the ubiquitin-mediated proteasomal degradation (23). Once Keep is activated by increased ROS, Nrf2 is allowed for disassociation from its interactor Keap1 to enter the nucleus, in which it enables to heterodimerize with sMaf and transactivate target genes.

**Figure1.**
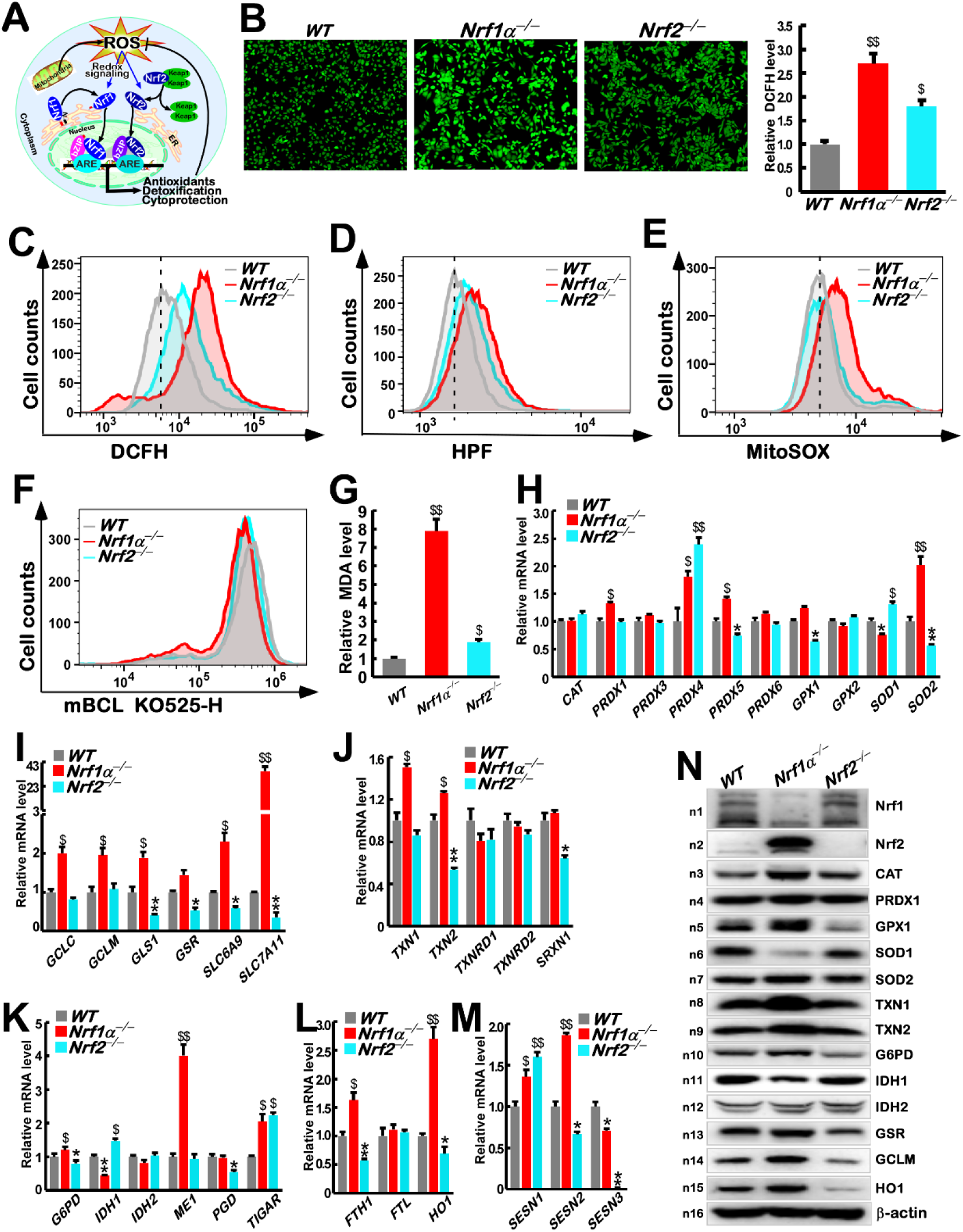
The Redox status and expression of antioxidant genes in *WT, Nrf1α^-/-^, Nrf2^-/-^* cells. (A) A schematic representation of signaling mechanisms that Nrf1 and Nrf2 are activated by ROS to play cytoprotective roles. (B) Fluorescence images representing the ROS levels in *WT*, *Nrf1α^-/-^* and *Nrf2^-/-^* cells, which were stained by DCFH-DA (excited in 488nm). The fluorescence intensity was statistically calculated as fold changes (mean ± SEM, n= 3 × 3, $, p < 0.05 and $$, p < 0.01). (C, D, E) Distinctive ROS levels were detected by flow cytometry in *WT, Nrf1α^-/-^* and *Nrf2^-/-^* cell lines that had been stained by DCFH-DA (C), HPF (D) or MitoSOX (E), respectively. (F) Different GSH levels were determined by flow cytometry in *WT*, *Nrf1α^-/-^* and *Nrf2^-/-^* cell lines that had all been stained by monochlorobimane (mBCI. Ex/Em=394/490nm). (G) Relative malondialdehyde (MDA) levels were measured in *WT*, *Nrf1α^-/-^* and Nrf2 cells. The data are shown as fold changes (mean ± SEM, n= 3 × 3; $, p < 0.05 and $$, p < 0.01). (H) The mRNA levels of those antioxidant genes *CAT, PRDX1, PRDX3, PRDX4, PRDX5, PRDX6, GPX1, GPX2, SOD1*, and *SOD2* were determined by RT-qPCR in *WT*, *Nrf1α^-/-^* and *Nrf2^-/-^* cells. The data are shown as mean ± SEM (n= 3 × 3; $, p < 0.05 and $$, p < 0.01; *, p < 0.05 and **, p < 0.01). (I) The mRNA levels of GSH metabolism genes *GCLC, GCLM, GLS1, GSR, SLC6A9* and *SLA7C11* were examined by RT-qPCR in *WT*, *Nrf1α^-/-^* and *Nrf2^-/-^* cells. The data are shown as mean ± SEM (n= 3 × 3; $, p < 0.05 and $$, p < 0.01; *, p < 0.05 and **, p < 0.01). (J) The mRNA levels of *TXN1, TXN2, TXNRD1, TXNRD2* and *SRXN1* determined by RT-qPCR in *WT*, *Nrf1α^-/-^* and *Nrf2^-/-^* cells are shown as mean ± SEM (n= 3 × 3; $, p < 0.05 and $$, p < 0.01; *, p < 0.05 and **, p < 0.01). (K) The mRNA levels of *G6PD, IDH1, IDH2, ME1, PGD* and *TIGAR* determined by RT-qPCR in *WT*, *Nrf1α^-/-^* and *Nrf2^-/-^* cells are shown as mean ± SEM (n= 3 × 3; $, p < 0.05 and $$, p < 0.01; *, p < 0.05 and **, p < 0.01). (L) The mRNA levels of *FTH1, FTL and HO1* determined by RT-qPCR in *WT*, *Nrf1α^-/-^* and *Nrf2^-/-^* cells are shown as mean ± SEM (n= 3 × 3; $, p < 0.05 and $$, p < 0.01; *, p < 0.05 and **, p < 0.01). (M) The mRNA levels of *SESN1, SESN2* and *SESN3* were determined by qPCR in *WT*, *Nrf1α^-/-^* and *Nrf2^-/-^* cells. The resulting data are shown as mean ± SEM (n= 3 × 3; $, p < 0.05 and $$, p < 0.01; *, p < 0.05 and **, p < 0.01). (N) Distinct protein levels of Nrf1, Nrf2, CAT, PRDX1, GPX1, SOD1, SOD2, TXN1, TXN2, G6PD, IDH1, IDH2, GSR, GCLM, and HO1 in *WT, Nrf1α^-/-^* and *Nrf2^-/-^* cells were visualized by Western blotting. The intensity of all immunoblots was also calculated as shown in Figure S3.

Those target genes mediated by Nrf1 and/or Nrf2 are mainly involved in: i) glutathione production and regeneration, which are regulated by glutamate–cysteine ligase modifier (GCLM) and catalytic subunits (GCLC), glutathione reductase (GSR)(24); ii) glutathione utilization, which is conducted by glutathione S-transferases (GSTs) and glutathione peroxidase 2 (GPXs); iii) thioredoxin production and utilization (e.g., TXNs, PRDXs); iv) NADPH production, controlled by glucose-6-phosphate dehydrogenase (G6PD), malic enzyme 1 (ME1) and isocitrate dehydrogenase 1 (IDH1) (25); and v) the other responsible genes for antioxidant and detoxification, e.g., both encoding NAD(P)H:quinone oxidoreductase 1 (NQO1) (26) and haem oxygenase 1 (HO-1)(6). Furthermore, Nrf1 acts as a vital player in maintaining protein homeostasis (proteostasis) by controlling transcriptional expression of genes encoding all proteasomal subunits and relevant co-factors (27). Nrf1 is essential for lipid metabolism by regulating PGC1β, Lipin1, PPARγ and CD36 (28–30), as well as for glucose metabolism by governing expression of HK1, Glut2, GCK, Sl2a2, Fbp1 and PCK1 (31,32). Moreover, Nrf1 is also indispensable for cell differentiation, embryonic development, and cytoprotection against inflammatory transformation into cancer (20). Of note, Nrf2 can also exert a protective effect on normal cells against chemical carcinogens, but rather promotes cancer development and progression, particularly when it is hyper-activated for a long term (33). By contrast, Nrf1α (and long isoform TCF11) is identified as a potent tumor-repressor to inhibit cancer malgrowth in xenograft model mice (34). This is supported by liver-specific knockout of Nrf1 in mice with obvious pathological phenotypes, resembling human non-alcoholic steatohepatitis (NASH) and hepatoma (35,36). Our previous evidence also revealed that knockout of Nrf1 from human hepatoma cells results in the exacerbation of tumor growth and migration, but such malignant growth appears to be strikingly prevented upon overexpression of Nrf1α or TCF11 (34,37).

In this study, we found that knockout of Nrf1α caused a substantial increase of the intracellular ROS determined in *Nrf1α^-/-^* cells, when compared with wild-type (*WT*) and *Nrf2^-/-^* cells. Such increased ROS levels were accompanied by up-regulation of most genes responsible for antioxidant and detoxification; this is attributable to aberrant accumulation of Nrf2 upon loss of Nrf1α. Interestingly, abnormal changes in the mitochondrial morphology of *Nrf1α^-/-^* rather than *Nrf2^-/-^* cells were observed, as accompanied by impaired ETC complexes. Further examinations also unraveled that *Nrf1α^-/-^*-derived mitochondrial dysfunction was owing to deregulation of two nuclear respiratory factors αPal^NRF1^ and GABPα^NRF2^, along with the co-factor PGC1α, resulting in strikingly increased ROS levels in *Nrf1α*-deficient cells. Besides, glycolysis of *Nrf1α^-/-^* cells was greatly augmented by hyperactive Nrf2, leading to an aggravated pressure of mitochondrial stress. The mitochondrial UCP2 was markedly suppressed by Nrf2 through AREs-driven miR-195 and miR-497. These collectively contribute to a drastic increase of ROS in *Nrf1α^-/-^* cells; this is concomitant with activation of multiple signaling pathways *via* MAPK, PI3K-AKT, HIF1α and NF-κB to the EMT process involved in cancer development and progression. Overall, Nrf1 acts as a vital player in determining and maintaining robust redox homeostasis by governing mitochondrial homeostasis integrated from multi-hierarchical signaling pathways towards cognate gene regulatory networks. Therefore, this work provides a novel understanding of distinct roles for Nrf1 and Nrf2 in cancer development and prevention.

## RESULTS

### A dramatic increase of ROS results from loss of Nrf1α, but is not eliminated by hyperactive Nrf2 in *Nrf1α^-/-^* cells

Albeit it was previously reported that the intracellular ROS levels were increased in mouse *Nrf1-* and *Nrf2*-deficient cells (38), but the underlying mechanism remains elusive, to date. Contrarily, Nrf2 was also shown to amplify oxidative stress by induction of Kruppel-like factor 9 (KLF9) in response to elevated ROS over presetting threshold (39). To gain an insight into distinct roles of Nrf1 and Nrf2 in determining and maintaining redox homeostasis, we reexamined human *Nrf1α^-/-^* and *Nrf2^-/-^* cell lines; both were established by the genomic editing of HepG2 cells (34,37). Here, we found that knockout of either *Nrf1α^-/-^* or *Nrf2^-/-^* caused an obvious increase in their intracellular ROS levels (detected by DCFH-DA, as the most commonly used probe with a relative large range from detecting H_2_O_2_, O_2_·^-^,·OH^-^ to ONOO^-^), but the increased ROS in *Nrf1α^-/-^* cells were significantly higher than those in *Nrf2^-/-^* cells (Figure 1B, C). To confirm this result, two additional fluorescent probes HPF (Hydroxyphenyl fluorescein, sensing particularly to intracellular O_2_·^-^ and ONOO^-^ changes) and MitoSOX (specifically sensing to mitochondrial O_2_·^-^ changes) were also employed (Figure S1B). The resulting data revealed that distinct extents of the increased ROS in between *Nrf1α^-/-^* and *Nrf2^-/-^* cell lines were amplified by HPF (Figure 1D). However, it is, to our surprise, found that almost no changes in the mitochondrial O_2_·^-^-sensing fluorescence were detected by MitoSOX in *Nrf2^-/-^* cells, but *Nrf1α^-/-^* cells were manifested with a significant magnitude of its mitochondrial ROS (Figure 1E), when compared to *WT* controls. A concomitant decrease in the reduced glutathione (GSH, as a free radical scavenger and potent detoxifying agent) was also determined in *Nrf1α^-/-^* and *Nrf2^-/-^* cell lines (Figure 1F); this is negatively correlated with the intracellular ROS levels, as described (40). As such, further examination of malondialdehyde (MDA, a marker of polyunsaturated fatty acids peroxidation in the cell) unraveled a substantial increased production in *Nrf1α^-/-^* cells, to 8 times higher than that obtained from *WT* cells, whereas *Nrf2^-/-^* cells only displayed ~2-fold changes (Figure 1G). Collectively, these demonstrate that loss of *Nrf1α* has led to severe endogenous oxidative distress caused primarily by increased ROS from mitochondria and lipid peroxidation, but, by contrast, loss of *Nrf2* only causes a modest oxidative stress.

To explore the reasons underlying such distinct increases of ROS by deficiency of *Nrf1α* from *Nrf2*, it was surprisingly unveiled by transcriptome sequencing that most of those differential expression genes responsible for antioxidant and detoxification were significantly up-regulated in *Nrf1α^-/-^* cells, but also markedly down-regulated in *Nrf2^-/-^* cells, when compared to *WT* cells (Figure S2). Further quantitative PCR of critical genes for ROS elimination revealed that *Nrf1α*^-/-^cells gave rise to obvious increases in basal expression of *SOD2* (superoxide dismutase 2, converting the mitochondrial O_2_·^-^into H_2_O_2_), *PRDX1* (peroxiredoxin 1)*, PRDX4, PRDX5* (all three scavenging H_2_O_2_ by consuming reduced thioredoxin) and *GPX1* (glutathione peroxidase 1), but extra-mitochondrial *SOD1* expression was decreased (Figure 1H). Conversely, *Nrf2^-/-^* manifested significantly decreased expression of *SOD2, PRDX5* and *GPX1* in, but *PRDX4* was highly expressed, while *SOD1* was modestly increased. Also, it was, much to our surprise, found that basal expression of those examined genes *GCLC, GCLM, GLS, GSR, SLC6A9* and particularly *SLC7A11* (all involved in GSH biosynthesis and regeneration) was significantly up-regulated in *Nrf1α*^-/-^cells, but down-regulated in *Nrf2^-/-^* cells, when compared with *WT* controls (Figure 1I). Additional two reducing powers thioredoxin-1 (*TXN1*) and −2 (*TXN2*) were obviously up-expressed in *Nrf1α^-/-^* cells, but down-expressed in *Nrf2^-/-^* cells, whereas basal expression of thioredoxin reductase-1 (*TXNRD1*) and −2 (*TXNRD2*) was almost not changed in both cell lines (Figure 1J). Moreover, basal expression of sulfiredoxin 1 (*SRXN1*) was reduced only in *Nrf2^-/-^* cells, but largely unaltered in *Nrf1α^-/-^* cells (Figure 1J). Collectively, these findings demonstrate that loss of *Nrf2* results in obvious defects in *de novo* biosynthesis of GSH and TXN and their recycling, but rather unusual increases of them have strikingly emerged upon loss of *Nrf1α*.

Further examination revealed that basal expression of *G6PD* (a key enzyme in the pentose phosphate pathway to yield NADPH) and 6-phosphogluconate dehydrogenase (*PGD*, as the second dehydrogenase in the pentose phosphate shunt) was slightly increased or unaltered in *Nrf1α^-/-^* cells respectively, but apparently decreased in *Nrf2^-/-^* cells (Figure 1K). Conversely, significant increases in basal *TIGAR* (TP53 induced glycolysis regulatory phosphatase, blocking glycolysis and directing the pathway into the pentose phosphate shunt to protects cells from DNA-damaging ROS) expression were observed in both *Nrf1α^-/-^* and *Nrf2^-/-^* cell lines (Figure 1K). Besides, basal expression of malic enzymes (*ME1*, as another NADPH supplier) was also substantially up-regulated in *Nrf1α^-/-^* cells to ~4-fold changes higher than its *WT* control, but almost unchanged in *Nrf2^-/-^* cells. By contrast, isocitrate dehydrogenase 1 (*IDH1*, a key enzyme catalyzing the oxidative decarboxylation of isocitrate to alpha-ketoglutarate to generate CO2 and NADPH in the citric acid cycle), rather than IDH2, was significantly lowered in *Nrf1α^-/-^* cells, but highly expressed in *Nrf2^-/-^* cells (Figure 1K). Altogether, these indicate knockout of *Nrf1α* leads to a redox metabolism reprogramming, that is distinctive from knockout of *Nrf2*.

Next, by quantitative investigation of another antioxidant capability to prevent free heme from forming free radicals during oxidative stress (6), it was unraveled that basal expression levels of *HO-1* and *FTH1* (ferritin heavy chain 1) were evidently up-regulated in *Nrf1α^-/-^* cells, but down-regulated in *Nrf2^-/-^* cells (Figure 1L), while no changes in the expression of *FTL* (ferritin light chain) were observed in both cell lines, when compared to *WT* controls. In addition, another potent antioxidant family of highly conserved sestrin (SESN, involved in the reduction of PRDXs, so to negatively regulate mTORC signaling pathways was also investigated herein(41). Among them, *SESN1* was up-expressed in both *Nrf1α^-/-^* and *Nrf2^-/-^* cell lines, whereas *SESN2* was only up-expressed in *Nrf1α^-/-^* cells, but rather down-expressed in *Nrf2^-/-^* cells (Figure 1M). By sharp contrast, basal expression of *SESN3* was significantly down-regulated in *Nrf1α^-/-^* cells and also almost abolished in *Nrf2^-/-^* cells when compared with *WT* controls. These imply distinct roles for Nrf1α and Nrf2 in governing antioxidant potentials and also redox signaling to mTORC-regulated networks so as to meet the needs of cell survival.

Further insights into the protein expression of critical genes by Western blotting revealed that, albeit the processed Nrf1-C/D isoform is only slightly decreased in *Nrf2^-/-^* cells, basal Nrf2 protein was aberrantly, substantially accumulated in *Nrf1α^-/-^* cells (Figure 1N, S3), as consistent with our previous results (37,42). Such hyperactive Nrf2 should be interpreted as a compensation for loss of Nrf1, and consequently attributable to the constant up-regulation of most antioxidant and detoxifying genes in *Nrf1α^-/-^* cells. As anticipated, most of examined proteins including CAT, PRDX1, GPX1, SOD2, TXN1, TXN2, G6PD, GSR, GCLM and HO1 were highly expressed as accompanied by hyperactive Nrf2 accumulation in *Nrf1α^-/-^* cells (Figure 1N, S3). However, basal abundances of SOD1 and IDH1, rather than IDH2, were down-regulated in *Nrf1α^-/-^* cells, but almost unaffected in *Nrf2^-/-^* cells, implying they may serve as Nrf1-specific targets.

### Loss of Nrf1α results in mitochondrial dysfunction and oxidative damage in *Nrf1α^-/-^* cells

Intriguingly, the above-described data indicate that, even though hyperactive Nrf2 accumulated with up-regulation of antioxidant and detoxification genes, *Nrf1α^-/-^* cells are still manifested with severe endogenous oxidative stress, caused primarily by ROS yielded from its mitochondria, but rather not a similar distress has been suffered from *Nrf2^-/-^* cells. To address this, we herein examine distinct subunits of mitochondrial ETC complexes I to IV in *Nrf1α^-/-^* cells, by comparison with *Nrf2^-/-^* cells and *WT* controls. As anticipated, most subunits of the mitochondrial respiratory chain were, to different extents, down-regulated in *Nrf1α^-/-^* cells (Figure 2A). By contrast, their mRNA expression in *Nrf2^-/-^* cells appeared to be unaffected or slightly enhanced, with an exception of *SDHA* (succinate dehydrogenase complex flavoprotein subunit A).

**Figure2.**
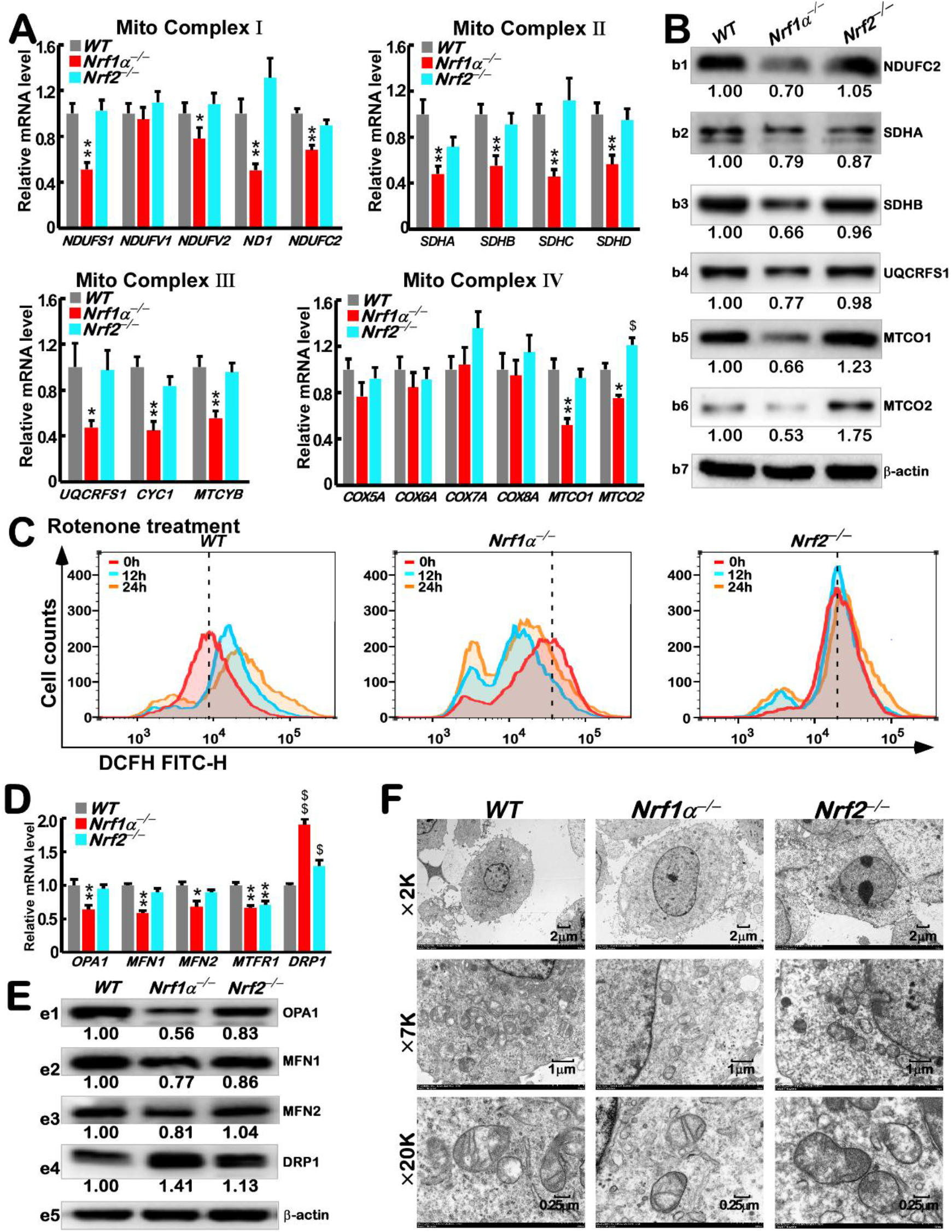
Expression of mitochondrial related genes and mitochondrial morphology in *WT, Nrf1α^-/-^*, *Nrf2^-/-^* cells. (A) The mRNA levels of *NDUFS1, NDUFV1, NDUFV2, ND1, NDUFC2, SDHA, SDHB, SDHC, SDHD, UQCRFS1, CYC1, MTCYB, COX5A, COX6A, COX7A, COX8A, MTCO1* and *MTCO2* were determined by RT-qPCR in *WT*, *Nrf1α^-/-^* and *Nrf2^-/-^* cells. The resulting data are shown as mean ± SEM (n= 3 × 3; *, p < 0.05 and **, p < 0.01). (B) The protein expression levels of NDUFC2, SDHA, SDHB, NQCRFS1, MTCO1 and MTCO2 in *WT, Nrf1α^-/-^* and *Nrf2^-/-^* cells were visualized by Western blotting. The intensity of all the immunoblots was calculated and shown on the bottom. (C) Distinct ROS levels and detected by flow cytometry in *WT*, *Nrf1α^-/-^* and *Nrf2^-/-^* cells that had been treated by Rotenone for 0 h, 12 h, or 24 h and then stained by DCFH for 30 minutes. (D) Differential mRNA expression levels of *Drp1, OPA1, MFN1, MFN2* and *MTFR1* determined by RT-qPCR in *WT*, *Nrf1α^-/-^* and *Nrf2^-/-^* cells are shown as mean ± SEM (n= 3 × 3; $, p < 0.05 and $$, p < 0.01; *, p < 0.05 and **, p < 0.01). (E) The protein levels of OPA1, MFN1, MFN2 and Drp1 in *WT*, *Nrf1α^-/-^* and *Nrf2^-/-^* cells were visualized by Western blotting. The intensity of all the immunoblots was calculated and shown on the bottom. (F) The electronic micrographs representative of *WT*, *Nrf1α^-/-^* and *Nrf2^-/-^* cells. The scale bar = 2 μm in ×2K pictures, or = 1 μm in ×7K pictures, or = 0.25 μm in ×20K pictures.

Of note, *ND1*, *MTCYB, MTCO1* and *MTCO2* are encoded by the mitochondrial genome *per se*, all other examined subunits are encoded by the nuclearly-located genes. Western blotting also revealed that protein abundances of NDUFC2, SDHA, SDHB, NQCRFS1, MTCO1, and MTCO2 were apparently decreased in *Nrf1α^-/-^*, but not *Nrf2^-/-^*, cell lines (Figure 2B). These demonstrate that loss of *Nrf1α*, rather than *Nrf2*, leads to a certain dysfunction of mitochondrial respiratory chain.

Next, treatment of *WT* cells with rotenone (a specific inhibitor of mitochondrial ETC complex I (43)) unraveled that the intracellular oxidative stress was aggravated by incrementing the mitochondrial ROS production in a time-dependent manner (Figure 2C, *left panel*), albeit Nrf2 protein expression was enhanced by this chemical (Figure S4A). Rather, similar rotenone treatment of *Nrf1α^-/-^* cells rendered its severe endogenous oxidative stress to be significantly mitigated to a certain extent (Figure 2C, *middle panel*), but roughly no obvious changes was determined after treatment of *Nrf2^-/-^* cells (Figure 2C, *right panel*). Also, no significant differences in rotenone-induced apoptosis were observed in these three cell lines (Figure S4B,C). These indicate that *Nrf1α^-/-^*-derived dysfunction of its mitochondrial respiratory chain to give rise to considerable higher ROS levels is not further worsened, but conversely relieved by rotenone, while not a similar event appears to take place in *Nrf2^-/-^* cells.

Electron micrographic imaging showed that morphological changes of *Nrf1α^-/-^* cells were distinctive from *Nrf2^-/-^* cells (Figure 2D). The mitochondria of *Nrf1α^-/-^* cells seem to be shrunk, with significant decreases in its mitochondrial number and its cristae in each mitochondrion (*middle panels*), whereas the mitochondria of *Nrf2^-/-^* cells appeared to be modestly reinforced (*right panels*), when compared with *WT* cells (*left panels*). Further examination of critical players in the mitochondrial fission and fusion (for maintaining a steady state of mitochondrial function under cellular metabolic or environmental stress (44))by RT-PCR and Western blotting revealed that *MFN1* (mitofusin 1) and *OPA1* (optic atrophy 1, serving mitochondrial dynamin-like GTPase) were obviously down-regulated in *Nrf1α^-/-^* cells (Figure 2E,F). By contrast, *DRP1* (dynamin related protein 1, a key regulator of mitochondrial division) was substantially up-regulated in *Nrf1α^-/-^* cells, even though *MTFR1* (mitochondrial fission regulator 1) was decreased. However, slightly changes in these players were examined in *Nrf2^-/-^* cells. Altogether, loss of *Nrf1α* results in dysfunctional mitochondria, thereby producing considerable higher levels of ROS along with oxidative damages.

### Nrf1 directly regulates two nuclear respiratory factors αPal^NRF1^ and GABPα^NRF2^, and co-factor PGC1α

To gain an insight into the underlying mechanism for mitochondrial dysfunction caused by loss of *Nrf1α^-/-^*, we here examined the constitutive expression of two nuclear respiratory factors αPal^NRF1^ and GABPα^NRF2^(45,46), cofactor PGC1α, and critical target genes [responsible for mitochondrial DNA replication and transcription, ETC/OXPHOS gene expression, and mitochondrial biogenesis (47)]. The results showed that basal mRNA expression levels of *αPal^NRF1^, GABPα^NRF2^* and co-targeting mitochondrial transcription factors *TFAM, TFB1M* and *TFB2M*, together with *PGC1α*, but not *PGC1β*, were obviously down-regulated, though to different extents, in *Nrf1α^-/-^* cells (Figure 3A). However, such key factors *αPal^NRF1^* and *GABPα^NRF2^* were not significantly affected in *Nrf2^-/-^* cells, which only gave rise to marginally decreased expression of *PGC1α, TFAM, TFB1M*, but not *TFB2M*. Further examinations also revealed that basal abundances of αPal^NRF1^, GABPα^NRF2^, PGC1α and TFAM proteins were down-regulated in *Nrf1α^-/-^* cells, while in *Nrf2^-/-^* cells only TFAM was down-expressed (Figure 3B, S5A). Collectively, these indicate that *Nrf1α^-/-^*-caused mitochondrial dysfunction is attributable to deregulation of nuclear respiratory controls towards the mitochondrial gene transcriptional networks. This is also further supported by restoration of ectopic Nrf1 factor into *Nrf1α^-/-^* cells (Figure S5B), showing greater or lesser extents of recovery of those four key factors (αPal^NRF1^, GABPα^NRF2^, PGC1α and TFAM), together with other mitochondrial proteins NDUFC2, NQCRFS1, MTCO1, MTCO2, and SDHB, but not SDHA. Additional supportive evidence was obtained from the tetracycline-inducible Nrf1-expressing cell lines (Figure S5C,D), in which almost all those examined mRNAs and proteins were up-regulated. By contrast, Nrf2 induction by tetracycline led to significantly increased expression of PGC1α, GABPα^NRF2^ and TFAM, besides HO-1 and GCLM. This implies at least a partial involvement of Nrf2 in regulating the mitochondrial function, particularly when it is required for certain inducible cues.

**Figure3.**
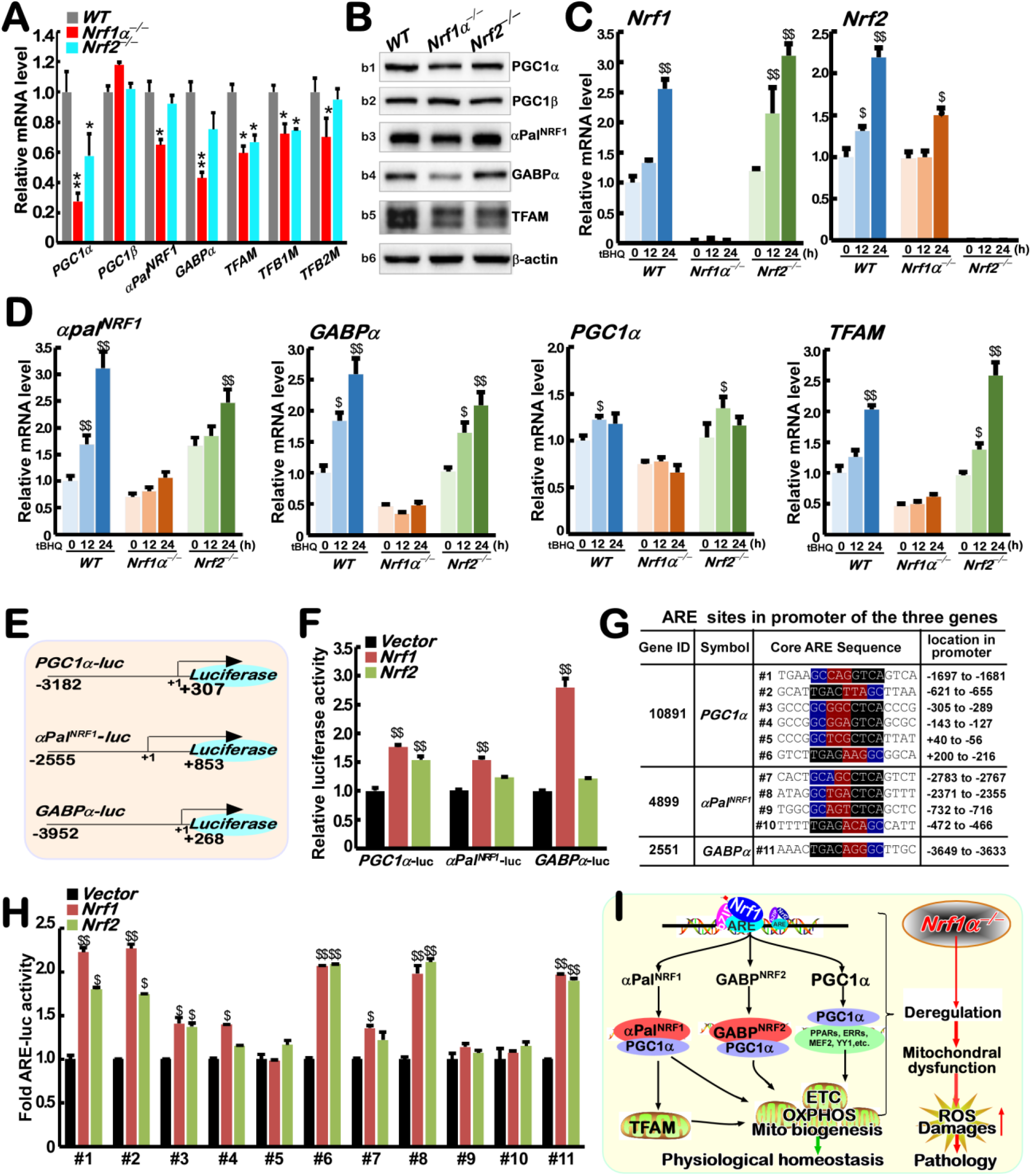
Regulating effect of Nrf1/2 on PGC1α, PGC1β, αPal^NRF1^, GABPα. (A) The mRNA expression levels of *PGC1α, PGC1β, αPal^NRF1^, GABPα, TFAM, TFB1M* and *TFB2M* were determined by RT-qPCR in *WT*, *Nrf1α^-/-^* and *Nrf2^-/-^* cells. The data are shown as fold changes (mean ± SEM, n= 3 × 3; *, p < 0.05 and **, p < 0.01). (B) The protein levels of PGC1α, PGC1β, αPal^NRF1^, GABPα and TFAM in *WT*, *Nf1α^-/-^* and *Nrf2^-/-^* cells were visualized by Western blotting. The intensity of all the immunoblots was calculated and shown in Figure S5A. (C) The mRNA levels of *Nrf1* and *Nrf2* were determined by RT-qPCR in *WT*, *Nrf1α^-/-^* and *Nrf2^-/-^* cells, which had been treated with tBHQ for 0 h, 12 h or 24 h. The data are shown as mean ± SEM (n= 3 × 3) with significant increases ($, p < 0.05 and $$, p < 0.01) as compared to the cells without this chemical treatment. (D) The mRNA levels of *αpal^NRF1^, GABPα, PGC1α*, and *TFAM* were determined by RT-qPCR in *WT*, *Nrf1α^-/-^* and *Nrf2^-/-^* cells, which had been treated with tBHQ for 0 h, 12 h or 24 h. The resulting data are shown as above. (E) Schematic representation of the *PGC1α-luc, αPal^NRF1^-luc* and *GABPα-luc*, which were constructed into the PGL3-Promoter plasmid. Their promoter regions were also indicated. (F) Relative luciferase activity of *PGC1α-luc, αPal^NRF1^-luc*, and *GABPα-luc* were determined in COS-1 cells that had been co-expressed with each reporter gene and pRL-TK (as an internal reference), plus an expression construct for Nrf1 or Nrf2, or empty pcDNA3 vector. The data are shown as mean ± SEM (n= 3 × 3; $, p < 0.05 and $$, p < 0.01). (G) The consensus ARE sites within the promoter of *PGC1α, αPal^NRF1^*, and *GABPα* were listed herein. (H) Fold *ARE-luc* activity mediated by Nrf1 or Nrf2 was determined. The indicated *ART*-adjoining sequences were cloned into PGL3-promoter vector, and were co-expressed with pRL-TK, plus an expression construct for Nrf1 or Nrf2, or empty pcDNA3 plasmid, then operated as above. (I) A proposed model that Nrf1 and Nrf2 regulates mitochondrial function by *αpal^NRF1^, GABPα* and *PGC1α*.

To further investigate distinct roles for Nrf1 and Nrf2 in the nuclear-to-mitochondrial regulation, the experimental *Nrf1α^-/-^, Nrf2^-/-^* and *WT* cell lines were treated with *tert*-butylhydroquinone [*t*BHQ, as a pro-oxidative stressor (48)]. The results revealed that *Nrf1, Nrf2, PGC1α, αPal^NRF1^, GABPα^NRF2^*, and *TFAM* (Figure 3C, D, S6A, B) along with *HO-1* and *GCLM* (Figure S6C) were significantly induced by *t*BHQ stimulation in *WT* cells. Such induction of *PGC1α, αPal^NRF1^, GABPα^NRF2^* and *TFAM* (controlling the mitochondrial function) was almost completely abolished in *Nrf1α^-/-^* cells (Figure 3C), even albeit hyperactive Nrf2 and its targeting HO-1 and GCLM were further enhanced by *t*BHQ (Figure S6A-C). Conversely, although induction of HO-1 and GCLM by *t*BHQ was abolished in *Nrf2^-/-^* cells, different inducible extents of *αPal^NRF1^, GABPα^NRF2^, TFAM* and *PGC1α* were still stimulated by this chemical, and also accompanied by induction of Nrf1 in *Nrf2*-deficient cells. Taken altogether, these demonstrate that Nrf1 rather than Nrf2 is essential for maintaining the mitochondrial functional homeostasis by governing the nuclear respiratory factors.

To clarify whether Nrf1 or Nrf2 directly regulates such key genes *αPal^NRF1^, GABPα^NRF2^* and *PGC1α*, their promoter regions were constructed into relevant luciferase reporters (as shown in Figure 3E). The results revealed that these reporter genes *αPal^NRF1^-Luc, GABPα^NRF2^-Luc* and *PGC1α-Luc* were transcriptionally activated by ectopic expression of Nrf1 (Figure 3F). Of note, transactivation activity of *GABPα^NRF2^-Luc* mediated by Nrf1 was substantially increased to ~2.5 times over its control values. By contrast, Nrf2 only gave rise to marginal transactivation of these reporter genes at considerably lower levels. Further sequence analysis unraveled that several *ARE* consensus sites are present in the promoter regions of *αPal^NRF1^, GABPα^NRF2^* and *PGC1α*, respectively (Figure 3G). Some of such *ARE*-driven luciferase reporter genes were also *trans*-activated by Nrf1 and/or Nrf2 (Figure 3H). Overall, these results indicate that Nrf1 alone or in combination with Nrf2 directly regulates *αPal^NRF1^, GABPα^NRF2^* and *PGC1α* for determining the mitochondrial homeostasis (Figure 3I). But, conversely loss of *Nrf1α^-/-^* leads to a fatal defect in these critical gene expression, resulting in mitochondrial dysfunction and oxidative damages.

### Glycolysis of *Nrf1α^-/-^* cells is enhanced so to aggravate its mitochondrial pressure with increased ROS production

Since as by-products of mitochondrial oxidative metabolism, ROS are allowed to connect with controls of metabolic process, such that the yield of ROS in this organelle depends on distinct types of fuel loading (carbohydrate, lipid, protein), besides mitochondrial functional homeostasis *per se* (49). Here, transcriptomic sequencing analysis revealed significant differential expression of those genes responsible for energy metabolism, as well as carbohydrate and lipid metabolisms of *Nrf1α^-/-^* cells (Figure S7), but these metabolic gene changes were strikingly diminished, abolished or even reversed in *Nrf2^-/-^* cells. Thereafter, *Nrf1α^-/-^, Nrf2^-/-^* and *WT* cell lines had been allowed for growth in a complete medium containing 25 mM *versus* 5 mM glucose for 24 h, before the putative effect of glucose, as a fuel load, on the amount of intracellular ROS production was determined. As shown in Figure 4A, the ROS levels detected by flow cytometry of *WT* cells fed with 25 mM glucose were augmented when compared with those arising from 5 mM glucose conditions. Such a supply of 25mM glucose deteriorated endogenous oxidative stress in *Nrf1α^-/-^* cells (*middle panel*), whereas *Nrf2^-/-^* cells appeared to be unaffected (*right panel*). Conversely, glycolytic inhibition of by 2-deoxy-D-glucose (2-DG in 25 mM glucose media) led to distinct extents of decreased ROS yield in *Nrf1α^-/-^* or *Nrf2^-/-^* cell lines (Figure S8A), but apparent increase of ROS in *WT* cells occurred after 2-DG treatment. However, treatment of *Nrf1α^-/-^* cells with insulin (to activate glycolysis) caused a modest increase of ROS, while no changes in ROS yielded from insulin-treated *WT or Nrf2^-/-^* cell lines were determined (Figure S8B, C). Collectively, these data suggest that glycolysis contributes to ROS production especially in *Nrf1α^-/-^* cells, and Nrf2 is required for the yield of ROS by feeding 25mM glucose to oxidative metabolism (e.g., glycolysis), albeit Nrf1 may also be involved in this process.

**Figure4.**
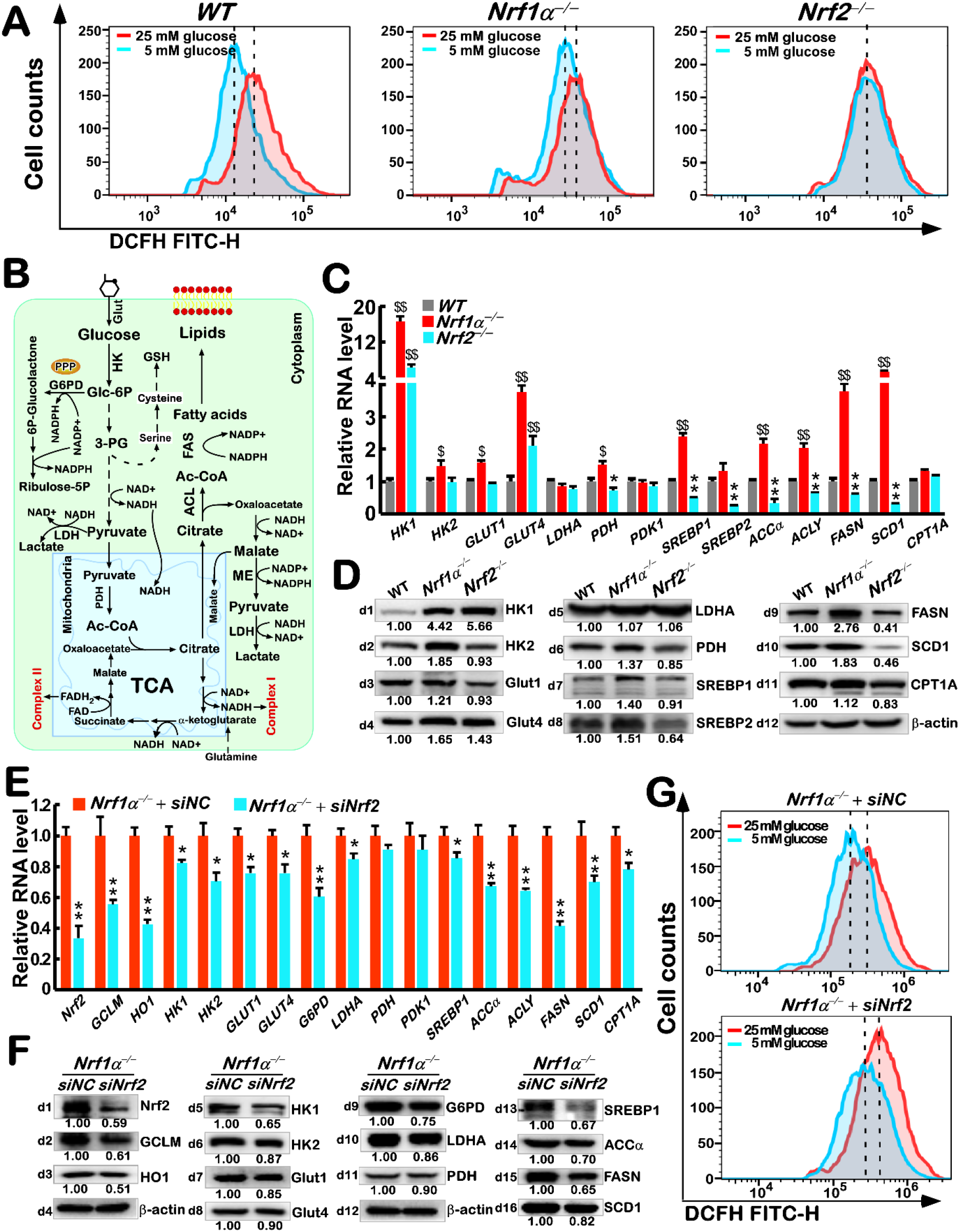
Expression of glycolysis-related genes and the effect of glycolysis on ROS. (A) Changes in ROS levels were detected by flow cytometry of *WT*, *Nrf1α^-/-^* and *Nrf2^-/-^* cell lines that had been cultured in 25 mM or 5 mM glucose media for 24 h, and then stained by DCFH for 30 minutes. (B) A schematic representation of redox metabolic process after glucose enters the cell. (C) Differential mRNA expression levels of *HK1, HK2, GLUT1, GLUT4, LDHA, PDH, PDK1, SREBP1, SREBP2, ACCα, ACLY, FASN, SCD1*, and *CPT1A* were determined by RT-qPCR in *WT*, *Nrf1α^-/-^* and *Nrf2^-/-^* cells. The data are shown as mean ± SEM (n= 3 × 3; $, p < 0.05 and $$, p < 0.01; *, p < 0.05 and **, p < 0.01). (D) Changed abundances of HK1, HK2, Glut1, Glut4, LDHA, PDH, SREBP1, SREBP2, FASN, SCD1, and CPT1A in *WT*, *Nrf1α^-/-^* and *Nrf2^-/-^* cells were visualized by Western blotting. The immunoblotting intensity was calculated as shown on the bottom. (E) The mRNA level of *Nrf2, GCLM, HO1, HK1, HK2, GLUT1, GLUT4, G6PD, LDHA, PDH, PDK1, SREBP1, ACCα, ACLY, FASN, SCD1*, and *CPT1A* were determined by RT-qPCR in *Nrf1α^-/-^* cells, which had been interfered by siRNA specific targeting Nrf2 (siNrf2) or scrambled siRNA (siNC). The resulting data are shown as mean ± SEM (n= 3 × 3; *, p < 0.05 and **, p < 0.01). (F) The protein levels of Nrf2, GCLM, HO1, HK1, HK2, Glut1, Glut4, G6PD, LDHA, PDH, SREBP1, ACCα FASN, and SCD1 were visualized by Western blotting in *Nrf1α^-/-^* cells that were interfered by siNrf2 or siNC. The intensity of all the immunoblots was calculated as shown on the bottom.

Those key genes involved in glucose and lipid metabolic pathways (as illustrated in Figure 4B) were further examined herein. Both RT-qPCR and WB results showed that hexokinases (HK1 and HK2, the first rate-limiting enzymes of glycolysis to yield glucose 6 phosphate), along with glucose transporters (Glut1, and Glut4) are all highly expressed in *Nrf1α^-/-^* cells, of which HK1 and Glut4 are also obviously up-regulated in *Nrf2^-/-^* cells to considerably higher levels, but that were rather lower than those obtained in *Nrf1α^-/-^* cells (Figure 4C, D). This implies an enhancement of glycolysis in *Nrf1α^-/-^* cells, as consistent with a previous report by pi et al (31). This is also evidenced by further examinations, revealing that pyruvate dehydrogenase (PDH) was modestly up-regulated in *Nrf1α^-/-^* cells (with hyperactive Nrf2 accumulated), and thus down-regulated in *Nrf2^-/-^* cells (Figure 4C, D). Rather, no changes in (G) Changes in ROS levels were detected by flow cytometry of *Nrf1α^-/-^* cells that had been interfered by siNrf2 or siNC and cultured in 25 mM or 5 mM glucose media for 24 h and then stained by DCFH for 30 minutes.basal expression of pyruvate dehydrogenase kinase 1 (PDK1, as a specific inhibitor of PDH) and lactate dehydrogenase A (LDHA) were observed in these experimental cells (Figure 4C, D). Such being the conditions, this facilitates the fuel loading by glycolysis insomuch as to enter the mitochondria, leading to an increased pressure on the oxidative respiratory chain to produce the excessive ROS in this organelles of *Nrf1α^-/-^* cells. Furthermore, it was also found that other critical genes *SREBP1, SREBP2 ACLY, ACCα, FASN* and *SCD1* (all responsible for fatty acid synthesis) were significantly up-regulated in *Nrf1α^-/-^* cells, but rather down-regulated in *Nrf2^-/-^* cells (Figure 4C, D). This implies that loss of *Nrf1α* rather than *Nrf2* results in an increase in fatty acid synthesis, providing a material basis for proliferation of *Nrf1α^-/-^* cells. Also, such increased synthesis of fatty acids has to consume a certain amount of NADPH generated by the pentose phosphate pathway, and hence leads to a decrease in antioxidant capability of the cell. Conversely, this appears to be supported by Scheffler’s group, demonstrating that a genetic respiratory chain deficiency could block the TCA cycle (50), serving as a vital hub towards *de novo* lipid synthesis (as illustrated in Figure 4B). Thereby, it is inferable that the elevated lipid synthesis is considered as an important outlet for glycolytic flux in *Nrf1α^-/-^* cells, as accompanied by its mitochondrial oxidative damages.

Since Nrf2 is aberrantly accumulated in *Nrf1α^-/-^* cells as aforementioned, the role of Nrf2 in this metabolic process was investigated by specific siRNAs interfering Nrf2. As shown in Figure 4(E, F), significant decreases in mRNA and protein expression of Nrf2 and target genes *HO1* and *GCLM* were caused by silencing of Nrf2 in *Nrf1α^-/-^* cells. In the meanwhile, all other examined genes except PDH and PDK (involved in glucose and lipid metabolism) are also down-regulated by knockdown of Nrf2 to varying degrees (Figure 4E,F). Among them, HK2, Glut1, Glut4, G6PD, ACCα, ACLY, FASN and SCD1 were substantially inhibited by interfering Nrf2 (Fig 4E, F). Further studies unraveled that such knockdown of Nrf2 in *Nrf1α^-/-^* cells did not reverse but shorten the reduced ROS led by 5mM glucose (Figures 4G), with the apoptosis unchanged (Figure S9A). In addition, knockdown of Nrf2 in *Nrf1α^-/-^* cells led to an increase in the intracellular ROS and apoptosis, when compared to equivalent controls (Figures S9B), implying an antioxidant cytoprotective role of Nrf2 against oxidative damages. Altogether, loss of Nrf1α may stimulate a surge of Nrf2 accumulated in order to enhance glycolysis and lipid synthesis; this is also accompanied by an increased pressure of the *Nrf1α^-/-^*-damaged mitochondria so as to generate the excessive ROS. In turn, the elimination of ROS by Nrf2-mediated antioxidant cytoprotective mechanism is also reinforced. Therefore, it is postulated that Nrf2 may provide a decisive interplay to balance between ROS arising from cellular metabolism and antioxidant response.

### Mitochondrial UCP2 is negatively regulated by Nrf2 *via* miR-195 and miR-497 to augment ROS, even in *Nrf1α^-/-^* cells

Besides the glycolytic flux to enter the mitochondria, its uncoupling proteins (UCPs) can also monitor the production of ROS in this organelles (51). Among them, UCP1 was reported to be positively correlated with Nrf1, but it is dominantly expressed in white adipose tissue (52), while UCP2 is widely expressed in various tissues, though they have an ability to reduce the mitochondrial Δψm by ‘mild uncoupling’ insomuch as to negatively regulate the yield of ROS(51,53). Here, our evidence revealed that basal mRNA and protein levels of *UCP2* were significantly decreased in *Nrf1α^-/-^* cells, but up-expressed to considerable higher extents in *Nrf2^-/-^* cells, when compared to *WT* controls (Figure 5A). Next, we examined the effect of UCP2 on production of ROS by forced expression of UCP2 in distinct genotypic cell lines (Figures 5B and S10A). The results unraveled that the ROS levels in *Nrf1α^-/-^* cells (with hyperactive Nrf2) were significantly decreased by over-expression of UCP2, but almost unaffected in *Nrf2^-/-^* and *WT* cell lines (Figure 5B). Collectively, these indicate that Nrf2 may exert a dominant inhibitory effect on UCP2 to promote the yield of ROS in *Nrf1α^-/-^* cells.

**Figure5.**
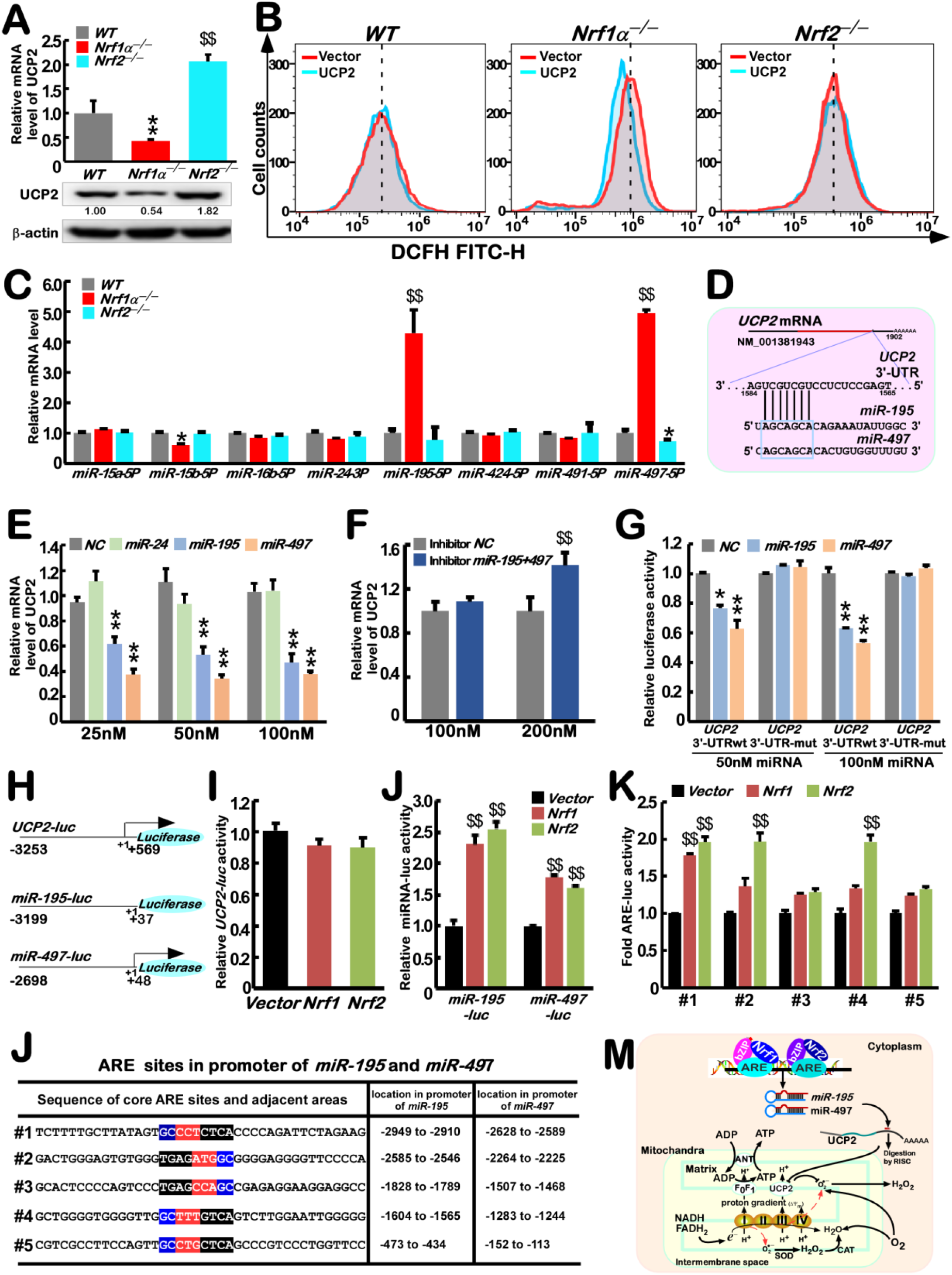
Nrf1 and Nrf2 affects ROS production by inhibiting UCP2 via miRNA-195 and miRNA-497. (A) The mRNA (*upper column*) and protein (*lower panel*) level of *UCP2* were determined by RT-qPCR and Western blotting of *WT*, *Nrf1α^-/-^* and *Nrf2^-/-^* cells. The qPCR data are shown as mean ± SEM (n= 3 × 3; $$, p < 0.01; **, p < 0.01). The intensity of all the immunoblots was calculated as shown on the bottom. (B) *WT*, *Nrf1α^-/-^* and *Nrf2^-/-^* cell lines were allowed for forced expression of UCP2 or empty vector, and then for 24-h recovery from transfection, before ROS levels of cells are detected by flow cytometry. (C) Relative mRNA level of *miR-15a-5P, miR-15b-5P, miR-16b-5P, miR-24-3P, miR-195-5P, miR-424-5P, miR-491-5P*, and *miR-497-5P* in *WT*, *Nrf1α^-/-^* and *Nrf2^-/-^* cells were determined by RT-qPCR. The data are shown as mean ± SEM (n= 3 × 3; $$, p < 0.01; *, p < 0.05). (D) Schematic representation of miR-195 and miR-497 targeting the 3’-UTR of *UCP2*. (E) WT cells were transfected with miR-24, miR-195 or miR-497 mimics in indicated concentration, and then allowed for 24-h recovery from transfection, because the mRNA levels of *UCP2* were determined by RT-qPCR. The data are shown as fold changes (mean ± SEM, n= 3 × 3; **, p < 0.01). (F) After *Nf1α^-/-^* cells were transfected with the inhibitors of miR-195 and miR-497, the mRNA levels of *UCP2* were determined by RT-qPCR. The data are shown as mean ± SEM (n= 3 × 3; $$, p < 0.01). (G). The 3’-UTR of *UCP2* were cloned into psiCHECK-2 plasmid, and co-transfected with pRL-TK (an internal control), plus miR-195, miR-497 or mutants, besides negative controls. After 24-h recovery from transfection, the miRNA activity of binding to 3’-UTR of *UCP2* was assayed by indicated luciferase reporters. The data are shown as fold changes (mean ± SEM, n= 3 × 3; **, p < 0.01). (H) Schematic representation of the *UCP2-luc, miR-195-luc and miR-497-luc* reporters, which were constructed into the PGL3-Promoter plasmid. These gene promoter regions were also indicated. (I) Relative activity of *UCP2-luc* were determined in the cells that were co-expressed with each of indicated luciferase reporters, and pRL-TK, plus an expression construct for Nrf1 or Nrf2, or empty pcDNA3 plasmid and then allowed for 24-h recovery from transfection. The resulting data are shown as mean ± SEM (n= 3 × 3). (J) Relative activity of *miR-195-luc and miR-497-luc* were determined as described above (mean ± SEM, n= 3 × 3; $$, p < 0.01). (K) Fold activity of *ARE-luc* activated by Nrf1 or Nrf2 was determined. These ARE-adjoining sequences listed in (L) were cloned into PGL3-promoter plasmid, and were co-transfected with pRL-TK, plus an expression construct for Nrf1 or Nrf2, or empty pcDNA3 plasmid. (L) Distinct locations of ARE sites from the promoters of *miR-195* and *miR-497* were listed herein. (M) A proposed model to explain distinct effects of Nrf1 and Nrf2 on ROS production by inhibiting UCP2 *via* miR195 and miR497.

To gain insight into such putative inhibitory effect of Nrf2 on UCP2, we further investigated this gene promotor and its mRNA regulatory mechanisms, As shown in Figure 5C, among eight candidate miRNAs for targeting 3’-UTR of *UCP2*’s mRNA predicated by online databases (http://www.targetscan.org/ and http://ophid.utoronto.ca/mirDIP/index.jsp#r), only miR-195 and miR-497 were highly expressed in *Nrf1α^-/-^* cells, but unaltered or even down-regulated, respectively, in *Nrf2^-/-^* cells, just as opposite to basal expression of UCP2 (Figure 5A). The sequence analysis of miR-195 and miR-497 deciphered a conserved homology targeting to their complementary 7-nucleotide motif within 3’-UTR of UCP2 (Figure 5D). As anticipated, the results from RT-qPCR revealed that the mRNA expression levels of UCP2 were effectively down-regulated by transfecting different concentrations of miR-195 or miR-497, but not miR-24, into *WT* cells (Figure 5E). In the parallel experiments, the decreased expression of *UCP2* in *Nrf1α^-/-^* cells was restored by a co-inhibitor of both miR-195 and miR-497 (Figure 5F).

To validate putative interaction of either miR-195 or miR-497 with *UCP2* mRNA, we cloned their 3’-UTR or mutants into a luciferase reporter vector before being co-transfected into *WT* cells. As expected, the results demonstrated that the 3’-UTR-driven reporter activity was inhibited by miR-195 or miR-497, but such this inhibitory effects were sufficiently abrogated by its mutants from 3’-UTR of *UCP2* mRNA (Figure 5G). Next, we constructed three reporter genes by cloning distinct lengths of the examined promoter regions of *UCP2, miR-195* and *miR-497* (i.e., *UCP2-Luc, miR-195-Luc*, and *miR-497-Luc*, Figure 5H), in order to verify whether they are regulated by Nrf1 and/or Nrf2. The luciferase assays showed that the *UCP2-Luc* transcriptional activity was almost unaffected by Nrf1 or Nrf2 (Figure 5I). However, transactivation activity of *miR-195-Luc* or *miR-497-Luc* reporters was significantly induced by co-transfection with Nrf1 or Nrf2 (Figure 5J). Further analysis of the promoter regions of *miR-195* and *miR-497* uncovered that their promoters are almost completely overlapped and comprise five common *ARE* consensus sites (Figure 5K). Among these ARE-driven luciferase reporters made by ligating the indicated sequences into the PGL3-promoter vector, only #1 transactivation activity was modestly mediated by Nrf1, while Nrf2 enabled to mediate significant induction of #1, #2 and #4 reporters (Figure 5L). Collectively, these indicate that Nrf2 (and Nrf1) possesses a capability to down-regulate the *UCP2* gene activity indirectly by cognate ARE-driven expression of miR-195 and/or miR-497 (Figure 5M). This notion is also further supported by additional evidence that a remarkable decrease of *UCP2* at its mRNA and protein expression levels occurred upon tetracycline-inducible expression of either Nrf1 or Nrf2 in HEK293 cells (Figure S10B).

### Loss of Nrf1α leads to inactivation of mitochondrial stress response, albeit Nrf2 is hyperactive, in *Nrf1α^-/-^* cells

In order to maintain the mitochondrial homeostasis, all eukaryotic cells have evolutionarily developed a nuclearly-controlled transcriptional responsive programme, called mitochondrial unfolded protein response (UPR^mt^)(54). If it is required for mitochondrial functioning, UPR^mt^ is triggered to actively promote the repair and recovery of mitochondrial function and integrity. In *Caenorhabditis elegans*, the UPR^mt^ is monitored primarily by the stress activated transcription factor ATFS-1 (of the basic-region leucine zipper family), as well by SKN-1 (skinhead-1, sharing an orthologous homology with Nrf1 and Nrf2) (55,56). The mammalian UPR^mt^ is regulated by ATF5-mediated expression of several mitochondrial chaperone and protease genes to promote its OXPHOS and cell growth during mitochondrial dysfunction (57). Besides, ATF4 and CHOP are also involved in activation of UPR^mt^ (58,59). Our previous study has found that loss of Nrf1 results in down-regulation of ER-stress related genes (Figure S11A, B)(60). Herein, our evidence revealed that basal expression of *ATF5, ATF4* and *CHOP*, along with their targets *HSP60, GRP75* (i.e. *mtHSP70*) and *FGF21* was, to different degrees, lowered in *Nrf1α^-/-^* cells (Figure S11B). Of note, these two key chaperones HSP60 and GRP75 are controlled by ATF5 in folding the denatured and nascent polypeptides in the mitochondria, while FGF21 is regulated by ATF4 in metabolic adaptation to fasting (61). By contrast with *Nrf1α^-/-^* cells, putative inactivation of UPR^mt^ did not appear to occur in *Nrf2^-/-^* cells, when compared with a similar status of *WT* cells (Figure S11B).

To determine the effect of Nrf1 or Nrf2 deficiency on UPR^mt^, *Nrf1α^-/-^* and *Nrf2^-/-^* cell lines were treated with carbonyl cyanide-p-trifluoromethoxyphenylhydrazone [(FCCP, an uncoupling reagent to disrupt ATP synthesis by transporting H^+^ ion through the mitochondrial membrane before being used for oxidative phosphorylation, thus to activate UPR^mt^ as reported by (59,62–64)]. The results unraveled that *Nrf1* and *Nrf2* at the mRNA and protein levels were obviously induced by FCCP in *WT* cells (Figure 6A, B). Albeit transcriptional expression of Nrf1 was reportedly regulated by Nrf2(37), it was still stimulated by FCCP in *Nrf2^-/-^* cells. Conversely, mRNA expression levels of Nrf2 were unaffected by FCCP in *Nrf1α^-/-^* cells, even though its protein appeared to be further accumulated (Figure 6A, B). Further examinations revealed that FCCP-inducible expression levels of *ATF4, ATF5, CHOP, HSP60, GRP75* and *FGF21* were almost completely abolished in *Nrf1α^-/-^* cells, but not or less altered in *Nrf2^-/-^* cells than those obtained from *WT* cells (Figure 6A, C). Collectively, these demonstrate that Nrf1, rather than Nrf2, is a dominant activator involved essentially in the FCCP-induced UPR^mt^.

**Figure 6.**
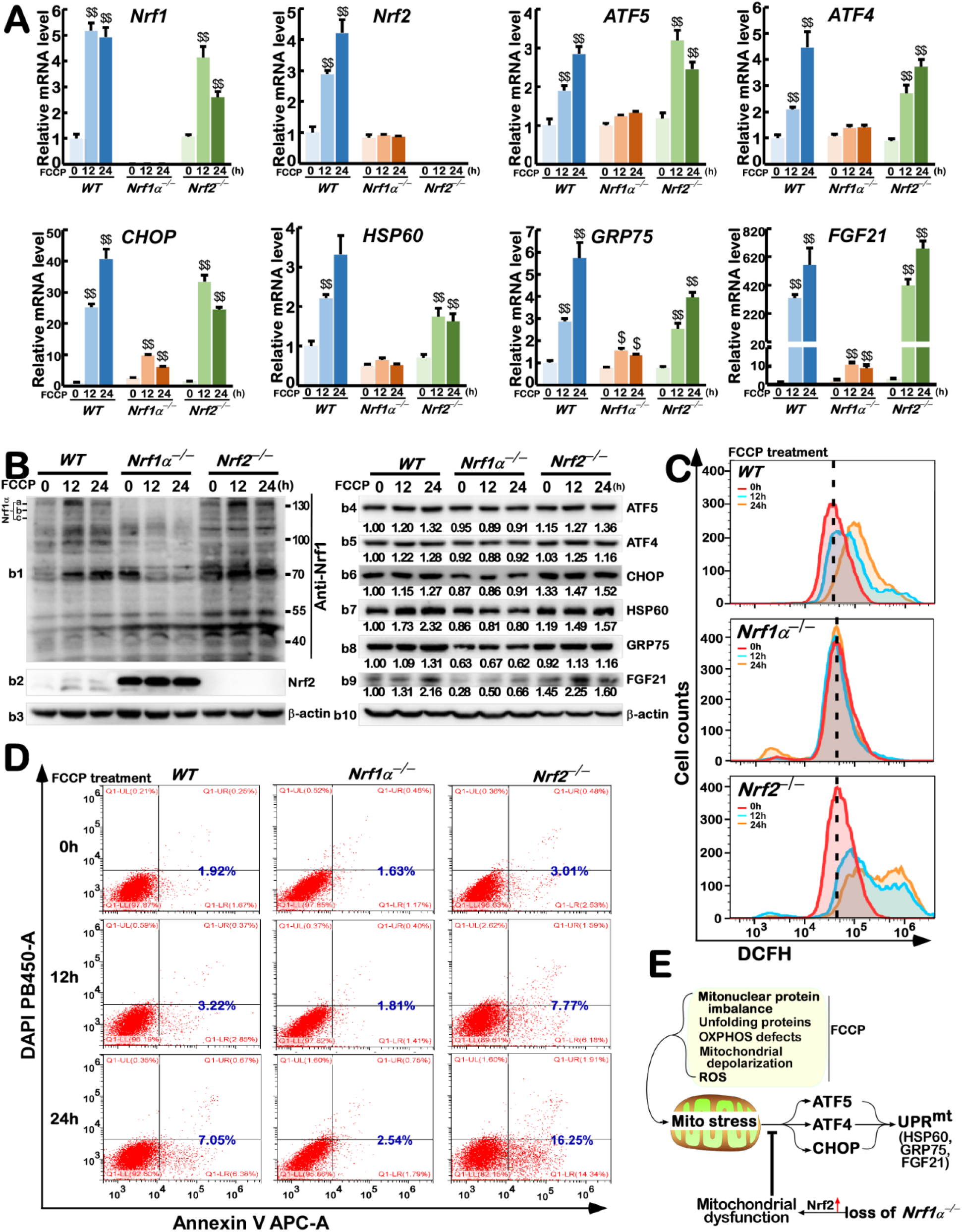
Knockout of Nrf1 inhibits the activation of mitochondrial stress related genes. (A) The mRNA levels of *Nrf1, Nrf2, ATF5, ATF4, CHOP, HSP60, GRP75*, and *FGF21* were determined by RT-qPCR in *WT*, *Nrf1α^-/-^* and *Nrf2^-/-^* cells, which had been treated with FCCP for 0 h, 12 h or 24 h. The data are shown as mean ± SEM (n= 3 × 3) with significant increases ($, p < 0.05 and $$, p < 0.01) as compared to the untreated cases. (B) The protein levels of indicated genes above were visualized by Western blotting in *WT*, *Nrf1α^-/-^* and *Nrf2^-/-^* cells, which had been treated with FCCP for 0 h, 12 h or 24 h. The intensity of all the immunoblots was calculated as shown on the bottom. (C) Changes in ROS levels were detected by flow cytometry in *WT*, *Nrf1α^-/-^* and *Nrf2^-/-^* cells that had been treated by FCCP for 0h, 12 h, or 24 h, and then stained by DCFH for 30 minutes. (D) The apoptosis was analyzed by flow cytometry in *WT*, *Nrf1α^-/-^* and *Nrf2^-/-^* cells, after they had been treated by FCCP for 0 h, 12 h, or 24 h and then incubated with Annexin V-FITC and PI. (E) A proposed model to explain an essential role of Nrf1 in mediating mitochondria unfolded protein response.

Further treatment of *WT* cells with FCCP caused a significant increment in production of ROS in a time-dependent manner (Figure 6D); this was also accompanied by increased apoptosis (Figure 6E). By sharp contrast, FCCP-caused ROS and apoptosis were substantially augmented in *Nrf2^-/-^* cells (Figure 6D, E). However, it is, much to our surprise, found that almost no changes in the intracellular ROS levels and apoptosis of *Nrf1α^-/-^* cells occurred after treatment of FCCP (Figure6 D,E). This implies that no proper UPR^mt^ to FCCP is instigated in the dysfunctional mitochondria caused by loss of Nrf1α, although hyperactive Nrf2 is retained in *Nrf1α^-/-^* cells (Figure 6F). Conversely, loss of Nrf2 may render its deficient cells to rely much on the energy supply of mitochondria.

### The EMT-relevant signaling pathways are constitutively activated in *Nrf1α^-/-^* cells

The above-described evidence revealed that mildly increased ROS in *Nrf2^-/-^* cells is attributable to diminishment of antioxidant capability to scavenge ROS, while *Nrf1α^-/-^* cells give rise to severe amount of ROS in impaired mitochondria, possibly leading to malignant cell proliferation as reported previously (37). Herein, to further determine the underlying mechanisms by which loss-of-function of Nrf1α enables to serve an original impact on tumorigenesis and development, we examined several discrete signaling pathways provoked by ROS, which are all converged on the EMT-relevant process involved in carcinogenesis and malignance (including invasion and migration). As illustrated in Figure 7A, the migrating ability of *Nrf1α^-/-^* cells through transwells was greatly enhanced, while migration of *Nrf2^-/-^* cells became rather weakened, when compared to that of *WT* cells. Further examinations revealed that the epithelial marker proteins E-cadherin (CDH1) was obviously down-regulated in *Nrf1α^-/-^* cells, but the mesenchymal marker proteins N-cadherin (CDH2), vimentin and fibronectin 1 (FN1) were apparently up-regulated in this deficient cells (Figure 7B, *b1 to b4*). Conversely, in *Nrf2^-/-^* cells, CDH1 was substantially incremented, while CDH2 and FN1 were roughly unaltered. Interestingly, ITGB4 (integrin subunit beta 4, a key node making the cell-cell interaction and communication with the extracellular matrix) was markedly increased in *Nrf1α^-/-^* cells, but rather diminished in *Nrf2^-/-^* cells (Figure 7B, *b6*). Aside from SNAI1 that was unaffected in these two deficient cell lines, SNAIL2 was significantly augmented in *Nrf1α^-/-^* cells, but was unchanged in *Nrf2^-/-^* cells (Figure 7B, *b7 & b8*), albeit both factors were shown to activate the EMT-relevant transcriptional programme during liver fibrosis and cancer development, but also repress the epithelial genes by binding to their E-box DNA sequences (65). In addition, matrix metallopeptidase 9 (MMP9, which can enhance the extracellular matrix protein degradation and hence enable invasion) was up-regulated in *Nrf1α^-/-^* cells, but down-regulated in *Nrf2^-/-^* cells (Figure 7B, *b9*). Taken altogether, these demonstrate that loss of Nrf1α leads to promotion of the EMT and malignant behavior, but such effects appear to be suppressed by loss of Nrf2, although its ROS levels were also increased.

**Figure 7.**
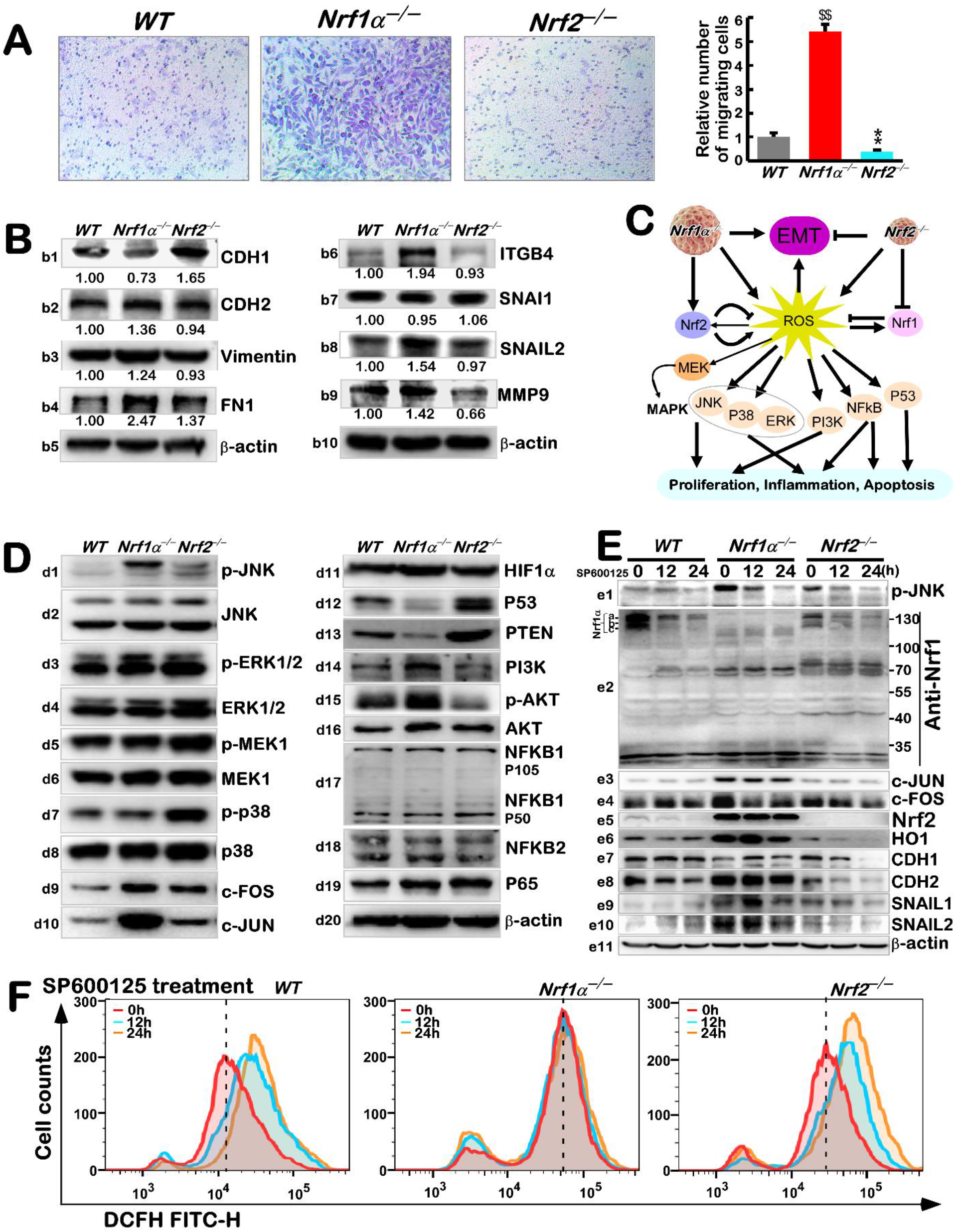
Effects of knockout of Nrf1 and Nrf2 on EMT-related genes and ROS-related pathways. (A) *WT*, *Nrf1α^-/-^* and *Nrf2^-/-^* cells were starved for 12 h in a serum-free medium and subjected to transwell migration, before Captured by microscope. Counts of migrated cells are shown as fold changes in right panel (mean ± SEM, n= 3 × 3; $$, p < 0.01; **, p < 0.01). (B) The protein levels of CDH1, CDH2, Vimentin, FN1, ITGB4, SNAI1, SNAIL2, MMP9, and P53 were visualized by Western blotting in *WT*, *Nrf1α^-/-^* and *Nrf2^-/-^* cells. The intensity of all the immunoblots was calculated as shown on the bottom. (C) A schematic representation of EMT and ROS-triggered signaling pathways. (D) Abundances of p-JNK, JNK, p-ERK1/2, ERK1/2, p-MEK1, MEK, p-p38, p38, HIF1α, PTEN, PI3K, p-AKT, NF-κB 2, NF-κB 1 (p105/p50), P65, and c-FOS, c-JUN proteins were visualized by Western blotting in *WT*, *Nrf1α^-/-^* and *Nrf2^-/-^* cells. The intensity of all the immunoblots was calculated and shown on Figure S12. (E) The protein levels of p-JNK, Nrf1, c-JUN, c-FOS, Nrf2, HO1, CDH, SNAIL were visualized by Western blotting in *WT*, *Nrf1α^-/-^* and *Nrf2^-/-^* cells, which had been treated with SP600125 for 0 h, 12 h or 24 h. (F) ROS levels were detected by flow cytometry in *WT*, *Nrf1α^-/-^* and *Nrf2^-/-^* cells, which had been treated by SP600125 for 0 h, 12 h, or 24 h, and then stained by DCFH for 30 minutes.

Such discrepant consequences in the migration of between *Nrf1α^-/-^* and *Nrf2^-/-^* cell lines, although both have given rise to evident increases in their intracellular ROS levels, suggest that distinct signaling pathways are likely to be activated by different extents of ROS-caused redox stress in the two distinctive genotypic cell lines (Figure 3C). Firstly, examinations of expression of mitogen-activated protein kinases [MAPKs, including ERKs, JNK and p38 kinase, that play key roles in tissue homeostasis, cell proliferation, differentiation survival and migration, as well in inflammation and carcinogenesis (66)] unraveled that phosphorylated abundance of JNK was substantially augmented in *Nrf1α^-/-^* cells, but only modestly increased in *Nrf2^-/-^* cells, albeit its total protein expression levels were slightly enhanced in the two cell lines (Figure 7D, *d1* & *d2*). Conversely, the phosphorylated ERKs and p38, as well as their upstream kinase MEK were strikingly elevated in *Nrf2^-/-^* cells, but almost unchanged in *Nrf1α^-/-^* cells (Figure 7D, *d3 to d8*). Their downstream c-JUN and c-FOS (two key constituents of AP1 involved in tumorigenesis) were markedly up-regulated in both *Nrf1α^-/-^* and *Nrf2^-/-^* cell lines (Figure 7D, *d9 & d10*). Such increased c-JUN abundance was nearly 6 folds, while the increased c-FOS was also nearly 4 folds in *Nrf1α^-/-^* cells, but both proteins were about 2 folds in *Nrf2^-/-^* cells, as compared to *WT* controls (Figure S12). Thereafter, it was found that HIF1α (a critical factor for hypoxic stress response) was obviously increased in *Nrf1α^-/-^* cells, but almost unaffected in *Nrf2^-/-^* cells (Figure 7D, *d11*). However, the versatile p53 expression was almost abrogated in *Nrf1α^-/-^* cells, but significantly up-regulated in *Nrf2^-/-^* cells (Figure 7D, *d12*).

Next, basal expression of key signal molecules involving the PTEN-PI3K-AKT and NF-κB pathways were examined, albeit a previous study showed that PTEN [a potent tumor suppressor, that negatively regulates the PI3K-AKT signaling to inhibit cell proliferation and survival (67)] is blocked by ROS-induced Nrf2 through miR-22 (37). Herein, basal PTEN expression was almost completely abolished in *Nrf1α^-/-^* cells, but conversely remarkably increased in *Nrf2^-/-^* cells (Figure 7D, *d13*). Accordingly, both phosphorylated proteins of PI3K (p110) and AKT were significantly up-regulated in *Nrf1α^-/-^* cells, but rather down-regulated in *Nrf2^-/-^* cells (Figure 7D, *d14* to *d15*). This finding provides a better understanding of malignant proliferation of tumors occurring upon loss of Nrf1α. Furthermore, only p65 subunit of NF-κB (as a key factor responsible for the inflammatory response to ROS) was only modestly up-regulated in both *Nrf1α^-/-^* and *Nrf2^-/-^* cell lines (Figure 7D, *d19*), but the other relevant NF-κB1 and NF-κB2 were unchanged (Figure 7D, *d17 & d18*).

Lastly, treatments of *WT* cells with a JNK-specific inhibitor SP600125 revealed that Nrf1, Nrf2 and all relevant target genes were suppressed significantly (Figure 7E). Similar inhibitory effects were also exerted in *Nrf2^-/-^* cells. By contrast, although auto-phosphorylated JNK, along with c-JUN and c-FOS in *Nrf1α^-/-^* cells were strikingly prevented by SP600125 (Figure 7E), Nrf2, CDH1 and CDH2 were largely unaffected by SP600125, while HO-1, SNAIL1 and SNAIL1 were partially altered (*e5 to e10*). This implies they may be modulated independently of the JNK-AP1 signaling, particularly in *Nrf1α^-/-^* cells. Intriguingly, inhibition of JNK caused a sharp increment intracellular ROS in *WT* and *Nrf2^-/-^* cells, but nearly unchanged in *Nrf1α^-/-^* cells (Figure 7F), reflecting a crucial cytoprotective role of JNK and AP1 in the response to oxidative stress. Overall, loss of *Nrf1α^-/-^* or *Nrf2^-/-^* leads to differential dysregulation of multi-hierarchical signaling pathways to cognate target gene networks at distinct layers, by varying extents of their endogenous oxidative stress during different cell processes (Figure 8).

**Figure8.**
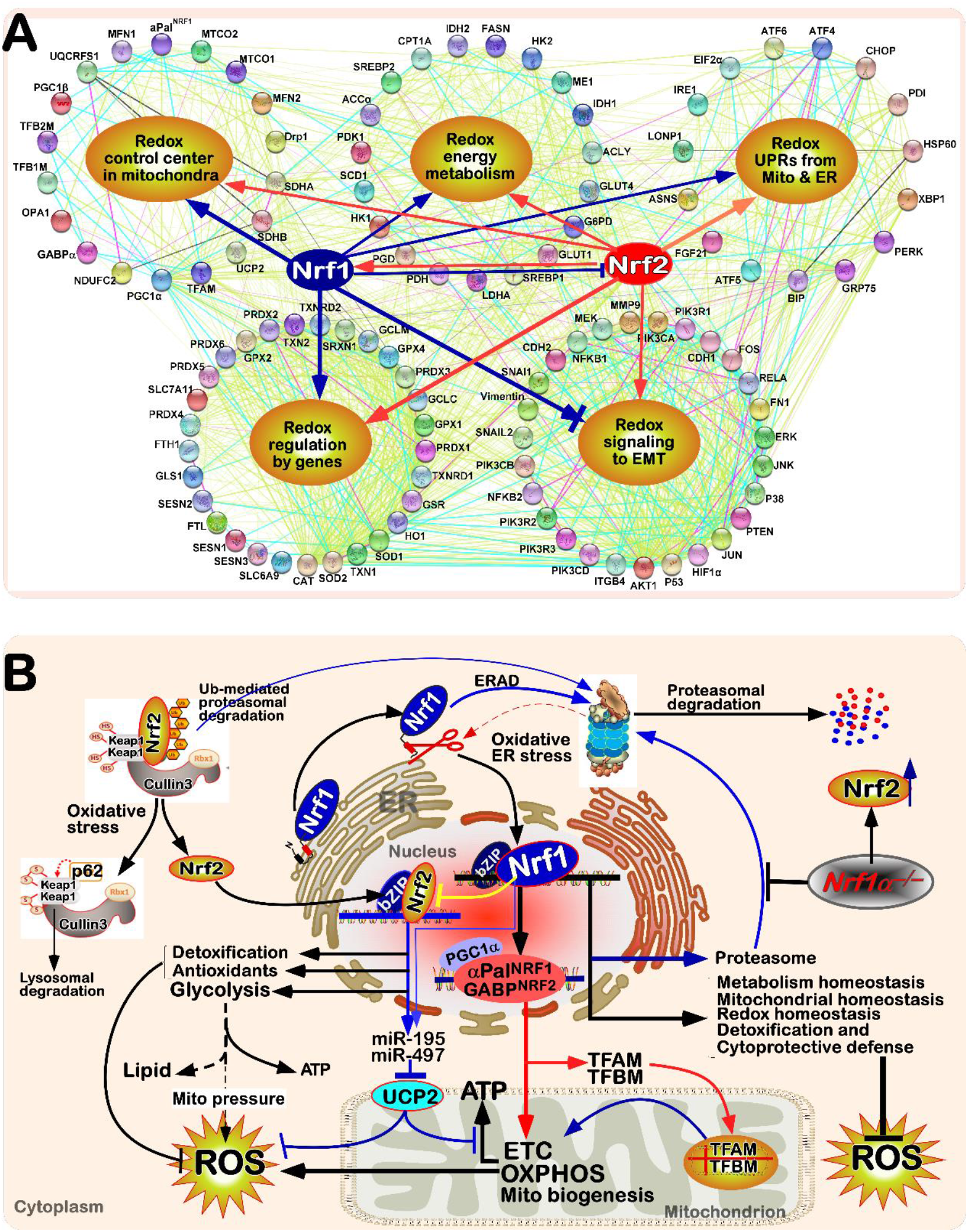
The functional protein association networks and regulatory model of Nrf1 and Nrf2. (A) A multi-hierarchical regulatory network monitored by Nrf1 and Nrf2 alone or together. Those protein-protein associations are determined by various ways, and thus represented by different colored edges as indicated. (B) A comprehensive regulatory model is proposed to explain an indispensable function of Nrf1 as a redox-determining factor for mitochondrial homeostasis.

## DISCUSSION

Just because ROS have a ‘double-edged sword’s effect with dual characteristics of beneficial hormesis and harmful cytotoxicity to all living cells, a steady-state of redox homeostasis is robustly maintained by balancing ROS production (primarily in mitochondria) and elimination by a set of antioxidant responses and detoxification protective systems. Such a certain homeodynamic range of redox threshold should be tightly controlled by redox signaling to gene regulatory networks. In this study, we have provided a holistic view of distinct roles for Nrf1 and Nrf2 played in redox regulation by distinctive mechanisms.

### Distinct roles of Nrf1 and Nrf2 in redox regulation

Herein, we found that loss of either Nrf1 or Nrf2 leads to a redox imbalance so as to increased levels of intracellular ROS, which occurs by distinct rational mechanisms. This is because knockout of Nrf2 results in a reduction of its cellular capability to eliminate ROS insomuch as to make a redox bias towards yield of ROS. By sharp contrast, similar capability to eliminate ROS appears to be undiminished by knockout of Nrf1, but conversely the production of ROS in the mitochondria is increased strikingly in *Nrf1α^-/-^* cells. Consequently, Nrf2 is accumulated in *Nrf1α^-/-^*-deficient cells by its proteasomal dysfunction and further activated by increased ROS, leading to an evident enhancement of antioxidant, detoxifying and cytoprotective systems. In turn, the activity of Nrf1 appears to be not further reinforced by elevated ROS in *Nrf2^-/-^* cells. This implies that Nrf1 acts as a key constitutive transcription factor of redox regulation, but it is activated predominantly by another main way rather than ROS, or the ROS activation of Nrf1 is dependent on Nrf2. Coincidently, this is also fully consistent with our previous finding that Nrf2 serves as an upstream regulator to activate the transcriptional expression of Nrf1(37). Of note, as an ER-anchored membrane protein, Nrf1 is *de facto* regulated by ER-derived unfolded proteins and/or other metabolic stressors(20), to coordinate the ER homeostasis with redox homeostasis (Figure 8), but this is required for further experiments to be elucidated.

With an aberrant accumulation of hyperactive Nrf2 in *Nrf1α^-/-^* cells, most of antioxidant and detoxifying enzymes, as well as relevant genes responsible for elimination of ROS are markedly augmented. Many of these up-expressed genes (e.g., encoding PRDX5, PRDX6, GPX1, SOD2, SESN2, G6PD, GCLC, GCLM, GLS1, GSR, SLC6A9, FTH1, HO1, TXN1 and TXN2) are diminished or abolished by knockout of *Nrf2^-/-^*, indicating they are Nrf2-specific target genes. Also, it cannot be ruled out that some genes are, to certain lesser extents, regulated by Nrf1, but its overlapping effects are much likely to be concealed by hyperactive Nrf2. For example, SRXN1 and PGD were reportedly regulated by Nrf2 (68,69), and thus down-regulated in *Nrf2^-/-^* cells, but largely unchanged in *Nrf1α^-/-^* cells, implying both are co-regulated by Nrf1 and Nrf2, but the regulatory effect of Nrf1 is postulated to be offset by hyperactive Nrf2. Conversely, SESN3 was down-regulated in *Nrf1α^-/-^* cells, but almost completely abrogated in *Nrf2^-/-^* cells, implying that this putative effect of Nrf1 on SESN3 was only partially counteracted by accumulated Nrf2. In addition, PRDX4, SESN1 and TIGAR are elevated in both *Nrf1α^-/-^* and *Nrf2^-/-^* cell lines, suggesting they may be over-stimulated by elevated ROS.

### Nrf1 serves as an indispensable redox-determining factor in the mitochondrial homeostasis

Albeit two independent groups had presumed that the loss of Nrf1 is likely to cause an impairment of mitochondrial functioning (29,52), no further experimental evidence has been provided in the current literature. Another group led by Yoon reported that Nrf1 acts as one of the most responsive genes when the mitochondrial respiratory chain is damaged (70,71). These putative effects of Nrf1 prompt us to gain insights into its redox regulation for mitochondrial homeostasis, because this organelle is a main resource of intracellular ROS production. The experimental evidence has been presented here, revealing that a dramatic increase of the mitochondrial ROS levels was manifested in *Nrf1α^-/-^* cells, simultaneously as accompanied by significant down-regulation of key mitochondrial complex subunits upon loss of its function. However, no further increases or even modest decreases in the yield of ROS were observed after *Nrf1α^-/-^* cells had been treated with either the mitochondrial respiratory chain inhibitor rotenone or another uncoupling agent FCCP. Such dysfunctional mitochondria in *Nrf1α^-/-^* cells were damaged as evidenced by further electroscopic observations. As such, Nrf2 is also inferred to be partially involved in mitochondrial redox stress response, because rotenone treatment of *Nrf2^-/-^* cells only modestly increased ROS generation to certain extents that are not higher than *WT* controls. But, remarkable changes in its ROS occurred after FCCP treatment of *Nrf2^-/-^* cells, implying that Nrf1 remains to be expressed to a considerably high degree in *Nrf2^-/-^* cells, so that deregulated mitochondria were further stressed by FCCP insomuch as to trigger excessive ROS products. In addition, it should also be noted that basal metabolism of *Nrf2^-/-^* cells is likely placed at a rather lower level, so to enable this deficient lines to make more resistance to glucose starvation, as reported previously (42).

It is of crucial importance to discover that loss of Nrf1α results in a defective down-regulation or even abolishment of the nuclearly-controlled mitochondrial respiratory factors involving PGC1α-αPal^NRF1^ and PGC1α-GABPα^NRF2^ pathways in *Nrf1α^-/-^* cells. Of note, PGC1α is a key transcriptional coactivator enabling for activation of estrogen-related receptors, peroxisome proliferator-activated receptors (PPARs), and other specific partners, in addition to two nuclear respiratory factors αPal^NRF1^and GABPα^NRF2^ (Figures 3I and 8B). Co-activation of nuclear respiratory factors αPal^NRF1^and GABPα^NRF2^ by PGC1α further induce the expression of mitochondrially-specific transcriptional factors TFAM and TFBM, which can bind directly to both strands of mtDNA and thus play key roles in the mtDNA replication, transcription and maintenance (72). In addition to regulating the expression of these mitochondrially-located genes through TFAM and/or TFBM, two nuclear respiratory factors αPal^NRF1^and GABPα^NRF2^ also directly activate those nuclearly-located genes involved in the oxidative respiratory chain and mitochondrial biogenesis. Herein, our experimental evidence has unraveled that overexpression of Nrf1 and Nrf2 activates the transcriptional expression of αPal^NRF1^, GABPα^NRF2^ and PGC1α, as well as respiratory chain subunits, and that they also exert transcriptional activation effects on each of αPal^NRF1^, GABPα^NRF2^ and PGC1α promoter-driven reporter genes. However, our further experimental evidence obtained from *Nrf1α^-/-^* cells has demonstrated that Nrf1, rather than Nrf2, can serve as a vital dominant-positive determinant of the mitochondrial functioning by governing transcriptional expression of αPal^NRF1^, GABPα^NRF2^ and PGC1α, because even though hyperactive Nrf2 was accumulated, it cannot compensate for a fatal defect in basal expression of αPal^NRF1^, GABPα^NRF2^ and PGC1α in *Nrf1α^-/-^* cells. But, not a decrease of PGC1β was determined in *Nrf1α^-/-^* cells, although it was reportedly regulated by Nrf1 in mouse cells (29); this difference may be attributable to distinct cell types between the human and mouse species. Moreover, when cells are required for impaired mitochondrial function, a highly-conserved mechanism called UPR^mt^ is activated to trigger cell repair of the mitochondrial function. Such UPR^mt^-related genes were still activated by FCCP-induced mitochondrial stress in *Nrf2^-/-^* cells, but almost unaffected in *Nrf1α^-/-^* cells. Thereby, it is inferable that the mitochondrial UPR^mt^ is prevented or even completely abolished in *Nrf1α^-/-^* cells, due to its fetal mitochondrial damage arising from loss of Nrf1α’s function. This finding further indicates that Nrf1, but not Nrf2, is an indispensable dominantly-determining factor in maintaining the mitochondrial homeostasis and its integrity, albeit Nrf2 is also involved in this process.

### Nrf2 exerts a ‘double-edged sword’s role in the redox regulation, particularly in *Nrf1α^-/-^* cells

Mitochondria produce ROS mainly from escaped electrons in the transport respiratory chain, and these electrons [e^-^, together with hydrogen ion (H^+^)] are originated primarily from aerobic glycolysis. Herein, it is, to our surprise, found that the apparent rise and fall of the intracellular ROS levels in *Nrf1α^-/-^* cells are accompanied by an increase or decrease of glycolysis, respectively when they had been treated with its activator insulin or inhibitor 2-DG. This is also supported by additional observations showing that the yield of ROS in 5mM-glucose cultured *Nrf1α^-/-^* cells is at a rather lower level than that arising from the 25mM glucose cultured conditions. Similar glucose effects on yield of ROS were also manifested in wild-type cells, but *Nrf2^-/-^* cells appeared to make little difference. Further experiments revealed that basal expression levels of glycolysis and lipid synthesis-related genes were up-regulated in *Nrf1α^-/-^* cells, but also enabled to be interfered to considerably lower extents by silencing of Nrf2. Collectively, these data suggest that hyperactive Nrf2 accumulated in *Nrf1α^-/-^* cells has a strong ability to promote glycolysis and lipid biosynthesis insomuch as to give rise to excessive ROS from its mitochondria, particularly under the pressure of a high glycolysis flow. In addition, we found that insulin can increase the expression of Nrf1, and 5mM glucose cultured conditions (glucose restricted medium) reduces the expression of Nrf1 (Figure S8C, D). When cells are treated with high concentrations of glycolysis inhibitors (2-DG) or starved in a non-glucose medium, glycosylated Nrf1 decrease and the active isoform of Nrf1 significant increase (Figure S8E)(60). This also implies that relevancy between Nrf1 and glucose metabolism. Perhaps under lower metabolic conditions Nrf1 is down-regulated to weaken mitochondrial-related metabolism, and under extreme glucose deficiency conditions Nrf1 is transitorily activated to enhance mitochondrial function to meet the needs of energy and promotes metabolism of non-sugar substances such as lipids, but that requires more experiments to prove.

Among those genes possibly mediated by Nrf2, PDH plays a role in controlling the entry of pyruvate produced by glycolysis into mitochondria, particularly when PDH is elevated concomitantly with unchanged LDHA, in *Nrf1α^-/-^* cells, making the flue flow produced by glycolysis more easily enter the mitochondria. However, a fatal damage of the electron transport chain slows down the TCA cycle(73), it is hence difficult for the impaired TCA cycle to consume acetyl-CoA that enters the mitochondria in *Nrf1α^-/-^* cells. The resulting increase of fatty acid biosynthesis provoked by hyperactive Nrf2 provides an alternative way to consume the metabolites from the increased glycolysis flow. This enables to allow the citric acid produced by oxaloacetate and acetyl-CoA to skip most of the damaged TCA cycle and then convert it into apple acids. Overall, deficiency of Nrf1α cannot only lead to a marked increase in intracellular ROS yield from its defective mitochondria, and also augment glucose consumption and lipid biosynthesis by increased Nrf2 so as to aggravate ROS production. Such incremented lipid biosynthesis may be explained as a reasonable cause of lipid deposition in *Nrf1α^-/-^* cells, ultimately resulting in spontaneous development of non-alcoholic steatohepatitis and ensuing hepatoma in liver-specific *Nrf1^-/-^* mice (35,37).

Notably, a direct main source of ROS arises from the electron escape of NADH produced by cellular metabolism in the respiratory chain. Such escaped electrons enable to exacerbate the mitochondrial membrane potential (Δψ m) to a rather higher degree due to the increasingly sluggish electron transport within a longer half-life of relevant respiratory chain intermediates(74). This is distinctive from destructive uncoupling, because a modest decrease in Δψm, called ‘mild uncoupling’, has been shown to exert a cytoprotective effect (75). Thereby, we also found that UCP2 is negatively regulated by hyperactive Nrf2 in *Nrf1α^-/-^* cells so as to increase the intracellular ROS production, even though Nrf2 has been generally accepted as a master regulator of antioxidant and detoxifying system and also as a key player in metabolic regulation. When required for cellular demands, UCP2 is widely expressed, so that it cannot only relieve the pressure of the electron transport chain and also reduce byproducts of ROS from mitochondria, but the resulting production of ATP is decreased concomitantly by reducing Δψm. Thus, it is inferable that Nrf2 is selectively allowed for inhibition of UCP2 to facilitate the energetic production of ATP, albeit with an accompanying increase in the mitochondrial ROS byproducts, particularly in *Nrf1α^-/-^* cells. Conversely, when UCP2 was over-expressed, the yield of ROS in *Nrf1α^-/-^* cells was greatly reduced, but in *Nrf2^-/-^* cells was unchanged similarly to *WT* controls, implying that UCP2 is also selectively contributable to antioxidant cytoprotection. Further experiments unraveled that UCP2 is negatively regulated by Nrf2 through ARE-driven miR-195 and/or miR-497. The expression levels of miR-195 and miR-497 were up-regulated by hyperactive Nrf2 in *Nrf1α^-/-^* cells, even albeit both can also be regulated directly by Nrf1. Altogether, these demonstrate that Nrf2 cannot only exert its intrinsic antioxidant effect, but also promote the generation of ROS within proper tempo-spatial contexts. And it also indicates that Nrf1 and Nrf2 maintain the balance between redox homeostasis and energy supply.

### Inter-regulatory effects of Nrf1 and Nrf2 are integrated by multiple signaling to gene expression networks

Several studies have uncovered that Nrf2 acts as a versatile chameleon-like player in both cancer prevention and promotion (76), but its oncogenic activity to promote cancer development is tightly confined by Nrf1 (along with its long isoform TCF11), which is endowed as a potent tumor-repressor(34,37,77). Such oppositely inter-regulatory effects of Nrf1 and Nrf2 are unified integrally by multiple redox signaling pathways to cognate gene expression networks at distinct levels (Figure 8A). These multi-hierarchical signaling networks were actually gradually converged on the base of redox regulation by Nrf1 and Nrf2 (Figure 8B). Of note, the primary ROS-caused stress and secondary oxidative damages are well known to serve as an initiator of cancer development and also as a promotor of cancer progression and malignance (78). However, our evidence revealed that significantly elevated ROS in *Nrf2^-/-^* cells does not render its migration ability and growth rate to be increased, but conversely become weaker than *WT* controls. Further xenograft animal experiments by inoculating *Nrf2^-/-^* cells unraveled that loss of Nrf2’s function leads to a striking reduction in *in vivo* malgrowth of hepatoma and its metastasis (37). These indicate that loss of Nrf2-mediated antioxidant and detoxifying cytoprotection results in certain ‘mild extents’ of ROS that are not enough to sufficiently initiate cancer development or promote malignant progression, but conversely can serve as a eustress to stimulate beneficial hormesis effects.

By contrast with *Nrf2^-/-^* cells, severe extents of ROS-caused distress and damages are determined in *Nrf1α^-/-^* cells, although as accompanied by hyperactive Nrf2 accumulation and reinforced antioxidant response. Such oxidative distress is attributable primarily to *Nrf1α^-/-^*-leading mitochondrial dysfunction and its UPR^mt^ failure, and secondarily to Nrf2-augmenting the yield of ROS by strengthening glycolysis (resembling the Warburg’s effect) to aggravate the pressure of electron respiratory chain and also inhibiting UCP2 to enable for electron escape from the transport respiratory chain. Consequently, loss of *Nrf1α^-/-^*-results in marked increases in its malgrowth, migration and metastasis, as well in the EMT process (37,79); such increased ability was also greatly prevented by silencing of Nrf2, in addition to restored expression of Nrf1α or TCF11 (37,77). These indicate that Nrf2 acts as a tumor-promoter to exert its oncogenic activity, only when Nrf1α/TCF11 is disrupted, whereas the latter Nrf1α/TCF11 is likely to possess an intrinsic capability to repress initiation of cancer development. This cancer-repressing effect may also be executed by two additional ‘star’ tumor-repressors PTEN and p53, because both were diminished in *Nrf1α^-/-^* cells. Besides, the EMT-relevant signaling, together with the MEK-MAPKs (i.e., JNK, ERKs, p38 kinase)-AP1 (JUN+FOS) and PI3K-AKT, as well as HIF1α and NF-κB, signaling networks are constitutively activated in *Nrf1α^-/-^* cells. In addition, the redox metabolic reprogramming of *Nrf1α^-/-^* cells (owing to dysfunctional mitochondria) may be contributable to cancer development and progression (42). Contrarily, most cell killing results from further elevated yield of ROS after *Nrf1α^-/-^* cells were stimulated by glucose deprivation. Overall, Nrf1 acts as a potent integrator of the cellular redox regulation by multi-hierarchical signaling to gene expression networks in order to maintain cell homeostasis and organ integrity.

In summary, systematic examinations of distinctive roles for Nrf1 and Nrf2 in redox regulation are carried out from a holistic view in combination of reductionist approaches. The results unveiled that Nrf1 functions as an indispensable redox-determining factor for mitochondrial homeostasis, because loss of its function leads to a potentially fetal defect in dysfunctional mitochondria to give rise to severe oxidative stress along with the failure of UPR^mt^, and such detriment effects cannot be counteracted by hyperactive Nrf2 accumulated in *Nrf1α^-/-^* cells, but conversely malgrowth of *Nrf1α^-/-^*-derived tumor appears to be protected by Nrf2-strengthening antioxidant response and glycolysis pathways. From this, it is inferable that Nrf2 plays a critical role for the Warburg’s effect in *Nrf1α^-/-^* cells. The reinforced glycolysis aggravates the mitochondrial pressure to allow for the electron escape to yield excessive ROS, particularly when UCP2 is suppressed by hyperactive Nrf2. Therefore, the versatile Nrf2 exerts a ‘double-edged sword’s role in redox regulation, particularly in *Nrf1α^-/-^* cells. Such inter-regulatory effects of Nrf1 and Nrf2 are integrated by multi-hierarchical signaling towards gene expression networks in order to perpetuate cell homeostasis and organ integrity during development and growth.

## Materials and Methods

### Cell lines, Culture and Transfection

The *Nrf1^-/-^* cells were established by TALENs-led genome editing(34). The *Nrf2^-/-^* cells were constructed by CRISPR/Cas9-editing system(37). These cell lines expressing Nrf1α, Nrf2, as wellas an empty control, were established by using the Flp-In™ TREx™-293 system (Invitrogen)(77). These cell lines were maintained in DMEM supplemented with 5 mM glutamine, 10% (v/v) foetal bovine serum (FBS), 100 units/mL of either of penicillin and streptomycin, in the 37°C incubator with 5% CO2. In addition, some of cell lines were transfected for 8 h with the indicated constructs mixed with the Lipofectamine®3000 agent in the *Opti*-MEM (gibca, Waltham, MA, USA). The cells were then allowed for recovery from transfection in a fresh complete medium for 24 h, before the other experiments were conducted.

### Plasmid construction

The expression constructs for human Nrf1 and Nrf2 were here created by inserting their full-length cDNA sequences into the KpnI/XbaI site of pcDNA3.1/V5His B. The six reporter gene plasmid for PGC1α-luc, αPal^NRF1^-luc, GABPα-luc, UCP2-luc, miR-195-luc and miR-497-luc were created by inserting their promoter sequences into the PGL3-basic vector, the promoter ranges of each gene are indicated in figure 3E and figure 5H. In addition, the ARE-luc plasmids were created by inserting ARE-adjoining sequences from corresponding gene promoters into the PGL3-promoter vector, the ARE-adjoining sequences are indicated in figure 3G and figure 5J. 3’-UTR of UCP2 were cloned into psiCHECK-2 plasmid.

### Real-time qPCR analysis of mRNA expression

Their total RNAs were extracted by using an RNA extraction kit (TIANGEN, Beijing, China), then approximately 2.0 μg of total RNAs were added in a reverse-transcriptase reaction to generate the first strand of cDNA (by using the Revert Aid First Strand Synthesis Kit, Thermo, Waltham, MA, USA). The synthesized cDNA was served as the template for qPCR, in the GoTaq®qPCR Master Mix (Promega, Madison, WI, USA). Subsequently, relative mRNA expression levels were measured by qRT-PCR with indicated primers. Of note, the mRNA expression level of β-actin served as an optimal internal standard control, relative to other mRNA expression levels presented as fold-changes. All the forward and reverse primers of those indicated genes were shown in Table S1.

### Western blotting

Experimental cells were harvested in a denatured lysis buffer (0.5% SDS, 0.04 mol/L DTT, pH 7.5, containing 1 tablet of cOmplete protease inhibitor EASYpacks in 10 ml of this buffer). The total lysates were further denatured by boiling at 100°C for 10 min, sonicated sufficiently, and diluted with 3× loading buffer (187.5 mmol/L Tris-HCl, pH 6.8, 6% SDS, 30% Glycerol, 150 mmol/L DTT, 0.3% Bromphenol Blue), before being re-boiled at 100°C for 5 min. Thereafter, equal amounts of protein extracts were subjected to separation SDS-PAGE before being transferred to polyvinylidene fluoride (PVDF) membranes (Millipore, Billerica, MA, USA), and subsequent visualization by Western blotting with distinct antibodies as indicated. β-actin served as an internal control to verify equal loading of proteins in each of electrophoretic wells.

### Intracellular ROS detection

Experimental cells incubated in a serum-free medium containing 20,70-Dichlorodihydrofluorescein diacetate (DCFH-DA, S0033, Beyotime, Shanghai, China), 3’-(p-hydroxyphenyl) fluorescein (HPA, H36004, ThermoFisher) or MitoSOX Red (40778ES50, Yeasen Biotechnology, Shanghai, China) at 37 °C for 20 min. After these cells were washed three times with a fresh serum-free medium, fluorescence intensity is measured by flow cytometer or fluorescent inverted microscope in specific excitation light (Ex) and emission(Em) light. Among them Ex/Em of DCFH-DA is 488/525nm, Ex/Em of HPF is 490/515nm, Ex/Em of MitoSOX is 510/580.

### Luciferase Reporter Assay

Equal numbers (1.0 × 10^5^) of COS1 cells were allowed for growth in each well of 12-well plates. After reaching 70-80% confluence, the cells were co-transfected for 8 h with an indicated luciferase plasmid alone or together with one of expression constructs with the Lipofectamine®3000 agent in the *Opti*-MEM (gibca, Waltham, MA, USA), meanwhile the Renilla-expressing pRL-TK plasmid served as an internal control for transfection efficiency. After cultured in a fresh complete medium for 24 h, the cells were lysed and the luciferase activity was measured by the dual-reporter assay (Promega, Madison, WI, USA).

### Transmission electron microscopy

Experimental cells were washed with PBS, and centrifuged (1000rpm) to form a cell mass before fixed with 2.5% glutaraldehyde in 0.1 M sodium cacodylate buffer pH 7.4 for 30 min at 37 °*C*. The fixed cells were stored in at 4 °*C* in 0.1 M sodium cacodylate buffer until postfixation, performed with a mixture of 1% OsO4, 1.5% K4Fe(CN)6 in 0.1 M sodium cacodylate pH 7.4 for 1 h at 4°*C* and overnight incubation in 0.25% uranyl acetate at 4 °*C*. After three water washes, samples were dehydrated in series of 15 min steps in 25%, 50%, 75%, 95% and 100% (v/v) ethanol and embedded in an epoxy resin (Sigma-Aldrich). Ultrathin sections (60–70 nm) were obtained with a Leica EM UC7 ultramicrotome and counterstained with 1% uranyl acetate for 15 min and 1% lead citrate for 6 min. Thin sections were imaged using a HITACHI HT 7800 120kv transmission electron microscope operating at 80kV.

### Flow cytometry analysis of apoptosis

Experimental cells (3 × 10^4^) were allowed for growth in each well of 6-well plates. After transfection or drug treatment, the cells were pelleted by centrifuging at 1000×g for 5 min and washed by PBS for three times, before being incubated for 15 min with 5 μL of Annexin V-FITC and 10 μL of propidium iodide (PI) in 195 μL of the binding buffer. The results were analyzed by the FlowJo 7.6.1 software (FlowJo, Ashland, OR, USA) before being presented.

### The transwell-based migration and invasion assays

The Transwell-based migration and invasion assays were conducted in the modified Boyden chambers (Transwell; Corning Inc. Lowell, MA, USA). Equal numbers cells were allowed for growth in each well of 12-well plates. After reaching 70-80% confluence, they were starved for 12 h in serum-free medium. The experimental cells (1 × 10^4^) were suspended in 0.5 ml medium containing 5% FBS and seeded in the upper chamber of a Transwell. The cell-seeded Transwells were placed in each well of 24-well plates containing 1 ml of complete medium (i.e., the lower chamber), and then cultured for 24 h in the incubator at 37°C with 5% CO2. Of note, the bottom of upper Transwell was pre-coated by matrigel basement matrix (BD, Biosciences, USA), before the cells were placed in the invasion assay. The remaining cells in the upper chamber were removed, and the cells attached to the lower surface of the Transwell membranes were fixed with 4% paraformaldehyde (AR10669, BOSTER) and stained with 1% crystal violet reagent (Sigma) before being counted.

### Statistical analysis

Statistical significance of changes in reporter activity and other gene expression was determined using either the Student’s t-test or Multiple Analysis of Variations (MANOVA). The resulting data are shown as a fold change (mean ± S.D), each of which represents at least 3 independent experiments that were each performed triplicate.

## Author contributions

S.H. performed most of the experiments except indicated elsewhere, made all figures and wrote the manuscript draft. J.F. performed Construction of relevant plasmid used in this study. M.W. performed the statistical analysis of transcriptome data. R.W. and K.L. participated in the detection of intracellular ROS and transmission electron microscopy. Z.Z. participated in drawing of some pictures and revising of manuscript draft. Y.Z. designed and supervised this study, analyzed all the data, helped to prepare all figures with cartoons, wrote and revised the paper.

## Acknowledgments

We are greatly thankful to Dr. Yonggang Ren (North Sichuan Medical College, Sichuan, China), Lu Qiu (Zhengzhou University, Henan, China) for their involvement in establishing the indicated cell lines used in this study. We also thank to the members of Prof. Zhang’s laboratory (at Chongqing University, China) for giving their invaluable help with this work. Notably, this study was funded by the National Natural Science Foundation of China (NSFC, with a key program 91429305 and additional two projects 81872336 and 82073079) awarded to Prof. Yiguo Zhang.

## References

1. Kong, H., and Chandel, N. S. (2018) Regulation of redox balance in cancer and T cells. The Journal of biological chemistry 293, 7499–7507

2. Handy, D. E., and Loscalzo, J. (2012) Redox regulation of mitochondrial function. Antioxidants & redox signaling 16, 1323–1367

3. Murphy, M. P. (2009) How mitochondria produce reactive oxygen species. The Biochemical journal 417, 1–13

4. Holmstrom, K. M., and Finkel, T. (2014) Cellular mechanisms and physiological consequences of redox-dependent signalling. Nature reviews. Molecular cell biology 15, 411–421

5. Harris, I. S., and DeNicola, G. M. (2020) The Complex Interplay between Antioxidants and ROS in Cancer. Trends in cell biology 30, 440–451

6. Gozzelino, R., Jeney, V., and Soares, M. P. (2010) Mechanisms of cell protection by heme oxygenase-1. Annual review of pharmacology and toxicology 50, 323–354

7. Sies, H. (2017) Hydrogen peroxide as a central redox signaling molecule in physiological oxidative stress: Oxidative eustress. Redox Biol 11, 613–619

8. Sies, H. (2021) Oxidative eustress: On constant alert for redox homeostasis. Redox Biol 41, 101867

9. Yuan, J., Zhang, S., and Zhang, Y. (2018) Nrf1 is paved as a new strategic avenue to prevent and treat cancer, neurodegenerative and other diseases. Toxicol Appl Pharmacol 360, 273–283

10. Yang, S., and Lian, G. (2020) ROS and diseases: role in metabolism and energy supply. Molecular and cellular biochemistry 467, 1–12

11. Gorrini, C., Harris, I. S., and Mak, T. W. (2013) Modulation of oxidative stress as an anticancer strategy. Nature reviews. Drug discovery 12, 931–947

12. Weinberg, F., Hamanaka, R., Wheaton, W. W., Weinberg, S., Joseph, J., Lopez, M., Kalyanaraman, B., Mutlu, G. M., Budinger, G. R., and Chandel, N. S. (2010) Mitochondrial metabolism and ROS generation are essential for Kras-mediated tumorigenicity. Proceedings of the National Academy of Sciences of the United States of America 107, 8788–8793

13. Morgan, M. J., and Liu, Z. G. (2011) Crosstalk of reactive oxygen species and NF-kappaB signaling. Cell research 21, 103–115

14. Zhong, W., Qian, K., Xiong, J., Ma, K., Wang, A., and Zou, Y. (2016) Curcumin alleviates lipopolysaccharide induced sepsis and liver failure by suppression of oxidative stress-related inflammation via PI3K/AKT and NF-kappaB related signaling. Biomedicine & pharmacotherapy = Biomedecine & pharmacotherapie 83, 302–313

15. Chatterjee, R., and Chatterjee, J. (2020) ROS and oncogenesis with special reference to EMT and stemness. European journal of cell biology 99, 151073

16. Dewhirst, M. W., Cao, Y., and Moeller, B. (2008) Cycling hypoxia and free radicals regulate angiogenesis and radiotherapy response. Nature reviews. Cancer 8, 425–437

17. Sosa, V., Moline, T., Somoza, R., Paciucci, R., Kondoh, H., and Me, L. L. (2013) Oxidative stress and cancer: an overview. Ageing research reviews 12, 376–390

18. Nel, A., Xia, T., Madler, L., and Li, N. (2006) Toxic potential of materials at the nanolevel. Science 311, 622–627

19. Yamamoto, M., Kensler, T. W., and Motohashi, H. (2018) The KEAP1-NRF2 System: a Thiol-Based Sensor-Effector Apparatus for Maintaining Redox Homeostasis. Physiological reviews 98, 1169–1203

20. Zhang, Y., and Xiang, Y. (2016) Molecular and cellular basis for the unique functioning of Nrf1, an indispensable transcription factor for maintaining cell homoeostasis and organ integrity. The Biochemical journal 473, 961–1000

21. Xiang, Y., Wang, M., Hu, S., Qiu, L., Yang, F., Zhang, Z., Yu, S., Pi, J., and Zhang, Y. (2018) Mechanisms controlling the multistage post-translational processing of endogenous Nrf1alpha/TCF11 proteins to yield distinct isoforms within the coupled positive and negative feedback circuits. Toxicology and applied pharmacology 360, 212–235

22. Zhang, Y., Li, S., Xiang, Y., Qiu, L., Zhao, H., and Hayes, J. D. (2015) The selective post-translational processing of transcription factor Nrf1 yields distinct isoforms that dictate its ability to differentially regulate gene expression. Scientific reports 5

23. Bellezza, I., Giambanco, I., Minelli, A., and Donato, R. (2018) Nrf2-Keap1 signaling in oxidative and reductive stress. Biochimica et biophysica acta. Molecular cell research 1865, 721–733

24. Lu, S. C. (2013) Glutathione synthesis. Biochimica et biophysica acta 1830, 3143–3153

25. Chartoumpekis, D. V., Wakabayashi, N., and Kensler, T. W. (2015) Keap1/Nrf2 pathway in the frontiers of cancer and non-cancer cell metabolism. Biochemical Society transactions 43, 639–644

26. Nioi, P., and Hayes, J. D. (2004) Contribution of NAD(P)H:quinone oxidoreductase 1 to protection against carcinogenesis, and regulation of its gene by the Nrf2 basic-region leucine zipper and the arylhydrocarbon receptor basic helix-loop-helix transcription factors. Mutation research 555, 149–171

27. Zhang, Y., Nicholatos, J., Dreier, J. R., Ricoult, S. J., Widenmaier, S. B., Hotamisligil, G. S., Kwiatkowski, D. J., and Manning, B. D. (2014) Coordinated regulation of protein synthesis and degradation by mTORC1. Nature 513, 440–443

28. Xue, P., Hou, Y., Zuo, Z., Wang, Z., Ren, S., Dong, J., Fu, J., Wang, H., Andersen, M. E., Zhang, Q., Xu, Y., and Pi, J. (2020) Long isoforms of NRF1 negatively regulate adipogenesis via suppression of PPARgamma expression. Redox biology 30, 101414

29. Hirotsu, Y., Hataya, N., Katsuoka, F., and Yamamoto, M. (2012) NF-E2-related factor 1 (Nrf1) serves as a novel regulator of hepatic lipid metabolism through regulation of the Lipin1 and PGC-1beta genes. Molecular and cellular biology 32, 2760–2770

30. Widenmaier, S. B., Snyder, N. A., Nguyen, T. B., Arduini, A., Lee, G. Y., Arruda, A. P., Saksi, J., Bartelt, A., and Hotamisligil, G. S. (2017) NRF1 Is an ER Membrane Sensor that Is Central to Cholesterol Homeostasis. Cell 171, 1094–1109 e1015

31. Zheng, H., Fu, J., Xue, P., Zhao, R., Dong, J., Liu, D., Yamamoto, M., Tong, Q., Teng, W., Qu, W., Zhang, Q., Andersen, M. E., and Pi, J. (2015) CNC-bZIP protein Nrf1-dependent regulation of glucose-stimulated insulin secretion. Antioxidants & redox signaling 22, 819–831

32. Hirotsu, Y., Higashi, C., Fukutomi, T., Katsuoka, F., Tsujita, T., Yagishita, Y., Matsuyama, Y., Motohashi, H., Uruno, A., and Yamamoto, M. (2014) Transcription factor NF-E2-related factor 1 impairs glucose metabolism in mice. Genes Cells 19, 650–665

33. Rojo de la Vega, M., Chapman, E., and Zhang, D. D. (2018) NRF2 and the Hallmarks of Cancer. Cancer cell 34, 21–43

34. Ren, Y., Qiu, L., Lu, F., Ru, X., Li, S., Xiang, Y., Yu, S., and Zhang, Y. (2016) TALENs-directed knockout of the full-length transcription factor Nrf1alpha that represses malignant behaviour of human hepatocellular carcinoma (HepG2) cells. Scientific reports 6, 23775

35. Xu, Z., Chen, L., Leung, L., Yen, T. S., Lee, C., and Chan, J. Y. (2005) Liver-specific inactivation of the Nrf1 gene in adult mouse leads to nonalcoholic steatohepatitis and hepatic neoplasia. Proceedings of the National Academy of Sciences of the United States of America 102, 4120–4125

36. Tsujita, T., Peirce, V., Baird, L., Matsuyama, Y., Takaku, M., Walsh, S. V., Griffin, J. L., Uruno, A., Yamamoto, M., and Hayes, J. D. (2014) Transcription factor Nrf1 negatively regulates the cystine/glutamate transporter and lipid-metabolizing enzymes. Molecular and cellular biology 34, 3800–3816

37. Qiu, L., Wang, M., Hu, S., Ru, X., Ren, Y., Zhang, Z., Yu, S., and Zhang, Y. (2018) Oncogenic Activation of Nrf2, Though as a Master Antioxidant Transcription Factor, Liberated by Specific Knockout of the Full-Length Nrf1alpha that Acts as a Dominant Tumor Repressor. Cancers 10

38. Leung, L., Kwong, M., Hou, S., Lee, C., and Chan, J. Y. (2003) Deficiency of the Nrf1 and Nrf2 transcription factors results in early embryonic lethality and severe oxidative stress. The Journal of biological chemistry 278, 48021–48029

39. Zucker, S. N., Fink, E. E., Bagati, A., Mannava, S., Bianchi-Smiraglia, A., Bogner, P. N., Wawrzyniak, J. A., Foley, C., Leonova, K. I., Grimm, M. J., Moparthy, K., Ionov, Y., Wang, J., Liu, S., Sexton, S., Kandel, E. S., Bakin, A. V., Zhang, Y., Kaminski, N., Segal, B. H., and Nikiforov, M. A. (2014) Nrf2 amplifies oxidative stress via induction of Klf9. Mol Cell 53, 916–928

40. Bansal, A., and Simon, M. C. (2018) Glutathione metabolism in cancer progression and treatment resistance. The Journal of cell biology 217, 2291–2298

41. Budanov, A. V., Lee, J. H., and Karin, M. (2010) Stressin’ Sestrins take an aging fight. EMBO molecular medicine 2, 388–400

42. Zhu, Y. P., Zheng, Z., Xiang, Y., and Zhang, Y. (2020) Glucose Starvation-Induced Rapid Death of Nrf1alpha-Deficient, but Not Nrf2-Deficient, Hepatoma Cells Results from Its Fatal Defects in the Redox Metabolism Reprogramming. Oxidative medicine and cellular longevity 2020, 4959821

43. Li, N., Ragheb, K., Lawler, G., Sturgis, J., Rajwa, B., Melendez, J. A., and Robinson, J. P. (2003) Mitochondrial complex I inhibitor rotenone induces apoptosis through enhancing mitochondrial reactive oxygen species production. The Journal of biological chemistry 278, 8516–8525

44. Youle, R. J., and van der Bliek, A. M. (2012) Mitochondrial fission, fusion, and stress. Science 337, 1062–1065

45. Zhang, S., Deng, Y., Xiang, Y., Hu, S., Qiu, L., and Zhang, Y. (2020) Synergism and Antagonism of Two Distinct, but Confused, Nrf1 Factors in Integral Regulation of the Nuclear-to-Mitochondrial Respiratory and Antioxidant Transcription Networks. Oxid Med Cell Longev 2020, 5097109

46. Zhu, Y. P., Xiang, Y., L’Honore, A., Montarras, D., Buckingham, M., and Zhang, Y. (2020) Commentary on Distinct, but Previously Confused, Nrf1 Transcription Factors and Their Functions in Redox Regulation. Dev Cell 53, 377–378

47. Di, W., Lv, J., Jiang, S., Lu, C., Yang, Z., Ma, Z., Hu, W., Yang, Y., and Xu, B. (2018) PGC-1: The Energetic Regulator in Cardiac Metabolism. Current issues in molecular biology 28, 29–46

48. Wufuer, R., Fan, Z., Liu, K., and Zhang, Y. (2021) Differential Yet Integral Contributions of Nrf1 and Nrf2 in the Human HepG2 Cells on Antioxidant Cytoprotective Response against Tert-Butylhydroquinone as a Pro-Oxidative Stressor. Antioxidants 10

49. Shadel, G. S., and Horvath, T. L. (2015) Mitochondrial ROS signaling in organismal homeostasis. Cell 163, 560–569

50. Breen, G. A., and Scheffler, I. E. (1979) Respiration-deficient Chinese hamster cell mutants: biochemical characterization. Somatic cell genetics 5, 441–451

51. Baffy, G. (2010) Uncoupling protein-2 and cancer. Mitochondrion 10, 243–252

52. Bartelt, A., Widenmaier, S. B., Schlein, C., Johann, K., Goncalves, R. L. S., Eguchi, K., Fischer, A. W., Parlakgul, G., Snyder, N. A., Nguyen, T. B., Bruns, O. T., Franke, D., Bawendi, M. G., Lynes, M. D., Leiria, L. O., Tseng, Y. H., Inouye, K. E., Arruda, A. P., and Hotamisligil, G. S. (2018) Brown adipose tissue thermogenic adaptation requires Nrf1-mediated proteasomal activity. Nature medicine 24, 292–303

53. Fink, B. D., Hong, Y. S., Mathahs, M. M., Scholz, T. D., Dillon, J. S., and Sivitz, W. I. (2002) UCP2-dependent proton leak in isolated mammalian mitochondria. The Journal of biological chemistry 277, 3918–3925

54. Shpilka, T., and Haynes, C. M. (2018) The mitochondrial UPR: mechanisms, physiological functions and implications in ageing. Nature reviews. Molecular cell biology 19, 109–120

55. Li., L., Chen., Y., Chenzhao., C., Fu., S., Xu., Q., and Zhao., J. (2018) Glucose negatively affects Nrf2/SKN-1-mediated innate immunity in C. elegans. Aging 10, 3089–3103

56. Haynes, C. M., Yang, Y., Blais, S. P., Neubert, T. A., and Ron, D. (2010) The matrix peptide exporter HAF-1 signals a mitochondrial UPR by activating the transcription factor ZC376.7 in C. elegans. Molecular cell 37, 529–540

57. Fiorese, C. J., Schulz, A. M., Lin, Y. F., Rosin, N., Pellegrino, M. W., and Haynes, C. M. (2016) The Transcription Factor ATF5 Mediates a Mammalian Mitochondrial UPR. Current biology: CB 26, 2037–2043

58. Michel, S., Canonne, M., Arnould, T., and Renard, P. (2015) Inhibition of mitochondrial genome expression triggers the activation of CHOP-10 by a cell signaling dependent on the integrated stress response but not the mitochondrial unfolded protein response. Mitochondrion 21, 58–68

59. Quiros, P. M., Prado, M. A., Zamboni, N., D’Amico, D., Williams, R. W., Finley, D., Gygi, S. P., and Auwerx, J. (2017) Multi-omics analysis identifies ATF4 as a key regulator of the mitochondrial stress response in mammals. The Journal of cell biology 216, 2027–2045

60. Zhu, Y. P., Zheng, Z., Hu, S., Ru, X., Fan, Z., Qiu, L., and Zhang, Y. (2019) Unification of Opposites between Two Antioxidant Transcription Factors Nrf1 and Nrf2 in Mediating Distinct Cellular Responses to the Endoplasmic Reticulum Stressor Tunicamycin. Antioxidants 9

61. Fusakio, M. E., Willy, J. A., Wang, Y., Mirek, E. T., Al Baghdadi, R. J., Adams, C. M., Anthony, T. G., and Wek, R. C. (2016) Transcription factor ATF4 directs basal and stress-induced gene expression in the unfolded protein response and cholesterol metabolism in the liver. Molecular biology of the cell 27, 1536–1551

62. Tremblay, B. P., and Haynes, C. M. (2020) Mitochondrial distress call moves to the cytosol to trigger a response to stress. Nature 579, 348–349

63. Fessler, E., Eckl, E. M., Schmitt, S., Mancilla, I. A., Meyer-Bender, M. F., Hanf, M., Philippou-Massier, J., Krebs, S., Zischka, H., and Jae, L. T. (2020) A pathway coordinated by DELE1 relays mitochondrial stress to the cytosol. Nature 579, 433–437

64. Berry, B. J., Nieves, T. O., and Wojtovich, A. P. (2021) Decreased Mitochondrial Membrane Potential Activates the Mitochondrial Unfolded Protein Response. MicroPubl Biol 2021

65. Barrallo-Gimeno, A., and Nieto, M. A. (2005) The Snail genes as inducers of cell movement and survival: implications in development and cancer. Development 132, 3151–3161

66. Wagner, E. F., and Nebreda, A. R. (2009) Signal integration by JNK and p38 MAPK pathways in cancer development. Nature reviews. Cancer 9, 537–549

67. Moloney, J. N., and Cotter, T. G. (2018) ROS signalling in the biology of cancer. Seminars in cell & developmental biology 80, 50–64

68. Mitsuishi, Y., Taguchi, K., Kawatani, Y., Shibata, T., Nukiwa, T., Aburatani, H., Yamamoto, M., and Motohashi, H. (2012) Nrf2 redirects glucose and glutamine into anabolic pathways in metabolic reprogramming. Cancer cell 22, 66–79

69. Zhou, Y., Duan, S., Zhou, Y., Yu, S., Wu, J., Wu, X., Zhao, J., and Zhao, Y. (2015) Sulfiredoxin-1 attenuates oxidative stress via Nrf2/ARE pathway and 2-Cys Prdxs after oxygen-glucose deprivation in astrocytes. Journal of molecular neuroscience: MN 55, 941–950

70. Lee, Y. K., Woo, H. G., and Yoon, G. (2015) Mitochondrial defect-responsive gene signature in livercancer progression. BMB reports 48, 597–598

71. Lee, Y. K., Kwon, S. M., Lee, E. B., Kim, G. H., Min, S., Hong, S. M., Wang, H. J., Lee, D. M., Choi, K. S., Park, T. J., and Yoon, G. (2020) Mitochondrial Respiratory Defect Enhances Hepatoma Cell Invasiveness via STAT3/NFE2L1/STX12 Axis. Cancers 12

72. Kukat, C., and Larsson, N. G. (2013) mtDNA makes a U-turn for the mitochondrial nucleoid. Trends in cell biology 23, 457–463

73. Donnelly., M., and Scheffler., I. E. (1976) Energy metabolism in respiration-deficient and wild type chinese hamster fibroblasts in culture. Journal of Cellular Physiology 89, 39–51

74. Brand, M. D., Affourtit, C., Esteves, T. C., Green, K., Lambert, A. J., Miwa, S., Pakay, J. L., and Parker, N. (2004) Mitochondrial superoxide: production, biological effects, and activation of uncoupling proteins. Free radical biology & medicine 37, 755–767

75. Starkov, A. A. (1997) “Mild” uncoupling of mitochondria. Biosci Rep 17, 273–279

76. Rojo de la Vega, M., Chapman, E., and Zhang, D. D. (2018) NRF2 and the Hallmarks of Cancer. Cancer Cell

77. Wang, M., Ren, Y., Hu, S., Liu, K., Qiu, L., and Zhang, Y. (2021) TCF11 Has a Potent Tumor-Repressing Effect Than Its Prototypic Nrf1alpha by Definition of Both Similar Yet Different Regulatory Profiles, With a Striking Disparity From Nrf2. Frontiers in oncology 11, 707032

78. Hayes, J. D., Dinkova-Kostova, A. T., and Tew, K. D. (2020) Oxidative Stress in Cancer. Cancer Cell 38, 167–197

79. Ren, Y., Qiu, L., Lü, F., Ru, X., Li, S., Xiang, Y., Yu, S., and Zhang, Y. (2016) TALENs-directed knockout of the full-length transcription factor Nrf1a that represses malignant behaviour of human hepatocellular carcinoma (HepG2) cells Scientific Reports 7, Accepted for publication

