## supplementary materials for "Nrf1 is an indispensable redox-determining factor for mitochondrial homeostasis by integrating multi-hierarchical regulatory networks"

**Supplementary Materials** included the relevant experimental data as shown in eight figures below, in addition to TableS1, showing a list of all key reagents and resources used in this work.

### Figure S1

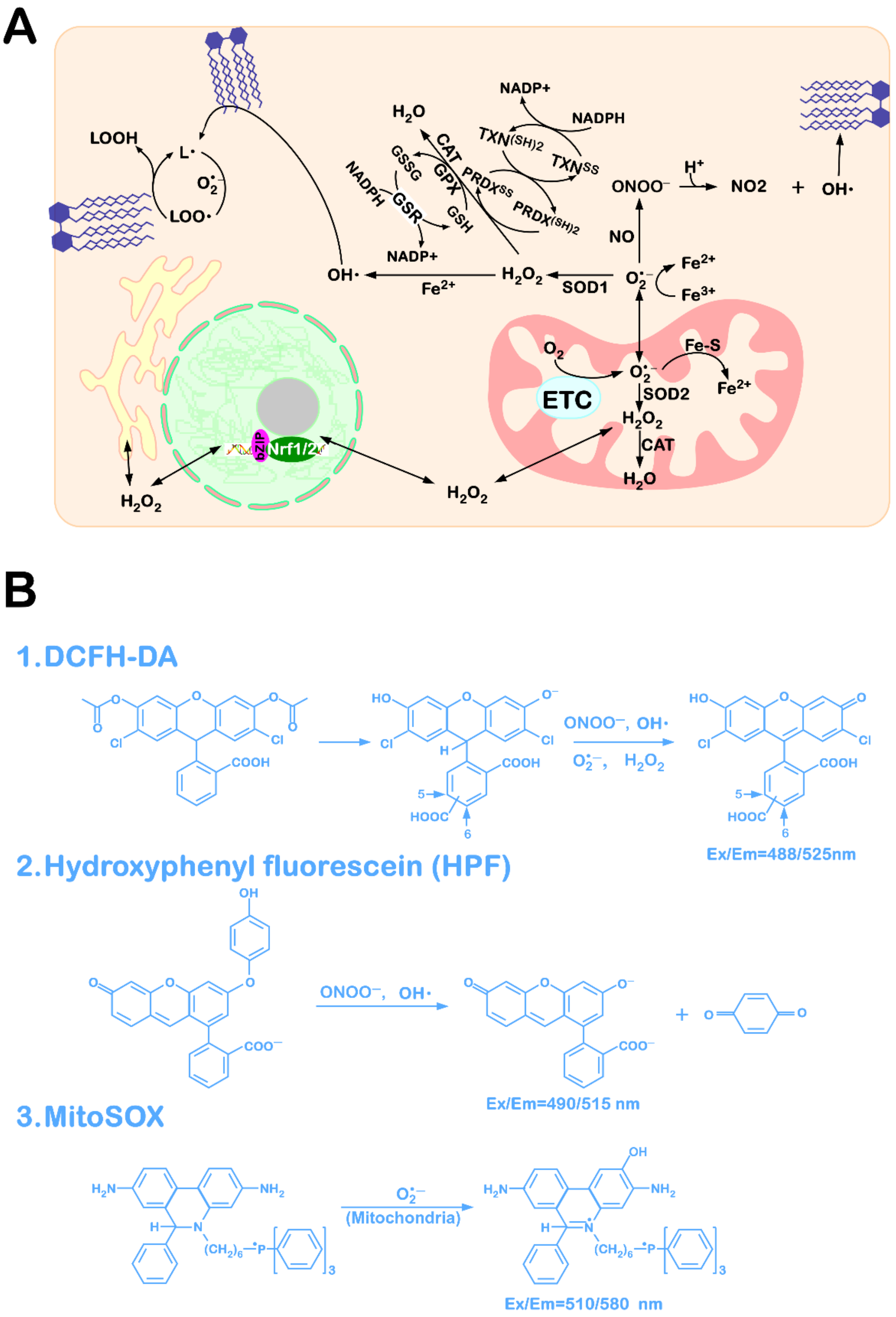


**Figure S1 Schematic representation for ROS generation and elimination and Schematic diagram of fluorescent probe**

(A) Schematic representation showing the process of intracellular ROS generation and elimination.

(B) Schematic diagram of fluorescent probe (DCFH-DA, Hydroxyphenyl fluorescein, MitiSOX) reacting with ROS to produce fluorescence.

### Figure S2

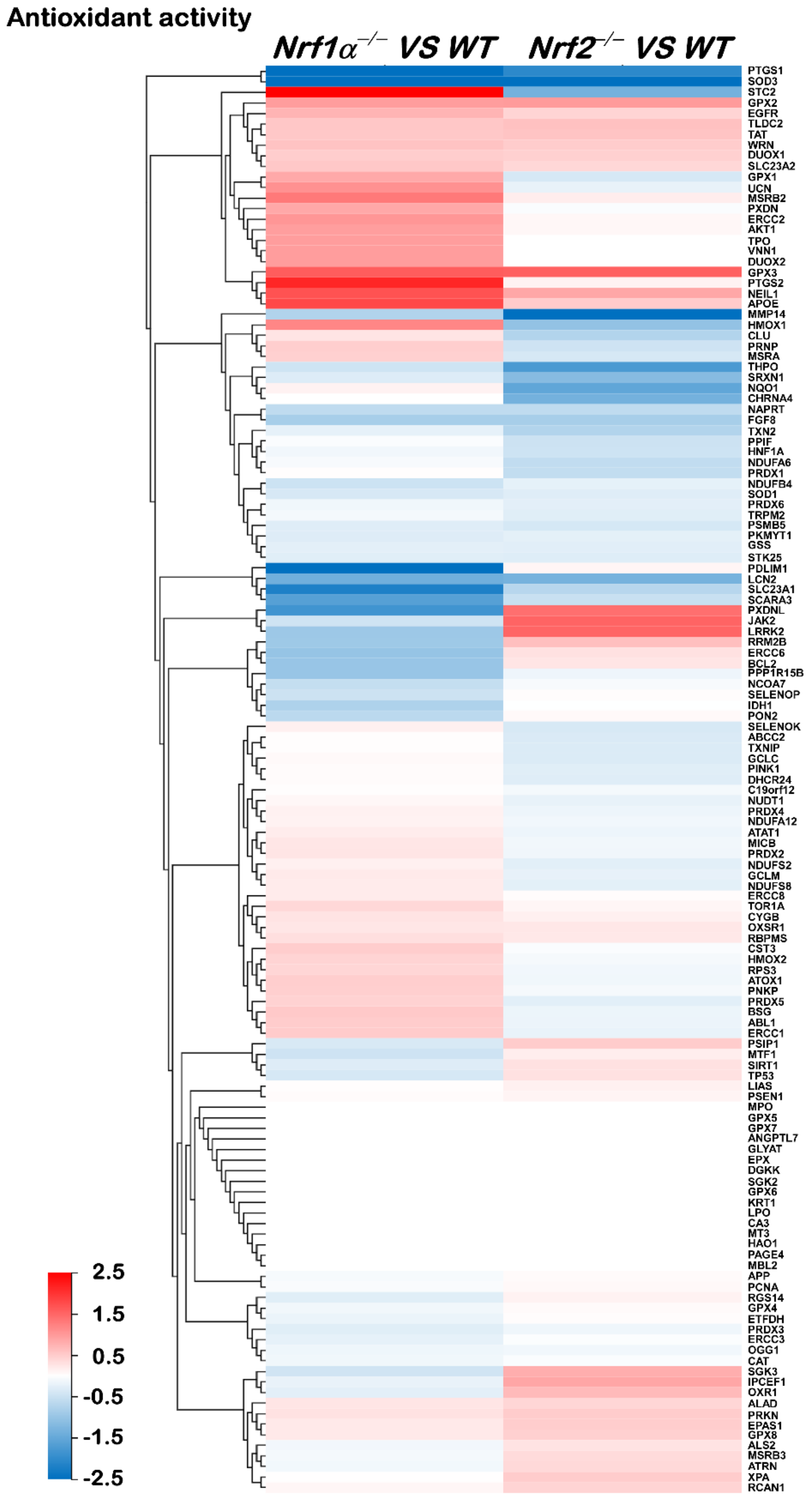


**Figure S2 The heatmap of the sequencing expression of genes with antioxidant activity in *Nrf1α*$^{-/-}$, *Nrf2*$^{-/-}$ cells compared with the *WT* cells**

The color of the nodes in the heatmap represents the value of $\log_2$ (fold change) as shown in the color bars, indicating the gene expression trend as compared with the wild type cells (upregulation or downregulation, were marked in red or blue, respectively).

### Figure S3

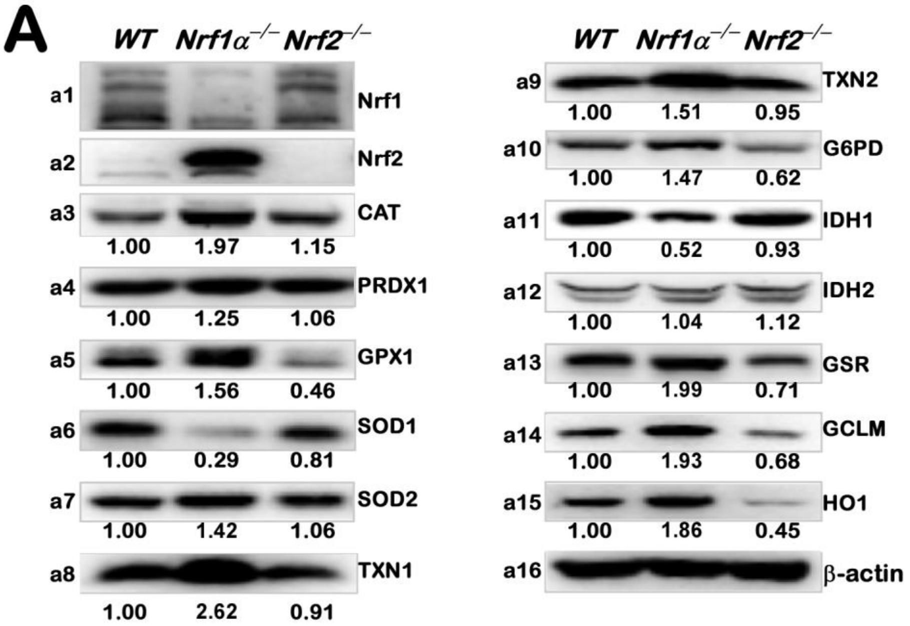


**Figure S3 The protein levels of antioxidant genes in *WT*, *Nrf1α*$^{-/-}$, *Nrf2*$^{-/-}$ cells**

The protein levels of Nrf1, Nrf2, CAT, PRDX1, GPX1, SOD1, SOD2, TXN1, TXN2, G6PD, IDH1, IDH2, GSR, GCLM, and HO1 in *WT*, *Nrf1α*$^{-/-}$ and *Nrf2*$^{-/-}$ cells were visualized by Western blotting. The intensity of all the immunoblots was calculated and shown on the bottom. The images without intensity of the immunoblots are shown in the main Figure 1N.

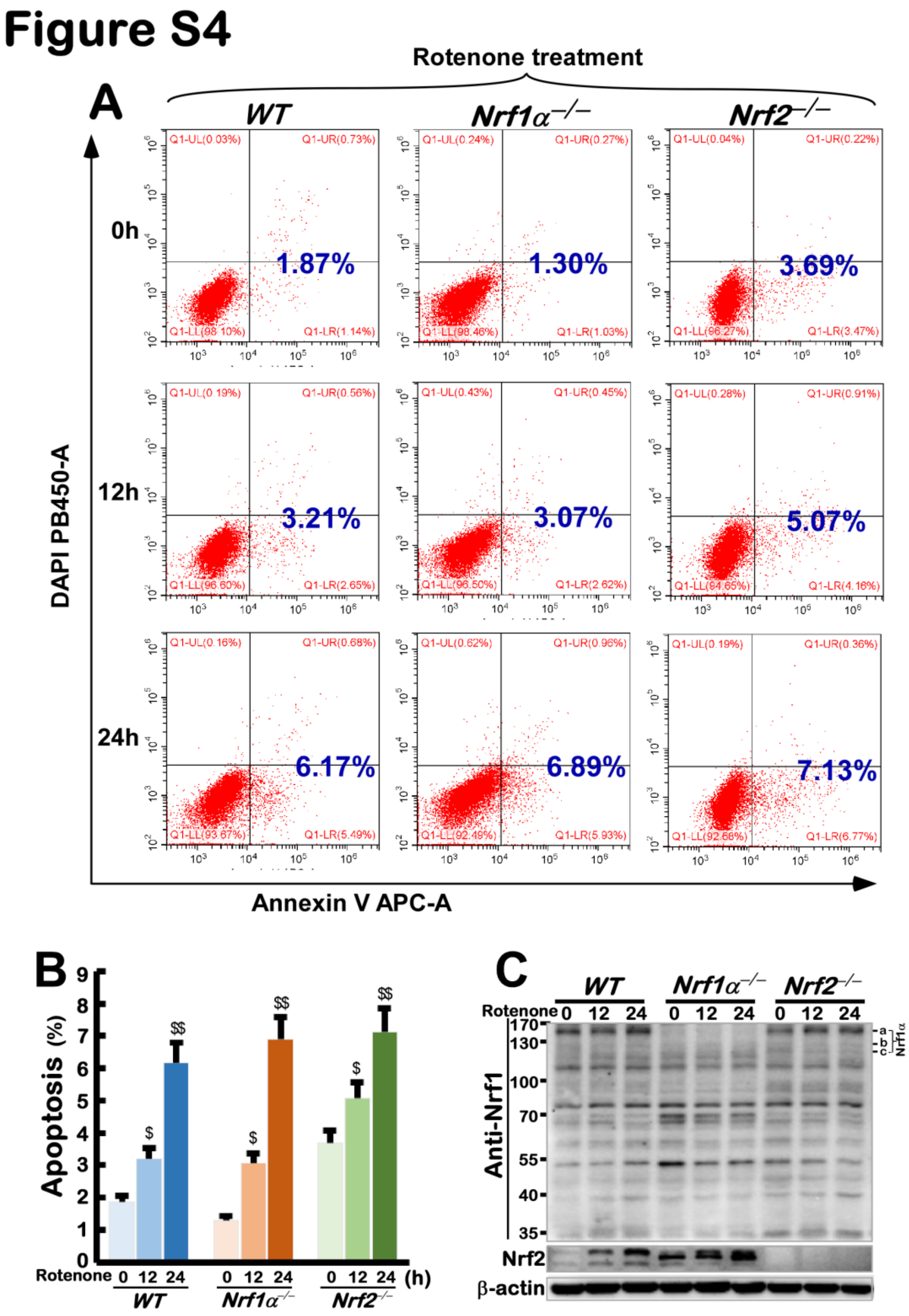


**Figure S4 The effect of rotenone on apoptosis in *WT*, *Nrf1α*$^{-/-}$, *Nrf2*$^{-/-}$ cells**

(A) The apoptosis was analyzed by flow cytometry in *WT*, *Nrf1α*$^{-/-}$ and *Nrf2*$^{-/-}$ cells, which were treated by Rotenone for 0h, 12h, or 24h and incubated with Annexin V-FITC and PI.

(B) The apoptosis analyzed by flow cytometry above were shown with the column charts. Significant statistical increases were indicated with \$, $p < 0.05$ and \$\$, $p < 0.01$.

(C) The protein levels of, Nrf1, Nrf2, were visualized by Western blotting in *WT*, *Nrf1α*$^{-/-}$ and *Nrf2*$^{-/-}$ cells, which had been treated with Rotenone for 0h, 12h or 24h.

### Figure S5

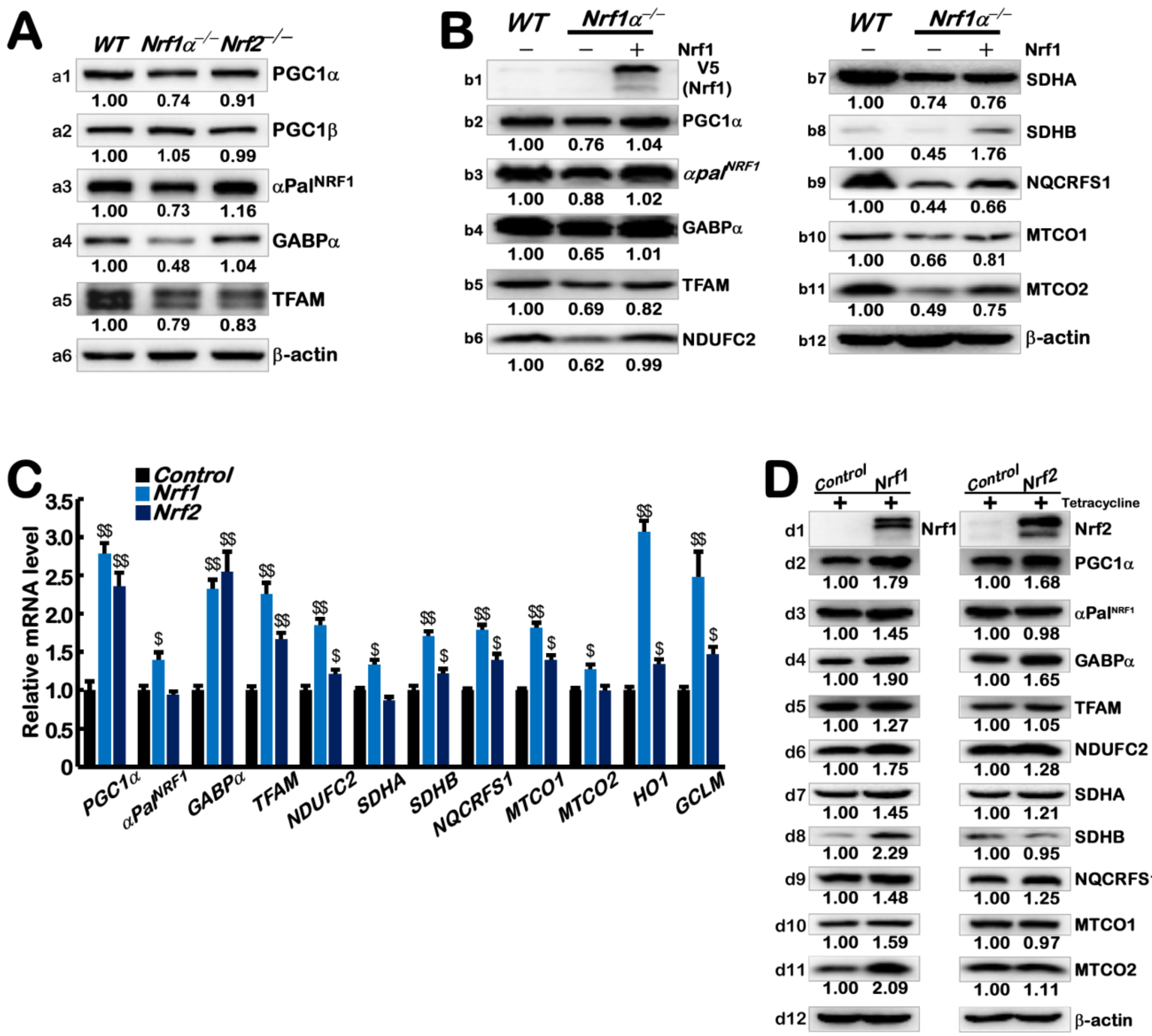


**Figure S5 The effect of overexpressing Nrf1/2 on PGC1α, PGC1β, αPal$^{NRF1}$, GABPα and other mitochondrial proteins**

(A) The protein levels of PGC1α, PGC1β, αPal$^{NRF1}$, GABPα and TFAM in *WT*, *Nrf1α*$^{-/-}$ and *Nrf2*$^{-/-}$ cells were visualized by Western blotting. The intensity of all the immunoblots was calculated and shown on the bottom. The images without intensity of the immunoblots are shown in the main Figure 3B.

(B) After Nrf1 was transfected into *Nrf1α*$^{-/-}$ cells, changed protein levels of PGC1α, αpal$^{NRF1}$, GABPα, TFAM, NDUFC2, SDHA, SDHB, UQCRFS1, MTCO1, MTCO2 were determined in *WT*, *Nrf1α*$^{-/-}$ and *Nrf2*$^{-/-}$ cells with Nrf1-restored. The intensity of all the immunoblots was calculated and shown on the bottom.

(C) The mRNA levels of *PGC1α, PGC1β, αPalNRF1, GABPα, TFAM, NDUFC2, SDHA, SDHB, NQCRFS1, MTCO1 and MTCO2* were determined by qPCR in Nrf1, Nrf2 and Control cells which were incubated with 1 mg/ml Tet for 24 h.

(D) The protein levels of PGC1α, PGC1β, αPalNRF1, GABPα, TFAM, NDUFC2, SDHA, SDHB, NQCRFS1, MTCO1 and MTCO2 were determined by Western blotting in Nrf1, Nrf2 and Control cells which were incubated with 1 mg/ml Tet for 24 h. The intensity of all the immunoblots was calculated and shown on the bottom.

### Figure S6

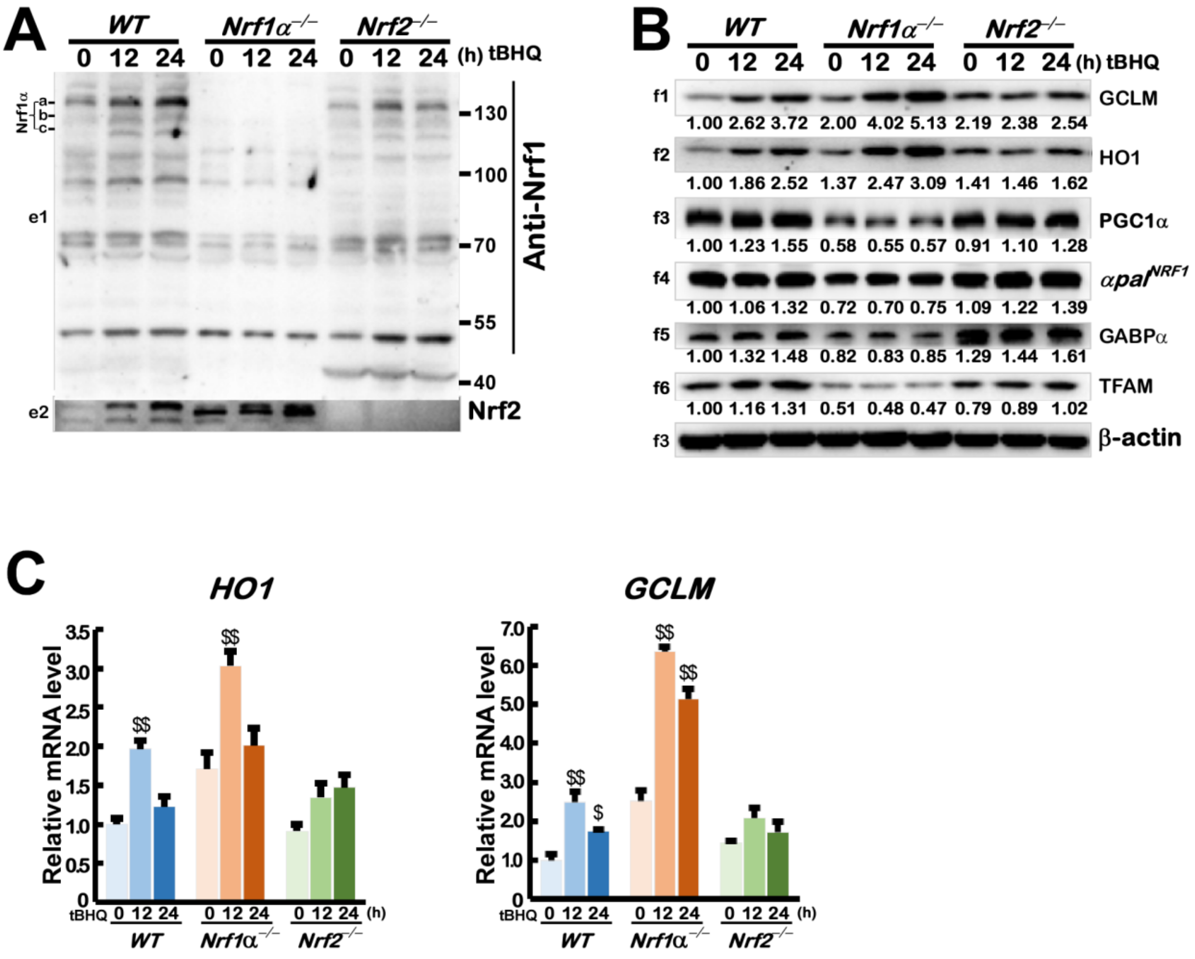


**Figure S6 Knockout of Nrf1α affects the activation of mitochondria-related genes by tbhQ**

(A) The protein levels of Nrf1 and Nrf2 were visualized by Western blotting in *WT*, *Nrf1α*$^{-/-}$ and *Nrf2*$^{-/-}$ cells, which had been treated with tbhQ for 0h, 12h or 24h.

(B) The protein levels of indicated genes above were visualized by Western blotting in *WT*, *Nrf1α*$^{-/-}$ and *Nrf2*$^{-/-}$ cells, which had been treated with tbhQ for 0h, 12h or 24h. The intensity of all the immunoblots was calculated and shown on the bottom.

(C) The mRNA levels of *HO1* and *GCLM* were determined by qPCR in *WT*, *Nrf1α*$^{-/-}$ and *Nrf2*$^{-/-}$ cells, which had been treated with tbhQ for 0h, 12h or 24h. The resulting data are shown as fold changes (mean ± SEM, n= 3 × 3) with significant increases ($, $p < 0.05$ and $$, $p < 0.01$) as compared to the cells without tbhQ treatment.

### Figure S7

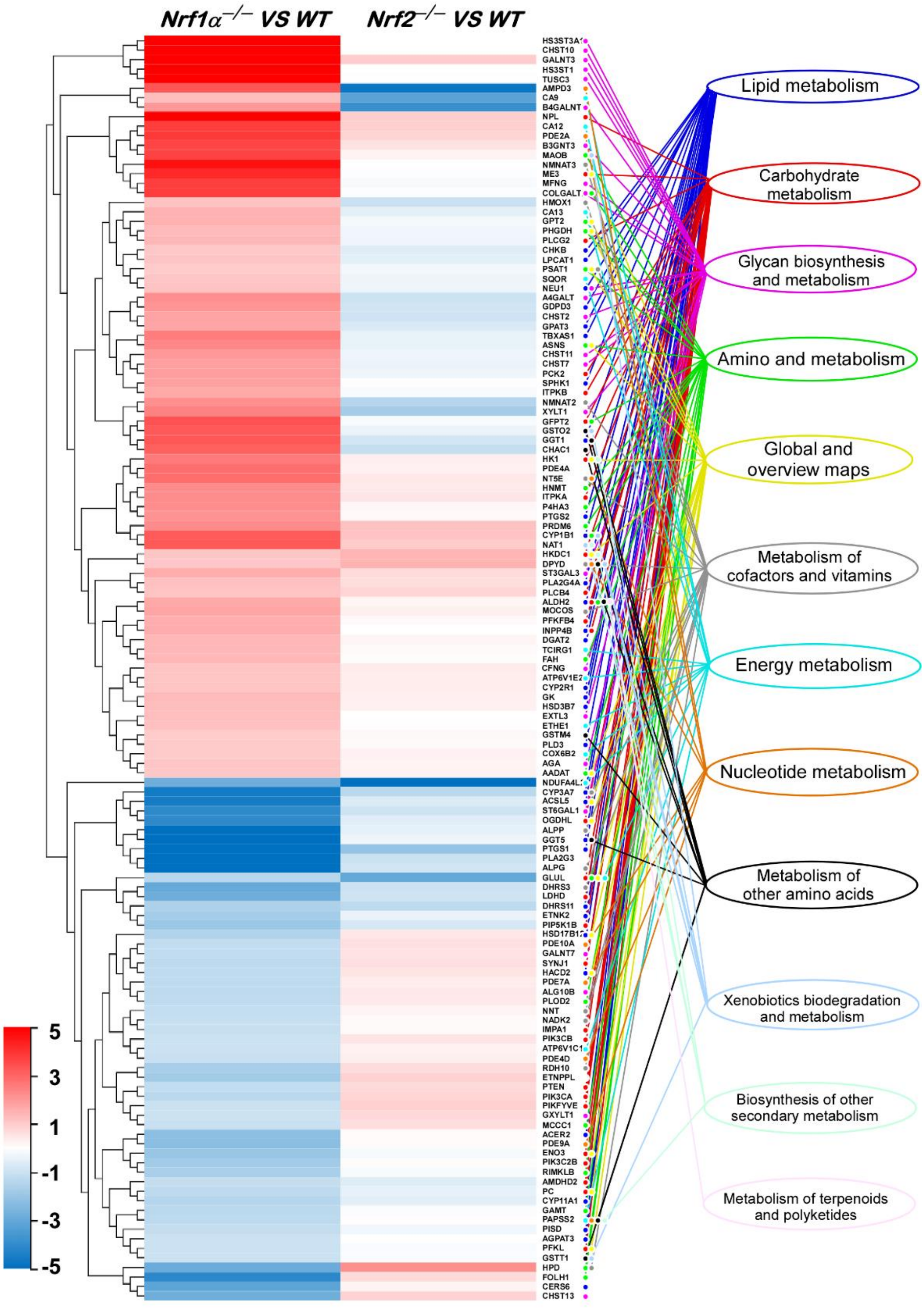


**Figure S7 The heatmap with hierarchical clustering of metabolism related DEGs in *Nrf1α*$^{-/-}$, *Nrf2*$^{-/-}$ cells**

The heatmap with hierarchical clustering of 133 metabolism related DEGs gathered by KEGG-pathway in *Nrf1α*$^{-/-}$ and *Nrf2*$^{-/-}$ cells. The color of the nodes in the heatmap represents the value of $\log_2$ (fold change) as shown in the color bars, indicating the gene expression trend as compared with the wild type cells (upregulation or downregulation, were marked in red or blue, respectively). Of note, the metabolic pathways to which the differential genes belong are indicated by different lines.

### Figure S8

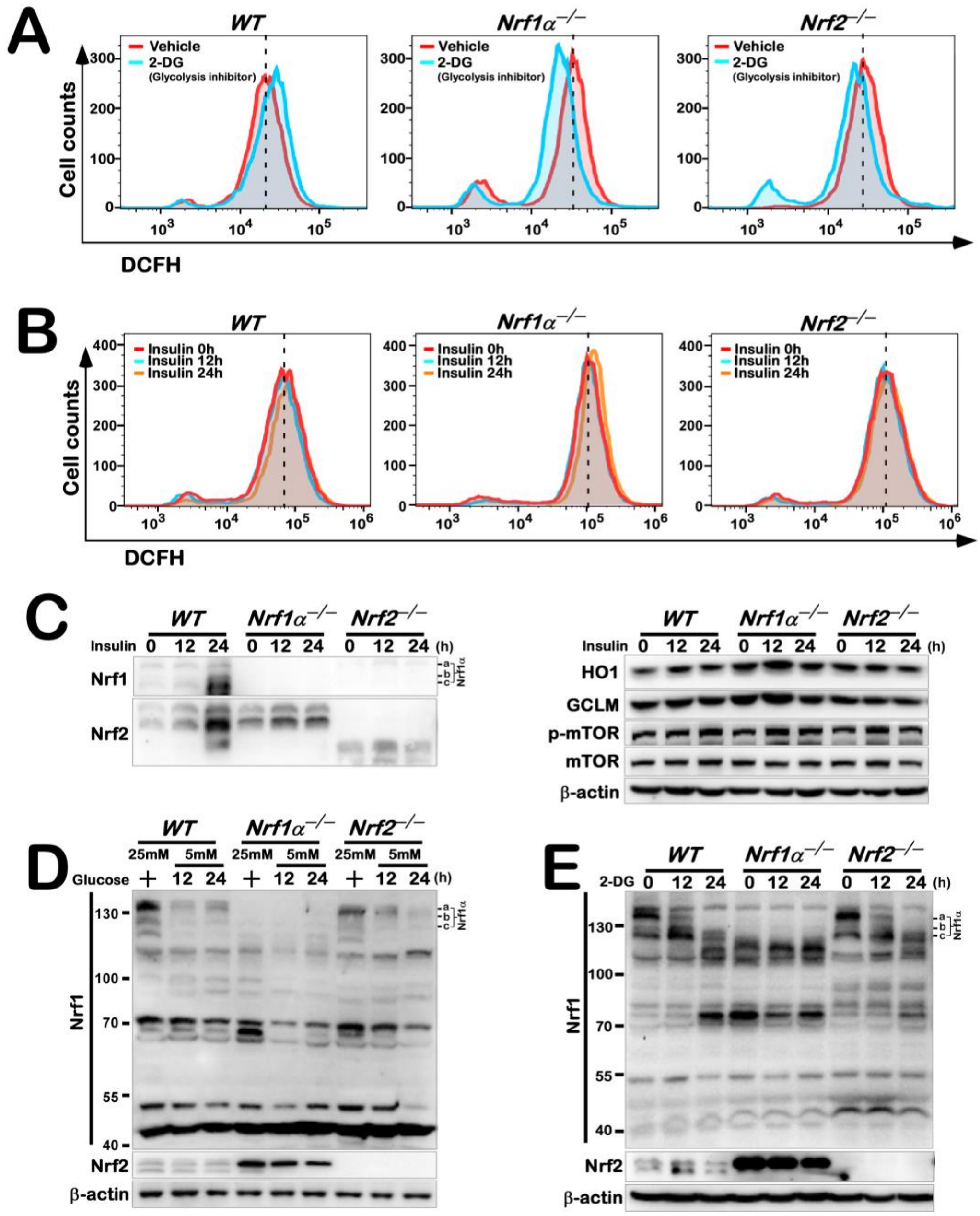


**Figure S8 Effects of glycolysis inhibitors 2-DG and activators insulin on intracellular ROS in *WT*, *Nrf1α*$^{-/-}$, *Nrf2*$^{-/-}$ cells**

(A) ROS levels were detected by flow cytometry in *WT*, *Nrf1α*$^{-/-}$ and *Nrf2*$^{-/-}$ cells, which were treated by 2-DG for 0h or 24h, and stained by DCFH.

(B) ROS levels were detected by flow cytometry in *WT*, *Nrf1α*$^{-/-}$ and *Nrf2*$^{-/-}$ cells, which were treated by insulin for 0h, 12h or 24h, and stained by DCFH.

(C) The protein levels of Nrf1, Nrf2, HO1, GCLM, p-mTOR, mTOR were visualized by Western blotting in *WT*, *Nrf1α*$^{-/-}$ and *Nrf2*$^{-/-}$ cells, which had been treated with insulin for 0h, 12h or 24h.

(D) The protein levels of Nrf1, Nrf2 were visualized by Western blotting in *WT*, *Nrf1α*$^{-/-}$ and *Nrf2*$^{-/-}$ cells, which were cultured by 5 mM glucose medium or 5 mM glucose medium for 12h and 24h.

(E) The protein levels of Nrf1, Nrf2 were visualized by Western blotting in *WT*, *Nrf1α*$^{-/-}$ and *Nrf2*$^{-/-}$ cells, which had been treated with 2-DG for 0h, 12h or 24h.

### Figure S9

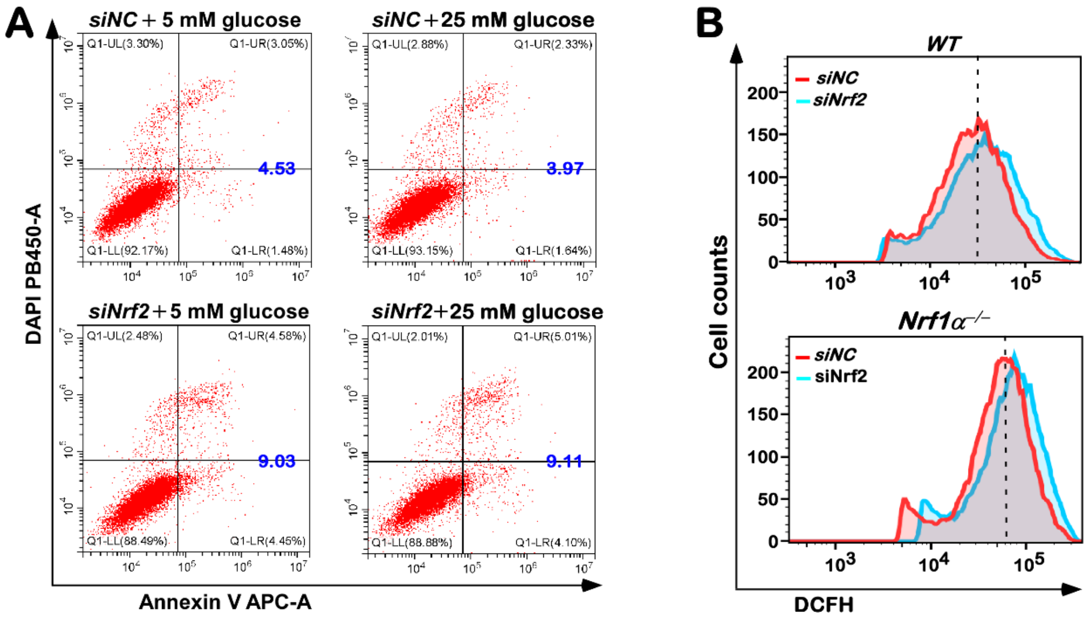


**Figure S9 The apoptosis of *Nrf1α*⁻/⁻ cells interfered by siNrf2 or scrambled siRNA and cultured by 25 mM glucose medium or 5 mM glucose medium**

(A) The *Nrf1α*⁻/⁻ cells were interfered by siNrf2 or scrambled siRNA and cultured by 25 mM glucose medium or 5 mM glucose medium for 24h, after incubated with Annexin V-FITC and PI, the apoptosis of cells was detected by flow cytometry.

(B) ROS levels detected by flow cytometry in *WT*, *Nrf1α*⁻/⁻ cells interfered by siNrf2 or scrambled siRNA.

### Figure S10

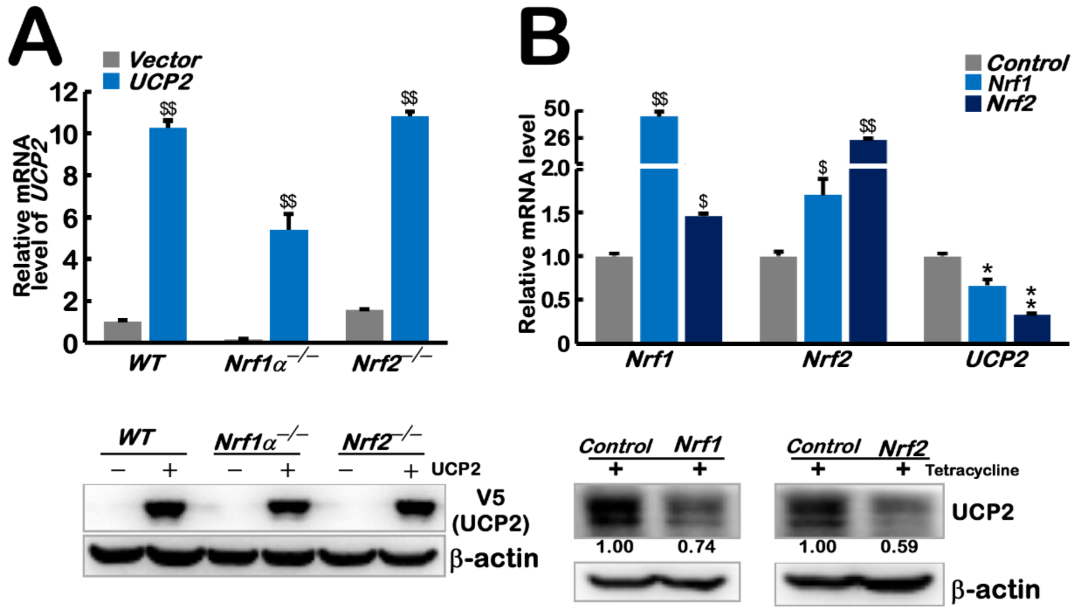


**Figure S10 Nrf1 and Nrf2 can inhibit the expression of UCP2**

(A) The mRNA (upper column) and protein (lower panel) levels of UCP2 were determined in *WT*, $Nrf1\alpha^{-/-}$ and $Nrf2^{-/-}$ cells, which were transfected with UCP2 or empty vector. The resulting data are shown as fold changes (mean ± SEM, n= 3 × 3; $$, $p < 0.01$).

(B) The mRNA (upper column) of Nrf1, Nrf2, UCP2 and protein (lower panel) levels of UCP2 were determined in in Nrf1a, Nrf2 and Control cells which were incubated with 1 mg/ml Tet for 24 h. The resulting data are shown as fold changes (mean ± SEM, n= 3 × 3; $, $p < 0.05$ and $$, $p < 0.01$; *, $p < 0.05$ and **, $p < 0.01$). The intensity of all the immunoblots was calculated and shown on the bottom.

### Figure S11

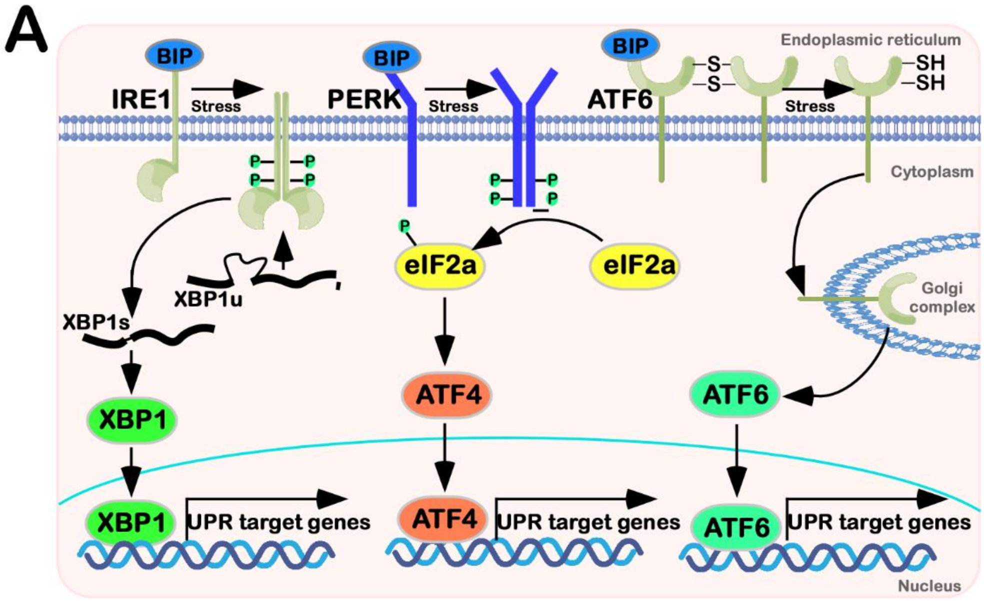


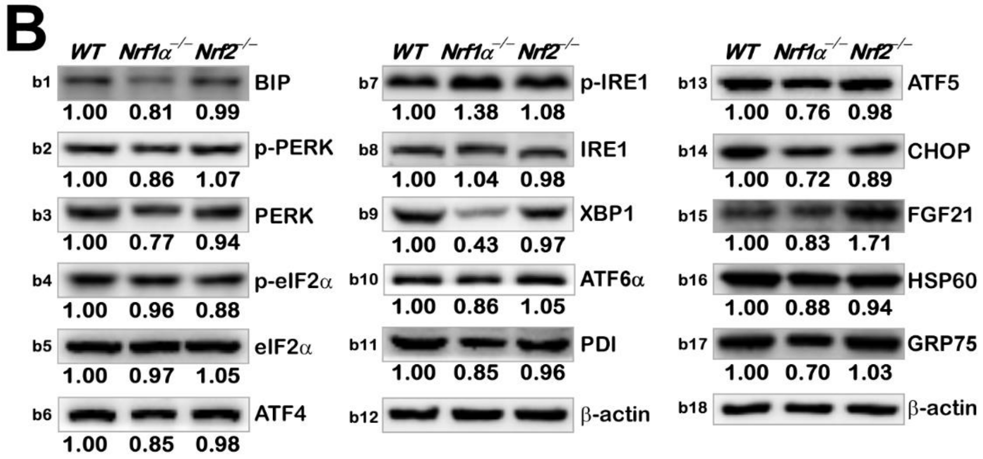


**Figure S11 The protein levels of endoplasmic reticulum stress-related genes in *WT*, *Nrf1α*$^{-/-}$, *Nrf2*$^{-/-}$ cells**

(A) A model of the three activated pathways in endoplasmic reticulum stress.

(B) The protein levels of BIP, p-PERK, PERK, p-eIF2a, eIF2a, ATF4, p-IRE1, IRE1, XBP1, ATF6, PDI, ATF5, CHOP, FGF21, HSP60, GRP75 were visualized by Western blotting in *WT*, *Nrf1α*$^{-/-}$ and *Nrf2*$^{-/-}$ cells. The intensity of all the immunoblots was calculated and shown on the bottom.

### Figure S12

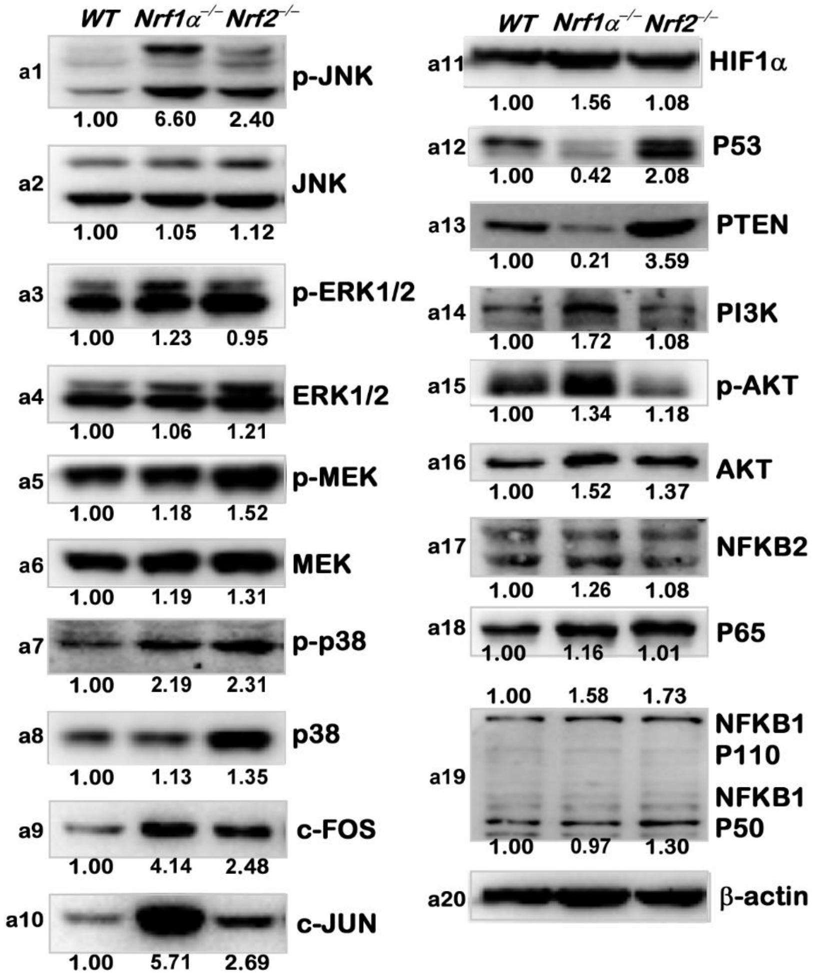


**Figure S12 The protein levels of different pathways in *WT*, *Nrf1α*$^{-/-}$, *Nrf2*$^{-/-}$ cells**

The protein levels of p-JNK, JNK, p-ERK1/2, ERK1/2, p-MEK1, MEK, p-p38, p38, HIF1α, PTEN, PI3K, p-AKT, NF-κB 2, NF-κB 1 (p105/p50), P65, c-FOS, c-JUN were visualized by Western blotting in *WT*, *Nrf1α*$^{-/-}$ and *Nrf2*$^{-/-}$ cells. The intensity of all the immunoblots was calculated and shown on the bottom. The images without intensity of the immunoblots are shown in the main Figure 7D.

**Table S1. A list of all key reagents and resources used in this work**

| Reagent or Resource | Source | Identifier |
|---|---|---|
| **antibodies** | | |
| CAT | BIMAKE | A5291 |
| PRDX1 | BIMAKE | A5448 |
| GPX1 | Abcam | ab108427 |
| SOD1 | R&D Systems | AF3418 |
| GCLC | Sangon Biotechnology | D123963 |
| GCLM | Abcam | ab126704 |
| GSR | Sangon Biotechnology | D220726 |
| TXN1 | BIMAKE | A5894 |
| TXN2 | ABclonal | A12591 |
| G6PD | ABclonal | A1537 |
| IDH1 | Sangon Biotechnology | D221821 |
| IDH2 | Sangon Biotechnology | D222530 |
| HO1 | Abcam | ab52947 |
| NDUFC2 | Abcam | ab192265 |
| SDHA | Sangon Biotechnology | D223104 |
| SDHB | Sangon Biotechnology | D262175 |
| UQCRFS1 | Abcam | ab191078 |
| MTCO1 | ABclonal | A17889 |
| MTCO2 | ABclonal | A11913 |
| OPA1 | Abcam | ab157457 |
| MFN1 | bimake | A5388 |
| MFN2 | bimake | A5754 |

| | | |
|---|---|---|
| DRP1 | Abcam | ab184247 |
| PGC1α | Sangon Biotechnology | D262041 |
| PGC1β | bimake | A5500 |
| αPal$^{NRF1}$ | Abcam | ab175932 |
| GABPα | Proteintech | 21542-1-AP |
| TFAM | Proteintech | 22586-1-AP |
| HK1 | Sangon Biotechnology | D221854 |
| HK2 | ABclonal | A0994 |
| GLUT1 | BIMAKE | A5514 |
| GLUT4 | | A11208 |
| LDHA | ABclonal | A7637 |
| PDH | ABclonal | A1146 |
| SREBP1 | Proteintech | 14088-1-AP |
| SREBP2 | Proteintech | 28212-1-AP |
| ACCα | Sangon Biotechnology | D155300 |
| FASN | Sangon Biotechnology | D262701 |
| SCD1 | Sangon Biotechnology | D162163 |
| CPT1A | Sangon Biotechnology | D120478 |
| UCP2 | Sangon Biotechnology | D262448 |
| ATF4 | BIMAKE | A5514 |
| ATF5 | Abcam | ab184923 |
| CHOP | Abcam | ab11419 |
| HSP60 | BIMAKE | A5629 |
| GRP75 | BIMAKE | A5420 |
| FGF21 | BIMAKE | A5710 |
| CDH1 | BIOSS | bs-10009R |

| | | |
|---|---|---|
| CDH2 | ABGENT | AO1377A |
| FN1 | BOSTER | BA1771 |
| ITGB4 | BOSTER | BA4486 |
| SNAI1 | BIOSS | bs-11961R |
| SNAIL2 | BOSTER | PB9439 |
| MMP9 | BIOSS | bs-4593R |
| JNK | Abcam | ab208035 |
| p-JNK | Abcam | ab124956 |
| ERK | BIMAKE | A5029 |
| p-ERK | BIMAKE | A5036 |
| MEK | BIMAKE | A5606 |
| p-MEK | BIMAKE | A5191 |
| P38 | BIMAKE | A5017 |
| p-P38 | Abcam | ab178876 |
| c-FOS | Abcam | ab134122 |
| c-JUN | Proteintech | 10024-2-AP |
| HIF1α | Abcam | ab51608 |
| P53 | Abcam | ab26 |
| PTEN | Abcam | ab32199 |
| PI3K | Sangon Biotechnology | D108795 |
| AKT1 | Abcam | ab32505 |
| NFKB1 | BIOSS | bs-1194R |
| NFKB2 | Sangon Biotechnology | D261024 |
| p-mTOR | Abcam | ab109268 |
| mTOR | Abcam | ab32028 |
| P65 | BIOSS | bs-0465R |

| | | |
|---|---|---|
| **Chemicals** | | |
| Rotenone | Sigma | R8875 |
| tert-Butylhydroquinone (TBHQ) | MCE | HY-100489 |
| 2-Deoxy-D-glucose (2-DG) | MCE | HY-13966 |
| Insilin | MCE | HY-P0035 |
| FCCP | MCE | HY-100410 |
| SP600125 | MCE | HY-12041 |
| **Experimental Models: Cell Lines** | | |
| HepG2 | Cell bank of the Chinese Academy of Sciences | TCHu72 |
| COS-1 | Cell bank of the Chinese Academy of Sciences | GNO28 |
| *Nrf1α*$^{-/-}$ | Zhang's laboratory | N/A |
| *Nrf2*$^{-/-}$ | Zhang's laboratory | N/A |
| Nrf1-expressing cells | Zhang's laboratory | N/A |
| Nrf2-expressing cells | Zhang's laboratory | N/A |
| **Oligonucleotides for qPCR** | | |
| CAT FW | Tsingke | CTCCAAATTACTACCCCAACAGC |
| CAT REV | Tsingke | ATCATTGGCAGTGTTGAATCTCC |
| PRDX1 FW | Tsingke | TTCACTGACAAACATGGGGAA |
| PRDX1 REV | Tsingke | CGCTCACTTCTGCTTGGAGA |
| PRDX3 FW | Tsingke | ATCTTGCCTGGATAAATACACC |
| PRDX3 REV | Tsingke | TGATCTTAGTGCAAGACCAGA |
| PRDX4 FW | Tsingke | GTGTGTCCAACTGAAATTATCGC |
| PRDX4 REV | Tsingke | ATTAATCCAGGCCAAATGGGT |
| PRDX5 FW | Tsingke | CATGGCCCCAATCAAGGTTCG |
| PRDX5 REV | Tsingke | CACTATGCCATCCTGTACCAC |
| PRDX6 FW | Tsingke | TAGGTTGCTATACCAATGGCTT |

| | | |
|---|---|---|
| PRDX6 REV | Tsingke | ATTTCAAACAGAACCAGCAG |
| GPX1 FW | Tsingke | CTTCTTGGACAATTGCGCCATG |
| GPX1 REV | Tsingke | CCGAGAAGGCATACACCGAC |
| GPX2 FW | Tsingke | CTACCTGAAGGACAAGCTCCC |
| GPX2 REV | Tsingke | CTTCTCAAAGTTCCAGGCCACA |
| SOD1 FW | Tsingke | TTGCATCATTGGCCGCACACT |
| SOD1 REV | Tsingke | ATTACACCACAAGCCAAACGACT |
| SOD2 FW | Tsingke | AATAGCTGGGATTACAGGTGCAT |
| SOD2 REV | Tsingke | CCAGACGGATCACTTGAGGTCAG |
| GCLC FW | Tsingke | TCAATGGGAAGGAAGGTGTGTT |
| GCLC REV | Tsingke | TTGTAGTCAGGATGGTTTGCGA |
| GCLM FW | Tsingke | TGCTGTGTGATGCCACCAGA |
| GCLM REV | Tsingke | CGCTTGAATGTCAGGAATGCTT |
| GLS1 FW | Tsingke | TTGGTCTTCCTGCAAAATCTGG |
| GLS1 REV | Tsingke | TCCCTTAACACTGTTGCCCAT |
| GSR FW | Tsingke | CACGAGTGATCCCAAGCCC |
| GSR REV | Tsingke | CAATGTAACCTGCACCAACAATG |
| SLC6A9 FW | Tsingke | GCCCTCACACTACTTCCCAT |
| SLC6A9 REV | Tsingke | GATCCACTCATTCCCCACCT |
| SLC7A11 FW | Tsingke | TAGACATGTTCCATTCGAGGT |
| SLC7A11 REV | Tsingke | TGTACACAACTTTGCTGGTC |
| TXN1 FW | Tsingke | CCTTGCAAAATGATCAAGCCTT |
| TXN1 REV | Tsingke | TGGTGGCTTCAAGCTTTTCCT |
| TXN2 FW | Tsingke | TGACCACACAGACCTCGCCAT |
| TXN2 REV | Tsingke | ATGCCCACAAACTTGTCCACCAC |
| TXNRD1 FW | Tsingke | GGCTCTATGCAGGTTCCACT |

| | | |
|---|---|---|
| TXNRD1 REV | Tsingke | TCAGAAAGGCCACAAGCACCA |
| TXNRD2 FW | Tsingke | AGATACCCCACGCACATCGAA |
| TXNRD2 REV | Tsingke | TTTTCCAGGGGATTCCTTCAGC |
| SRXN1 FW | Tsingke | GCTGTGTAACAAACCATCCCCAA |
| SRXN1 REV | Tsingke | CAGCCCAAATTGCCAACCCAT |
| G6PD FW | Tsingke | CGAGGCCGTCACCAAGAAC |
| G6PD REV | Tsingke | GTAGTGGTCGATGCGGTAGA |
| IDH1 FW | Tsingke | ACACCAAGTGACGGAACCCAA |
| IDH1 REV | Tsingke | AGGCCAACCCTTAGACAGAGC |
| IDH2 FW | Tsingke | AGTGCGTTTTACCTCAGCCAGT |
| IDH2 REV | Tsingke | GTCCCACTACCTCCTCCCCTC |
| ME1 FW | Tsingke | GGGTCGGCTTTATCCTCCTT |
| ME1 REV | Tsingke | ATGCTTCTTTGTTTTGCGGTT |
| PGD FW | Tsingke | GTGGCCCCACATCAAGACC |
| PGD REV | Tsingke | GTCCCCATACTCTATCCCGTT |
| TIGAR FW | Tsingke | TCACTGCCCTAAAAGTCCTCT |
| TIGAR REV | Tsingke | AAACTTCTGCCCATTCCCTC |
| FTH1 FW | Tsingke | CTACTGGAACTGCACAAACTGG |
| FTH1 REV | Tsingke | CACCCAATTCTTTGATGGCTT |
| FTL FW | Tsingke | TGAGCCACTTCTTCCGCGAAT |
| FTL REV | Tsingke | CGCCACGCTGGTTTTGCATC |
| HO1 FW | Tsingke | CAGTGCCACCAAGTTCAAGCA |
| HO1 REV | Tsingke | CAGCTCCTGCAACTCCTCA |
| SESN1 FW | Tsingke | CCTCAATGCTTAGACGGGCAA |
| SESN1 REV | Tsingke | TTCAGGAGTGCAAACAACAGT |
| SESN2 FW | Tsingke | CACCCAGACATGCTGTGCTT |

| | | |
|---|---|---|
| SESN2 REV | Tsingke | TAGCCATGGTCTTCCCAGGT |
| SESN3 FW | Tsingke | AGAACTTGTGCCATATCTCGT |
| SESN3 REV | Tsingke | CCATTTGTGTTTTGCCATTCTCC |
| NDUFS1 FW | Tsingke | TTCCAGCAAGCAAATGAGCTCT |
| NDUFS1 REV | Tsingke | AGGCTCTGCTAATTGAATCTGT |
| NDUFV1 FW | Tsingke | GAGAACGCAACTCAGGCACCAA |
| NDUFV1 REV | Tsingke | CTTTCAAGGGCACAGACATCTCC |
| NDUFV2 FW | Tsingke | TCAGGTCTGCACTACTACACCC |
| NDUFV2 REV | Tsingke | CTCCCCAACCTTTATTCCAAGC |
| ND1 FW | Tsingke | ATTACTCCTGCCATCATGACCC |
| ND1 REV | Tsingke | TTCGGACTCCCCTTCGGCAAG |
| NDUFC2 FW | Tsingke | ACTCAGGAGCTAATACTGTCT |
| NDUFC2 REV | Tsingke | GGACAAGCATTTGATAAGCCAT |
| SDHA FW | Tsingke | CCTACTTCAGCTGCACGTCT |
| SDHA REV | Tsingke | ACAACCAGCACCATATATGCC |
| SDHB FW | Tsingke | ACCTTCCGAAGATCATGCAG |
| SDHB REV | Tsingke | CAATCCTTCGGGTGCAAGCTA |
| SDHC FW | Tsingke | CCATATAGGCCAGGCTTTAGCA |
| SDHC REV | Tsingke | GGCCAACAACTCGAAGTGACA |
| SDHD FW | Tsingke | CCTCTTTGCCTCTGCTTTGTCA |
| SDHD REV | Tsingke | AGAAGAAGGCTGTCCACCAAT |
| UQCRFS1 FW | Tsingke | TCCCTACTGTGTAAGCCGTGT |
| UQCRFS1 REV | Tsingke | ACCAGACGTGAAGATAAACCCT |
| CYC1 FW | Tsingke | CTCGAGCTGCCAACAACGGA |
| CYC1 REV | Tsingke | TAGAGACCTTCCCGCAGTGAC |
| MTCYB FW | Tsingke | CAGTCCCACCCTCACACGAT |

| | | |
|---|---|---|
| MTCYB REV | Tsingke | GGTTGTTTGATCCCGTTTCGT |
| COX5A FW | Tsingke | ACAAAGCAGGACCTCATAAGGA |
| COX5A REV | Tsingke | CCATGCGGTTTACACTTTGTCA |
| COX6A FW | Tsingke | AGACCCGAGTTCATCGCCTA |
| COX6A REV | Tsingke | TCATCTTCGTAGCCAGTTGGA |
| COX7A FW | Tsingke | AGGAGGACAATGACATCCCGTT |
| COX7A REV | Tsingke | ACACAGCGTCATTGTCACTCG |
| COX8A FW | Tsingke | CGCGCCAAGATCCATTCGTTG |
| COX8A REV | Tsingke | CACTCTGGCCTCCTGTAGGTC |
| MTCO1 FW | Tsingke | ACCCCATTCTATACCAACACCT |
| MTCO1 REV | Tsingke | ATGGGAGATTATTCCGAAGCCT |
| MTCO2 FW | Tsingke | GCTGTCCCCACATTAGGCTT |
| MTCO2 REV | Tsingke | AGATTTCAGAGCATTGACCGTA |
| OPA1 FW | Tsingke | ACATCAACGTCTTTTGTCCAGC |
| OPA1 REV | Tsingke | ACTGTTCTTCCATTCGAGAGGC |
| MFN1 FW | Tsingke | TACCAACTTATGTGACCCCTG |
| MFN1 REV | Tsingke | AAAGGCTCAATCTCTAACTCC |
| MFN2 FW | Tsingke | CGCATCTTCTTTGTGTCTGCT |
| MFN2 REV | Tsingke | GATGCACTCCTCAAATCTCCT |
| DRP1 FW | Tsingke | ATTCCAATTATGCCAGCCAGT |
| DRP1 REV | Tsingke | TCCCGAGCAGATAGTTTTCGT |
| MTRF1 FW | Tsingke | CACCCCTTCAGGATGACCTT |
| MTRF1 REV | Tsingke | CGCAAATCTTCTGCAGTGCTT |
| PGC1α FW | Tsingke | AGACCTGACACAACACGGACA |
| PGC1α REV | Tsingke | CAAGAGCAGCAAAAGCATCACA |
| PGC1β FW | Tsingke | ACAGGCTAATGTAATTTGTTGCT |

| | | |
|---|---|---|
| PGC1β REV | Tsingke | CATGGGCTCATTTTACCTACTGA |
| αPal$^{NRF1}$ FW | Tsingke | TGACCATCCAGACGACGCAAG |
| αPal$^{NRF1}$ REV | Tsingke | CGCCATAGTGACTGTAGCTCC |
| GABPα FW | Tsingke | TAAATCAGCCTGAACTGGTTGC |
| GABPα REV | Tsingke | TTACAAATCATGTCCCCATCGT |
| TFAM FW | Tsingke | CAAGTGATCCTTCCGAGTCGTT |
| TFAM REV | Tsingke | GGGCAACATAGCAAGATCCCAT |
| TFB1M FW | Tsingke | AGATAGAGCAGCCATTCAAGC |
| TFB1M REV | Tsingke | GTTCTTCTCTGAGCCCTCGAT |
| TFB2M FW | Tsingke | ACCACCTGCTATGTCTTCTCG |
| TFB2M REV | Tsingke | AAGTGCCCTTTTCTCACCT |
| Nrf1 FW | Tsingke | AACATTCTGGTCCTTCAGCAATGCT |
| Nrf1 REV | Tsingke | ACCCGTACCCCAATCAAACTCAGCA |
| Nrf2 FW | Tsingke | GCCCACATTCCCAAATCAGATGC |
| Nrf2 REV | Tsingke | ACAAGTGACTGAAACGTAGCCGAA |
| HK1 FW | Tsingke | CACATGGAGTCCGAGGTTTATG |
| HK1 REV | Tsingke | CGTGAATCCCACAGGTAACTTC |
| HK2 FW | Tsingke | GAGCCACCACTCACCCTACT |
| HK2 REV | Tsingke | CCAGGCATTCGGCAATGTG |
| GLUT1 FW | Tsingke | TCCAGCTGCCATTGCCGTTG |
| GLUT1 REV | Tsingke | AGGGACCACACAGTTGCTCCAC |
| GLUT4 FW | Tsingke | TGACCAGATCTCAGCTGCCTT |
| GLUT4 REV | Tsingke | CCTGGCCCCTCAGTCGTTC |
| LDHA FW | Tsingke | ATGGCAACTCTAAAGGATCAGC |
| LDHA REV | Tsingke | CCAACCCCAACAACTGTAATCT |
| PDH FW | Tsingke | AGCGCCTCTCTAAAATGCTT |

| | | |
|---|---|---|
| PDH REV | Tsingke | AGCCACTCACTATTCTTGGTT |
| PDK1 FW | Tsingke | ATTCAACTGCACCAAGACCT |
| PDK1 REV | Tsingke | TGCATCTGTCCCGTAACCCTC |
| SREBP1 FW | Tsingke | GGAGCCATGGATTGCACTTT |
| SREBP1 REV | Tsingke | CAGGAAGGCTTCAAGAGAGG |
| SREBP2 FW | Tsingke | CGACTCTGACCAGCACCCACA |
| SREBP2 REV | Tsingke | GACGCTCAGGACAATCACACC |
| ACCα FW | Tsingke | GTTATTTCTCAGAGCTTCCGAAC |
| ACCα REV | Tsingke | GAATTTCTTCTGCCAGTCCGAT |
| ACLY FW | Tsingke | ATCGGTTCAAGTATGCTCGGG |
| ACLY REV | Tsingke | GACCAAGTTTTCCACGACGTT |
| FASN FW | Tsingke | GCCTGCCACAACTCCAAGGAC |
| FASN REV | Tsingke | TTCCTCAGCTGCTCCACGAAC |
| SCD1 FW | Tsingke | CACCACATTCTTCATTGATTGC |
| SCD1 REV | Tsingke | TCAGCCACTCTTGTAGTTTCCA |
| CPT1A FW | Tsingke | TGCAAAGGCGACATCAATCCG |
| CPT1A REV | Tsingke | AAATCCACGTCGTTTGCCAGA |
| UCP2 FW | Tsingke | GCTTTGCCTCTGTCCGCATC |
| UCP2 REV | Tsingke | CATAGGTCACCAGCTCAGCAC |
| ATF4 FW | Tsingke | CCCTTCACCTTCTTACAACCTC |
| ATF4 REV | Tsingke | TGCCCAGCTCTAAACTAAAGGA |
| ATF5 FW | Tsingke | GCCTCTACCTTCATTCCAAACCC |
| ATF5 REV | Tsingke | TTGACCTCCTGGCCTCAAACGA |
| CHOP FW | Tsingke | GAAACAGAGTGGTCATTCCCC |
| CHOP REV | Tsingke | GATTTCCTGCTTGAGCCGTTC |
| HSP60 FW | Tsingke | GCCCTCCTTCGATGCATTCCA |

| | | |
|---|---|---|
| HSP60 REV | Tsingke | ACCTGCATTCTTAGCAATGGTCA |
| GRP75 FW | Tsingke | AGCAGCAGATTGTAATCCAGT |
| GRP75 REV | Tsingke | TCGTTCCTTCTTTCGCCGGTCT |
| FGF21 FW | Tsingke | GGAGCTTCTGCATCTATCCCAA |
| FGF21 REV | Tsingke | GTGCCAGATTCCAGTTGTCCA |
| ACTB FW | Tsingke | TGGCATCCACGAAACTACCTT |
| ACTB REV | Tsingke | CTTCTGCATCCTGTCGGCAAT |
| **Oligonucleotides for construct** | | |
| UCP2 FW | Tsingke | GGGGTACCATGGTTGGGTTCAAGGCCACAGATG |
| UCP2 REV | Tsingke | GCTCTAGACTGAAGGGAGCCTCTCGGGAAGTGCAG |
| PGC1α-luc FW | Tsingke | CTAGCTAGCCTTCAGACTTCTGAATACTGCACAG |
| PGC1α-luc REV | Tsingke | CCCAAGCTTAATGGCAAAGGGAACGGAAACTCTC |
| αPal$^{NRF1}$-luc FW | Tsingke | CTAGCTAGCGCGTATTCTTCCACGTAGGCTAAGC |
| αPal$^{NRF1}$-luc REV | Tsingke | CCGCTCGAGGTTTCCTCGGACTATGCCTCACAAC |
| GABPα-luc FW | Tsingke | CTAGCTAGCCCAGCCTGCCCAAAATGGTGAAAC |
| GABPα-luc REV | Tsingke | CCGCTCGAGATGCGTACAGAGCAAGGTTTCAAG |
| UCP2-luc FW | Tsingke | CTAGCTAGCAATTCCATCTGGCTTTGTGTCACCA |
| UCP2-luc REV | Tsingke | CCGCTCGAGTGGGAGAGAAGGTAAATGGAAGAGC |
| miR-195-luc FW | Tsingke | CTAGCTAGCAGTCTTGTGGACACTCACGTTCCAC |
| miR-195-luc REV | Tsingke | CCGCTCGAGGTGCCAATATTTCTGTGCTGCTAGA |
| miR-497-luc FW | Tsingke | CTAGCTAGCGAAGCTAAGGGATAATCATAAGGTGAG |
| miR-497-luc REV | Tsingke | CCGCTCGAGCCGTACAAACCACAGTGTGCTGCTG |
| 3 -UTRwt of UCP2 FW | Tsingke | CGCTCGAGTGCCCTCCTTTCTCCGCTTGGGTT |
| 3 -UTRwt of UCP2 REV | Tsingke | ATAAGAATGCGGCCGCATTCTGGCTGAACTTTCCAAGGGAC |
| **Software and Algorithms** | | |
| Canvas X | Canvas GFX, Inc. | https://www.canvasgfx.com/ |

| | | |
|---|---|---|
| FlowJo 7.6.5 | FlowJo | https://www.flowjo.com/ |
| Primer Premier 5 | PREMIER Biosoft International | https://www.PremierBiosoft.com/ |
| CFX Manager 3.1 | Bio-Rad | https://bio-rad-cfx-manager.com/ |
| Quantity One | Bio-Rad | https://bio-rad-cfx-manager.com/ |
